# *De novo* protein design enables precise induction of functional antibodies *in vivo*

**DOI:** 10.1101/685867

**Authors:** Fabian Sesterhenn, Che Yang, Jaume Bonet, Johannes T Cramer, Xiaolin Wen, Yimeng Wang, Chi-I Chiang, Luciano A Abriata, Iga Kucharska, Giacomo Castoro, Sabrina S Vollers, Marie Galloux, Elie Dheilly, Stéphane Rosset, Patricia Corthésy, Sandrine Georgeon, Mélanie Villard, Charles-Adrien Richard, Delphyne Descamps, Teresa Delgado, Elisa Oricchio, Marie-Anne Rameix-Welti, Vicente Más, Sean Ervin, Jean-François Eléouët, Sabine Riffault, John T Bates, Jean-Phillipe Julien, Yuxing Li, Theodore Jardetzky, Thomas Krey, Bruno E Correia

## Abstract

*De novo* protein design has been successful in expanding the natural protein repertoire. However, most *de novo* proteins lack biological function, presenting a major methodological challenge. In vaccinology, the induction of precise antibody responses remains a cornerstone for next-generation vaccines. Here, we present a novel protein design algorithm, termed TopoBuilder, with which we engineered epitope-focused immunogens displaying complex structural motifs. Both in mice and non-human primates, cocktails of three *de novo* designed immunogens induced robust neutralizing responses against the respiratory syncytial virus. Furthermore, the immunogens refocused pre-existing antibody responses towards defined neutralization epitopes. Overall, our *de novo* design approach opens the possibility of targeting specific epitopes for vaccine and therapeutic antibody development, and more generally will be applicable to design *de novo* proteins displaying complex functional motifs.

## Introduction

The computational design of novel proteins from first principles has revealed a variety of rules for the accurate design of structural features in *de novo* proteins (*1–4*). However, the *de novo* design of functional proteins remains far more challenging (*5, 6*). A commonly used strategy to design functional proteins is to transplant structural motifs from other proteins to pre-existing or *de novo* protein scaffolds (*7–10*). In nearly all cases previously reported, the motifs transplanted mediated protein-protein interactions. These structural motifs were common in the natural protein repertoire, such as linear helical segments, allowing their grafting without extensive backbone adjustments (*7–9*). Most protein functional sites, however, are not contained within regular single segments in protein structures, but arise from the three-dimensional arrangement of multiple and often irregular, structural elements supported by the overall architecture of the protein structure (*11–13*). Additionally, many functional sites involved in molecular recognition events form protein clefts, which mediate the interaction with its cognate partners but have proven very challenging to encode in designed proteins (*2*). Thus, the development of computational approaches to endow *de novo* proteins with irregular and multi-segment motifs is crucial to expand their function and scope of applications.

Protein design has sparked hopes in the field of rational vaccinology, in particular, to elicit targeted neutralizing antibody (nAb) responses (*10, 14*). Although many potent nAbs have been identified and structurally characterized in complex with their target antigens, the design of immunogens that elicit precise and focused antibody responses remains a major challenge (*15, 16*). To date, structure-based immunogen design efforts have mostly focused on modifying viral fusion proteins, through conformational stabilization to present neutralization-sensitive epitopes, silencing of non-neutralizing epitopes through glycosylation or domain deletions and germline-targeting approaches (*17*). Unlike RSV, several major human pathogens only display a limited number of broadly neutralizing epitopes, surrounded by strain-specific, non-neutralizing, or in some cases, disease-enhancing epitopes (*18–20*). Thus, one of the central goals for vaccine development is to elicit antibody responses with precisely defined epitope specificities, and, in some cases, constrained molecular features (e.g. antibody lineage, CDR length, binding angle) (*21–25*).

The difficulty in developing immunogens that empower elicitation of antibodies specific for a restricted subset of epitopes on a single protein, and consequently the fine specificity of the B cell response following immunization continues to be a critical barrier to rational vaccine design. Towards this goal, previous studies have sought to elicit epitope-specific responses using peptide-based approaches (*26*) or computationally designed epitope-scaffolds (*10, 14, 27-29*). Leveraging computational design, the RSVF antigenic site II, a linear helix-turn-helix motif, was transplanted onto a heterologous protein scaffold, which was shown to elicit nAbs in non-human-primates (NHPs) after repeated boosting immunizations (*10*). Despite the proof-of-principle for the induction of functional antibodies using a computationally designed immunogen, several major caveats emerged: on the vaccine side, the neutralization titer observed in the immunogenicity studies were inconsistent; and technically, the computational approach was not suitable for structurally complex epitopes.

To address these limitations, here we used *de novo* design approaches, including a newly developed method, to engineer epitope-focused immunogens mimicking irregular and discontinuous RSV neutralization epitopes (site 0 (*30*) and IV (*31*), Fig 1). We designed a trivalent cocktail formulation (“Trivax”) consisting of a previously published immunogen for site II (*14*), and the newly designed immunogens mimicking sites 0 and IV. *In vivo*, Trivax induced a balanced antibody response against all three epitopes, resulting in consistent levels of serum neutralization in 6/7 NHPs. When used as boosting immunogens upon priming with a viral antigen, the computationally designed immunogens were functional to reshape the serum composition towards higher levels of site-specific antibodies, resulting in an improved antibody quality. Our approach enables the targeting of specific epitopes for vaccine and therapeutic antibody development and more broadly will be applicable to design *de novo* proteins displaying complex functional motifs.

**Fig 1.**
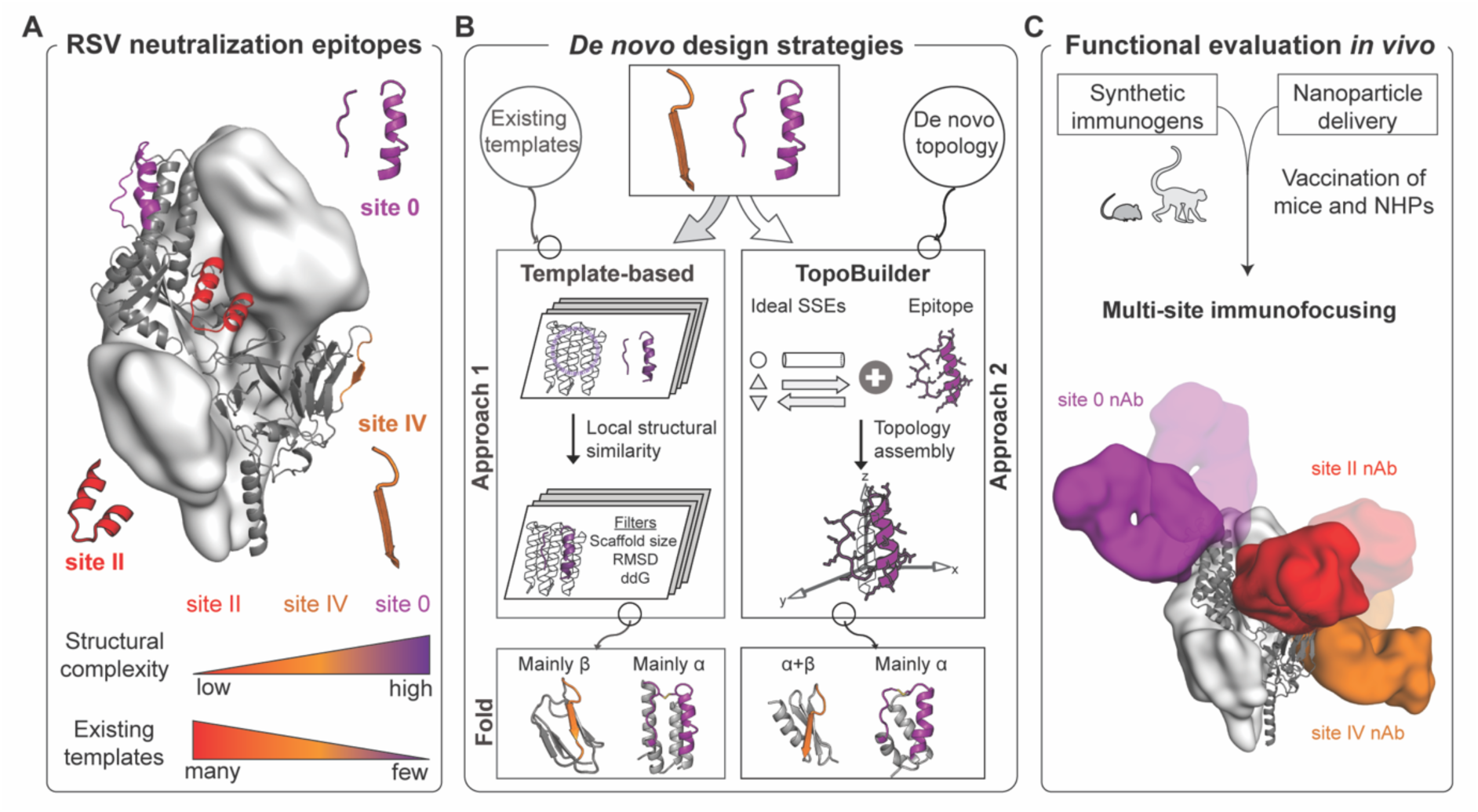
Computational design of epitope-focused immunogens to elicit RSV nAbs targeting three distinct epitopes. (A) Prefusion RSVF structure (PDB 4JHW) with sites 0, II and IV highlighted. An immunogen for site II was previously reported (*10, 14*). (B) Computational protein design strategies. Approach 1: Design templates were identified in the PDB based on loose structural similarity to site 0 and IV, followed by *in silico* folding and design, and sequence optimization through directed evolution. Approach 2: A motif-centric *de novo* design approach was developed (“TopoBuilder”) to tailor the protein topology to the motif’s structural constraints. Bottom: Computational models of designed immunogens using different approaches. (C) Cocktail formulations of three synthetic immunogen nanoparticles elicit nAbs focused on three non-overlapping epitopes. SSEs - secondary structure elements; α - alpha-helix; β - beta-strand; ddG - computed binding energy.

## Results

### *De novo* design of immunogens presenting structurally complex epitopes

The computational design of proteins mimicking structural motifs has been performed previously by first identifying compatible protein scaffolds which then serve as design templates on which to graft the motifs (*7, 27-29, 32, 33*). This approach, referred as template-based design, has been used to transplant functional sites to both structures from the natural repertoire (*27-29, 32, 33*) or *de novo* designed proteins (*7*). While most studies focused on linear, regular binding motifs, Azoitei et al. successfully grafted a structurally complex HIV epitope into an existing protein scaffold, but both the overall structure and sequence of the template remained mostly native (*32*).

Here, we sought to design accurate mimetics of RSVF neutralization epitopes based on *de novo* proteins and evaluate their functionality in immunization studies. We chose antigenic sites 0 and IV (Fig 1a), which are both targeted by potent nAbs, and have a high structural complexity: antigenic site 0 presents a structurally complex and discontinuous epitope consisting of a kinked 17-residue alpha helix and a disordered loop of 7 residues (*30, 34*), while site IV presents an irregular 6-residue bulged beta-strand (*31*).

In a first effort we used a template-based *de novo* design approach relying on Rosetta FunFolDes (*35*) to fold and design novel proteins to present sites IV and 0. Given the structural complexity of these sites, few structures in the Protein Data Bank (PDB) matched the backbone configuration of the epitopes, even using loose structural criteria (Fig S1). Briefly, our best computational design for site IV (S4_1.1), based on a domain excised from prefusion RSVF (preRSVF), bound with weak affinity to the target nAb 101F (K_D_ > 85 µM). After *in vitro* evolution, we obtained a double mutant (S4_1.5) that bound 101F with a K_D_ of 35 nM and was thermostable up to 65 °C (Fig 2b-d and Fig S2-3). For site 0, we used a designed helical repeat protein (PDB 5CWJ (*36*)) as design template. Our first computational design showed a K_D_ of 1.4 µM to the target D25 nAb, which we improved to a K_D_ of 5 nM upon several truncations, and iterative rounds of computational design and *in vitro* evolution (Fig 2 and Fig S4-5).

**Fig 2.**
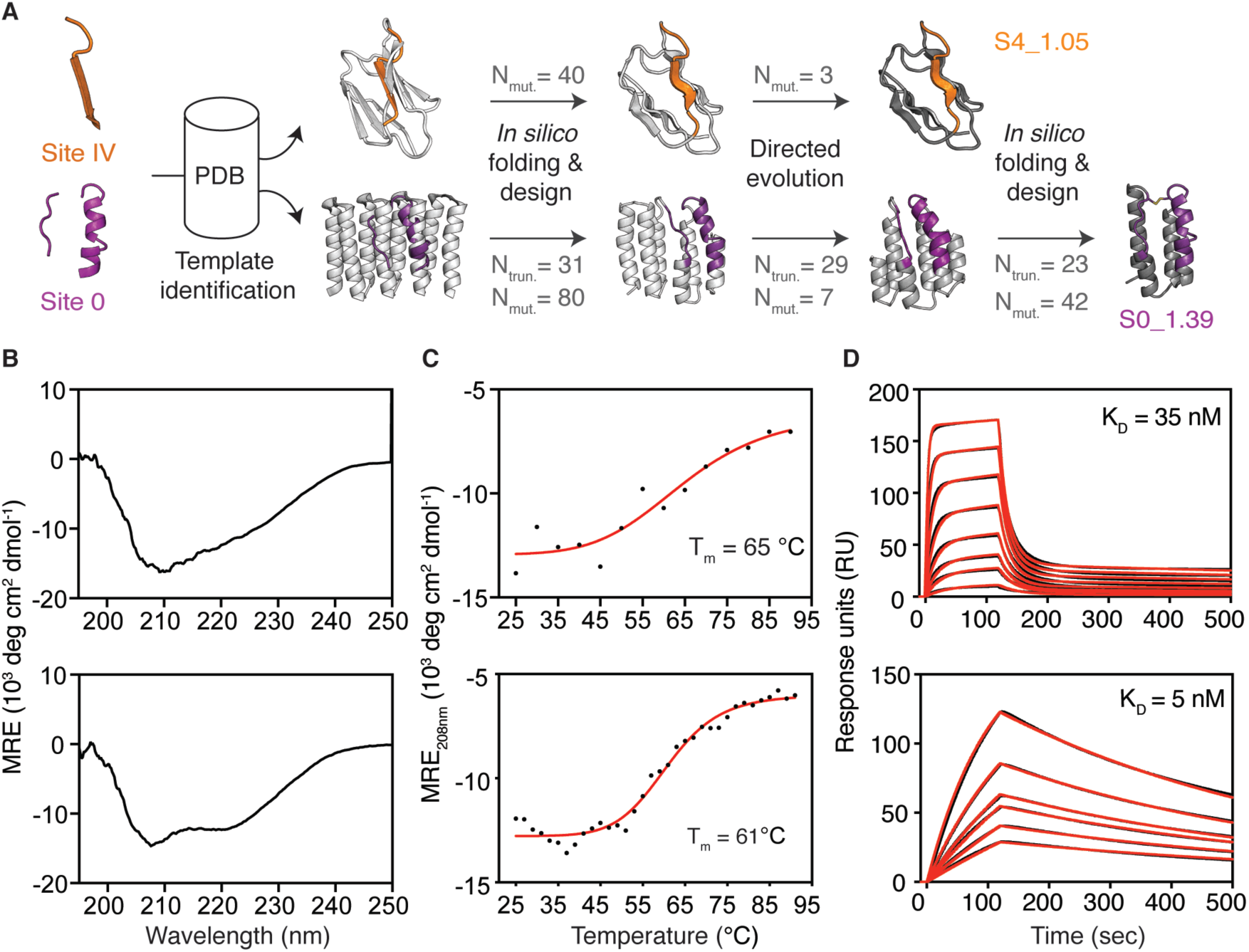
Templated computational design and biophysical characterization of synthetic immunogens. (A) Protein design strategy. Templates with structural similarity to sites IV and 0 were identified by native domain excision or loose structural matching, followed by *in silico* folding, design and directed evolution. An additional *in silico* folding and design step was necessary to install site 0 on a truncated template sequence revealed by directed evolution. Computational models of intermediates and final designs (S4_1.5 and S0_1.39) are shown, and the number of mutations (Nmut) and truncated residues (Ntrun) are indicated for each step. (B) CD spectra measured at 20 °C of S4_1.5 (top) and S0_1.39 (bottom), are in agreement with the expected secondary structure content of the design model. (C) Thermal melting curves measured by CD in presence of 5 mM TCEP reducing agent. (D) Binding affinity measured by SPR against target antibodies 101F (top) and D25 (bottom). Sensorgrams are shown in black and fits in red. CD - circular dichroism, Tm - melting temperature, SPR - surface plasmon resonance.

The template-based approach led to designs that presented several desired features (e.g. stability and antibody binding), however important limitations emerged during the design process: (1) extensive *in vitro* evolution optimization was required; (2) binding affinities to target nAbs were one to two orders of magnitude lower than those of the viral protein (preRSVF); (3) suboptimal template topologies constrained the epitope accessibility (see details in Fig S6).

In order to address these limitations, we developed a template-free design protocol - TopoBuilder - that generates tailor-made protein topologies to stabilize complex functional motifs. The TopoBuilder consists of three stages (Fig 3a): (1) Topological sampling in 2D space - to quickly define the fold space compatible with the target structural motif we use the αβα-Form topology definition scheme, a string-based descriptor that allows the extensive enumeration of multilayer protein topologies with alternating secondary structure elements and all possible connections between the secondary structural elements (*37, 38*). Putative folds are then selected according to basic topological rules (e.g. lack of crossover loops and chain directionality of the functional motif). Together, this allowed the definition of the fold space for a given design task, thereby overcoming a main hurdle in *de novo* design approaches. (2) 3D projection and parametric sampling - the selected 2D topologies are projected into the 3D space by assembling idealized secondary structures (SSE) around the fixed functional motif. These 3D structures, referred to as ‘sketches’, are further refined by coarsely sampling structural features of the fold (e.g. distances and orientations between SSEs) using parametric sampling. (3) Flexible backbone sampling and sequence design - to refine the structural features of the sketches at the all-atom level and design sequences that stabilize these structures, we use Rosetta FunFolDes as described before (*10, 35*). Here, we leverage all the functionalities of FunFolDes in terms of coupling protein folding and sequence design to stabilize the motifs of interest within the context of the *de novo* topology, a conceptually different approach of grafting structural motifs into largely static scaffolds (*7, 8, 27, 28, 32*).

**Fig 3.**
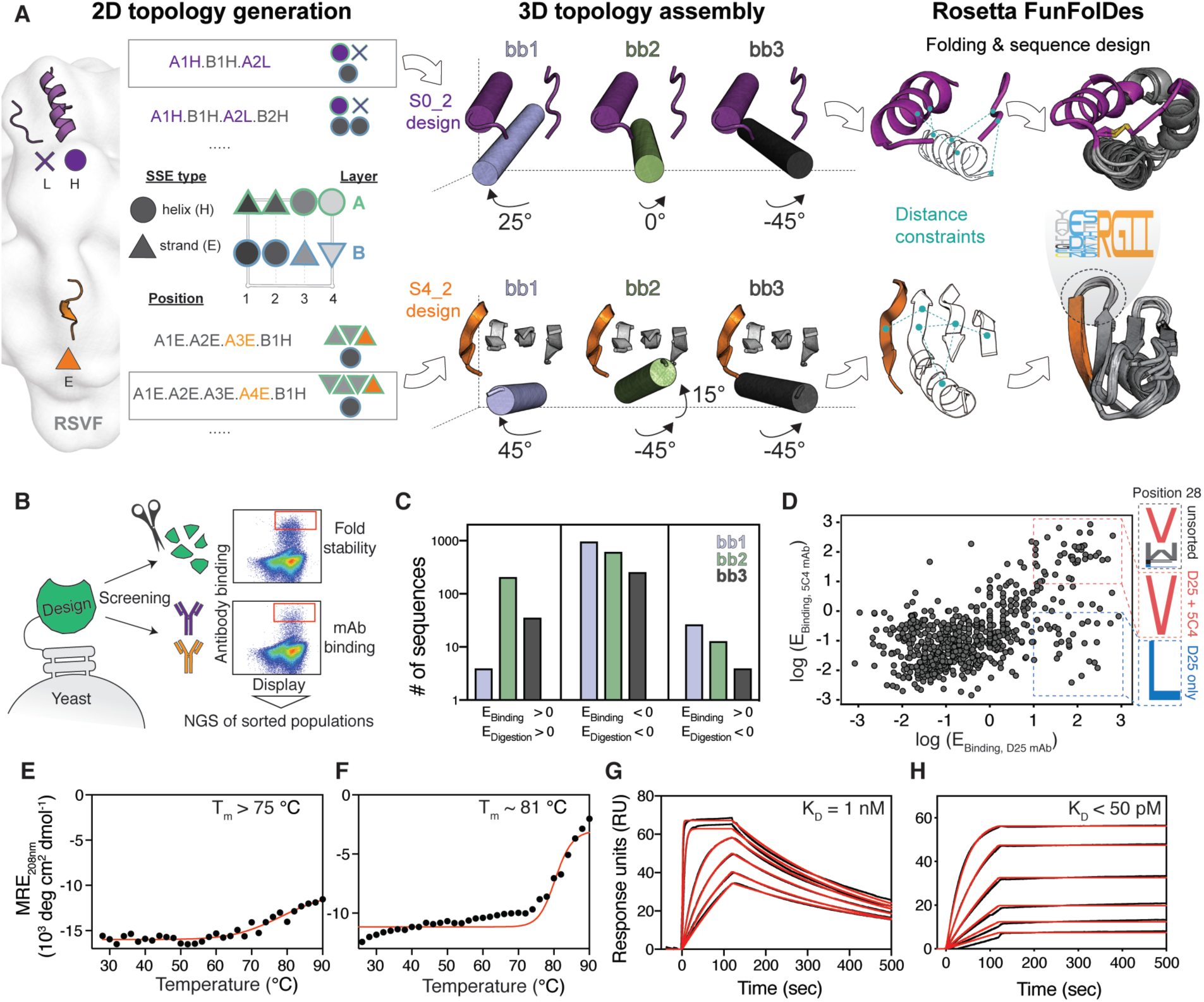
Template-free *de novo* design of epitope-focused immunogens. (A) Protein topologies that are compatible with each motif are enumerated in the 2D space. Selected topologies are then projected into the 3D space using idealized SSEs, and their relative orientation is sampled parametrically. Distance constraints are derived from selected topologies to guide *in silico* folding and sequence design using Rosetta. (B) Designed sequences were screened for high affinity binding and resistance to chymotrypsin to select stably folded proteins, as revealed by next-generation sequencing (NGS). (C) For the S4_2 design series, enrichment analysis revealed a strong preference for one of the designed helical orientations (S4_2_bb2, green) to resist protease digestion and to bind with high affinity to 101F. (D) To ensure epitope integrity, S0_2_bb3 was screened for binding to both D25 and 5C4. Sequences highly enriched for both D25 and 5C4 binding show convergent sequence features in the critical core position 28 of the site 0 scaffold. (E-F) Thermal melting curves measured by CD for best designs (S4_2.45 (E) and S0_2.126 (F)) showing high thermostability. (G-H) Dissociation constants (K_D_) of S4_2.45 to 101F (G) and S0_2.126 to D25 (H) as measured by SPR. E - enrichment.

To present antigenic site IV, we designed a fold composed of a β-sheet with 4 antiparallel strands and one helix (Fig 3a), referred to as S4_2 fold. Within the S4_2 topology, we generated three structural variants (S4_2_bb1, S4_2_bb2 and S4_2_bb3), by sampling three distinct orientations of the helical element, varying both orientations and lengths to optimize the packing interactions with the β-sheet. Sequences generated from two structural variants (S4_2_bb2 and S4_2_bb3) showed a strong propensity to recover the designed structures in Rosetta *abinitio* simulations (Fig S7).

To evaluate our design approach, we screened a library of designed sequences using yeast display and applied two selective pressures – binding to the 101F antibody and resistance to the nonspecific protease chymotrypsin (Fig 3b), an effective method to digest partially unfolded proteins (*7, 39, 40*). To reveal structural and sequence determinants of designs that led to stable folds and high-affinity binding to 101F, we performed next-generation sequencing of populations sorted under different conditions, retrieving stability and binding scores for each design. We found that S4_2_bb2-based designs were preferentially enriched over the bb1 and bb3 design series, showing that subtle topological differences in the design template can have substantial impact on function and stability (Fig 3c). 13 out of the 14 best-scoring S4_2 variants, bearing between 1 and 38 mutations compared to each other, were successfully purified and biochemically characterized (Fig S8). The designs showed mixed alpha/beta CD spectra and bound to 101F with affinities ranging from 1 nM to 200 nM (Fig S8). The best variant, S4_2.45 was well folded according to CD and NMR, and only showed partial unfolding even at 90 °C (Fig 3e and Fig S9). S4_2.45 showed a K_D_ = 1 nM to the target antibody 101F (Fig 3g), in line with the preRSVF-101F interaction (K_D_ = 4 nM).

Similarly, we built a minimal *de novo* topology to present the tertiary structure of the site 0 epitope. The choice for this topology was motivated by the native environment of site 0 in preRSVF, where it is accessible for antibodies with diverse angles of approach (*34*) (Fig S6). We explored the topological space within the shape constraints of preRSVF and built three different helical orientations (S0_2_bb1-3) that supported the epitope segments. Rosetta *abinitio* folding predictions showed that only designs based on one topology (S0_2_bb3) presented funnel-shaped energy landscapes (Fig S10). A set of computationally designed sequences based on the S0_2_bb3 template was screened in yeast under the selective pressure of two site 0-specific antibodies (D25 and 5C4), to ensure the presentation of the native epitope conformation. Deep sequencing of the double-enriched clones revealed that subtle sequence variants (e. g. position 28) are sufficient to change the antibody binding properties of the designs, highlighting the challenges of designing functional proteins (Fig 3d). From the high-throughput screening, we selected five sequences, bearing between 3 and 21 mutations compared to each other in a protein of 58 residues, for further biochemical characterization (Fig S11). The design with best solution behaviour (S0_2.126) showed a CD spectrum of a predominantly helical protein, with extremely high thermostability even under reducing conditions (T_m_ = 81 °C, Fig 3f) and a well-dispersed HSQC NMR spectrum (Fig S9). Strikingly, S0_2.126 bound with K_D_s of ∼50 pM and 4 nM to D25 and 5C4, respectively, which is in line with the affinities of the nAbs to preRSVF (∼150 pM and 13 nM for D25 and 5C4, respectively) (Fig 3h and Fig S12).

To investigate whether the template-free design approach together with the screening procedure yielded scaffolds that better mimicked the viral epitopes, we determined the affinities of S4_2.45 and S0_2.126 against a panel of site-specific human nAbs (*41*). Compared to the first-generation designs, S4_2.45 and S0_2.126 showed large affinity improvements across the antibody panels, exhibiting a geometric mean affinity closely resembling that of the antibodies to preRSVF (Fig S12). These results suggest that the immunogens designed using TopoBuilder were superior mimetics of sites IV and 0 as compared to the template-based designs.

### *De novo* designed topologies adopt the predicted structures with high accuracy

To evaluate the structural accuracy of the computational design approach, we solved the crystal structure of S4_2.45 in complex with 101F at 2.6 Å resolution. The structure closely resembled our design model, with a backbone RMSD of 1.5 Å (Fig 4a). The epitope was mimicked with an RMSD of 0.23 Å, and retained all essential interactions with 101F (Fig 4d, e and Fig S13). Together, the structural data confirmed that we accurately presented an irregular beta strand, a common motif found in many protein-protein interactions (*42*), in a fully *de novo* designed protein with sub-angstrom accuracy.

**Fig 4.**
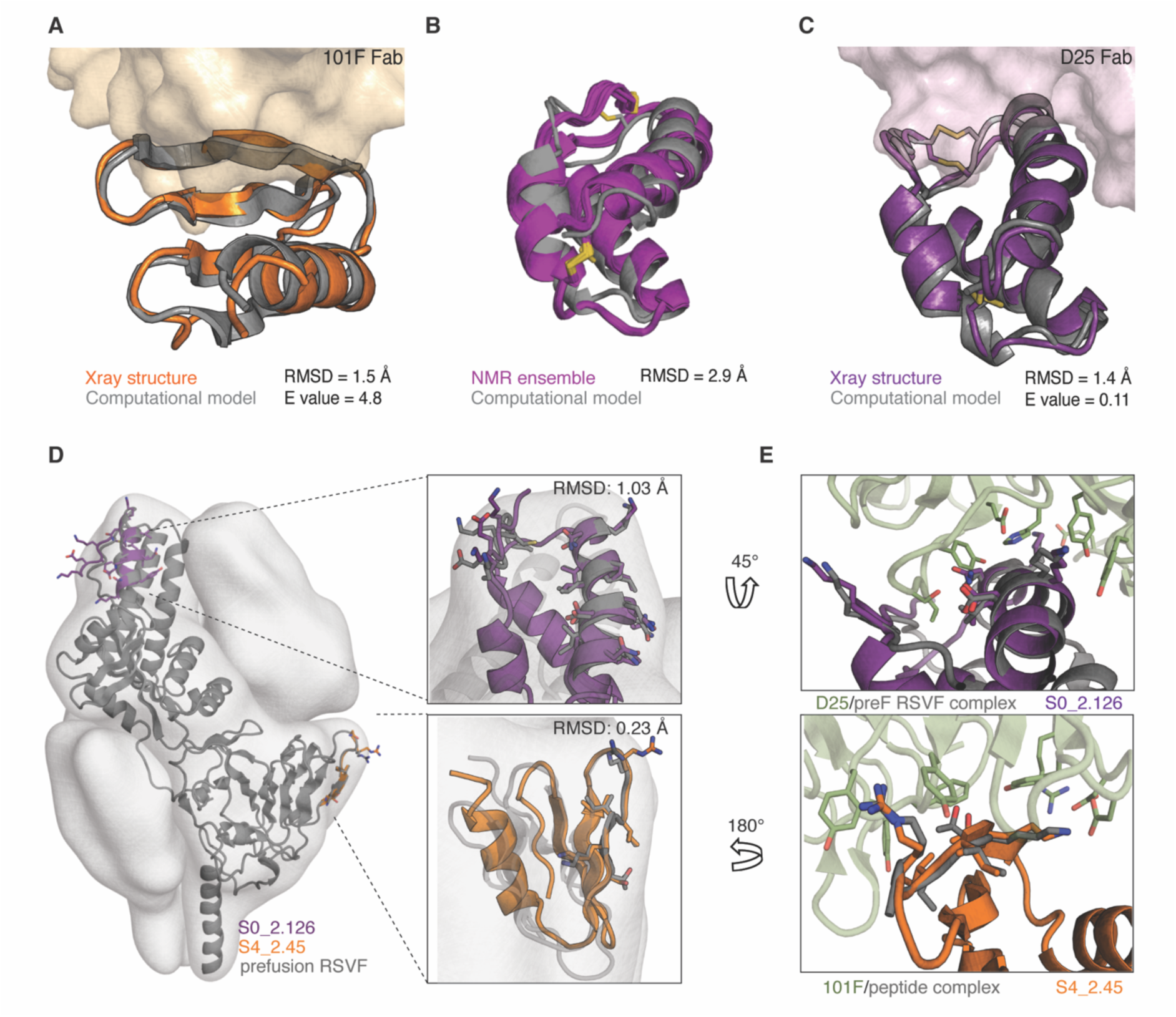
Structural characterization of *de novo* designed immunogens. (A) Crystal structure of S4_2.45 (orange) bound to 101F Fab closely matches the design model (grey, RMSD = 1.5 Å). (B) NMR structural ensemble of S0_2.126 (purple) superimposed to the computational model (grey). The NMR structure shows overall agreement with the design model (backbone RMSD of 2.9 Å). (C) Crystal structure of S0_2.126 (purple) bound to D25 Fab closely resembles the design model (grey, RMSD = 1.4 Å). (D) Superposition of the preRSVF sites 0/IV and designed immunogens shows atomic-level mimicry of the epitopes. Designed scaffolds are compatible with the shape constraints of preRSVF (surface representation). (E) Close-up view of the interfacial side-chain interactions between D25 (top) and 101F (bottom) with designed immunogens as compared to the starting epitope structures.

Next, we solved an unbound structure of S0_2.126 by NMR, confirming the accuracy of the designed fold with a backbone RMSD of 2.9 Å between the average structure and the computational model (Fig 4b). Additionally, we solved a crystal structure of S0_2.126 bound to D25 at a resolution of 3.0 Å. The structure showed backbone RMSDs of 1.4 Å to the design model and 1.03 Å over the discontinuous epitope compared to preRSVF (Fig 4c-e and Fig S13). In comparison with native proteins, S0_2.126 showed exceptionally low core packing due to a large cavity (Fig S14), but retained a very high thermal stability. The core cavity was essential for antibody binding and highlights the potential of *de novo* approaches to design small proteins hosting structurally challenging motifs and preserving cavities required for function (*2*). Notably, due to the level of control and precision of the TopoBuilder, both designed antigens respected the shape constraints of the epitopes in their native environment within preRSVF, a structural feature that may be important for the induction of functional antibodies (Fig S6).

### Cocktails of designed immunogens elicit nAbs *in vivo* and reshape pre-existing immunity

Lastly, we evaluated the ability of the designed antigens to elicit targeted nAb responses *in vivo*. Our rationale for combining site 0, II and IV immunogens in a cocktail formulation is that all three sites are non-overlapping in the preRSVF structure (Fig S15), and thus might induce a more potent and consistent nAb response *in vivo*. To increase immunogenicity, each immunogen was multimerized on self-assembling protein nanoparticles. We chose the RSV nucleoprotein (RSVN), a self-assembling ring-like structure of 10-11 subunits, previously shown to be an effective carrier for the site II immunogen (S2_1.2) (*14*), and formulated a trivalent immunogen cocktail containing equimolar amounts of S0_1.39, S4_1.5 and S2_1.2 immunogen nanoparticles (“Trivax1”, Fig S16). The fusion of S0_2.126 and S4_2.45 to RSVN yielded poorly soluble nanoparticles, prompting us to use ferritin particles for multimerization, with a 50% occupancy (∼12 copies), creating a second cocktail comprising S2_1.2 in RSVN and the remaining immunogens in ferritin (“Trivax2”, Fig S17).

In mice, Trivax1 elicited low levels of preRSVF cross-reactive antibodies, and sera did not show RSV neutralizing activity in most animals (Fig S18). In contrast, immunization with Trivax2 (Fig 5a) induced robust levels of preRSVF cross-reactive serum levels (Fig 5b) and 6/10 mice showed neutralizing activity (Fig 5c). In these mice, the serum antibody binding responses were equally directed against all three sites (site 0: 32 ± 6 %, site II: 38 ± 7 %, site IV: 30 ± 6 %) (Fig 5d). This is an important finding, as in previous studies mice have been a difficult model to induce serum neutralization with scaffold-based immunogens (*10, 14, 29*). Furthermore, Trivax2 only presents ∼14% of the preRSVF surface area to be targeted by the immune system. While serum neutralization titers in mice remained substantially lower compared to the titers induced by preRSVF (Fig 5c), these results show that vaccine candidates composed of multiple *de novo* proteins can induce physiologically relevant neutralizing serum levels (defined as similar or higher *in vitro* neutralizing activity compared to clinically protective serum concentrations of palivizumab (*43*)).

**Fig 5.**
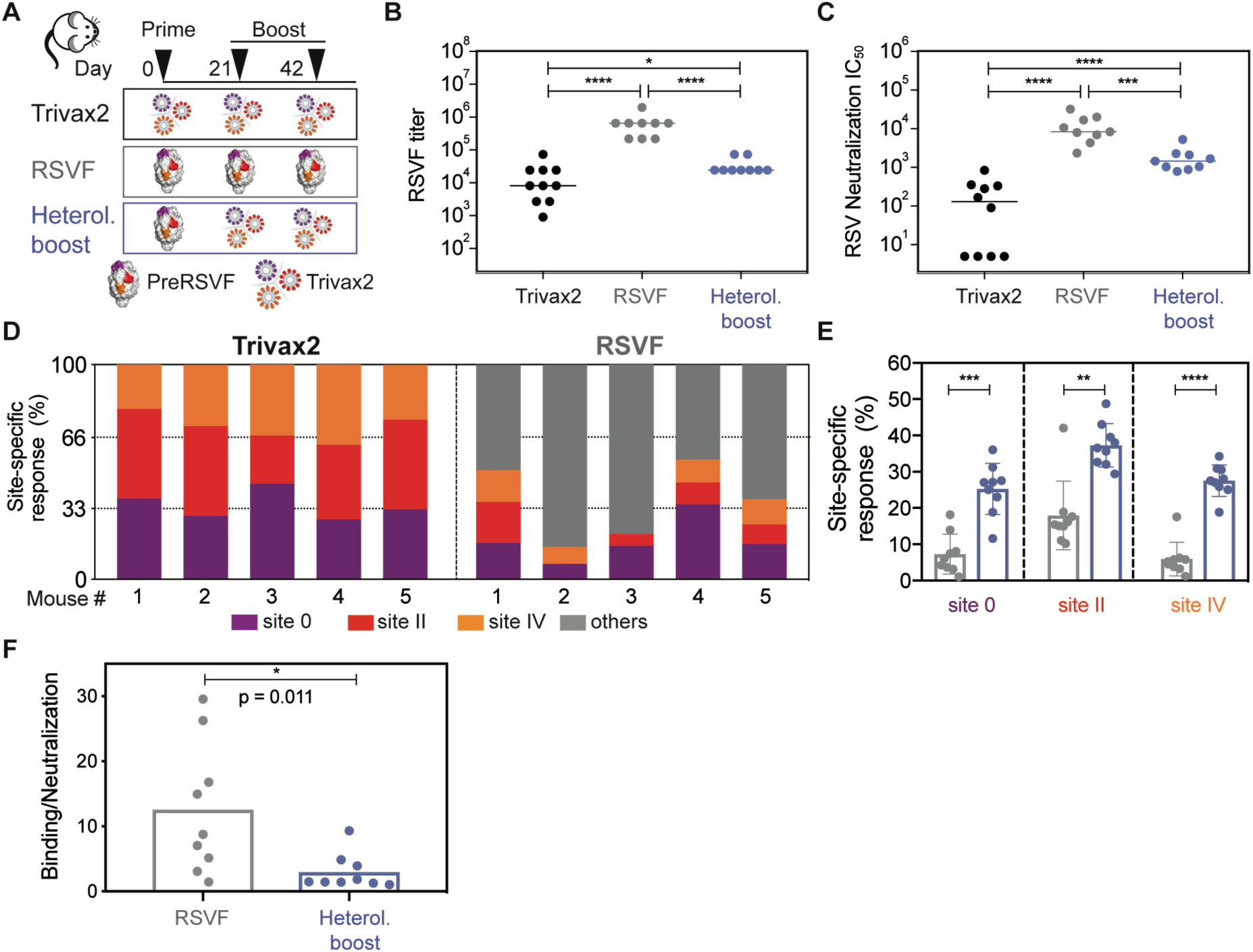
Immunogenicity of Trivax2 in mice. (A) Three groups of mice (n=9-10) were immunized with Trivax2, prefusion RSVF, or as heterologous prime-boost as indicated. All immunogens were formulated in Sigma Adjuvant System. (B) Binding titers measured against prefusion RSVF using ELISA. (C) RSV serum neutralizing titers. (D) Serum composition following three immunizations with Trivax2 or RSVF. Site-specific responses were dissected from serum at day 56 using competition ELISA. (E) Site-specific responses following RSVF immunization compared to a heterologous prime-boost cohort, as measured by a competition SPR assay. On average, comparing Trivax2 with preRSVF, the response against sites 0, II and IV increased 4.4, 2.3 and 5.3-fold, respectively. (F) Ratio of RSVF binding to neutralizing antibody titers. Each symbol represents the ratio of binding IC_50_ to neutralization IC_50_ for an individual animal, the bar represents the mean for each group. Data are representative from at least two independent experiments. Statistics were computed using Mann-Whitney test, where * p < 0.05; ** p < 0.01; *** p < 0.001; **** p < 0.0001.

Given the well-defined epitope specificities induced by the *de novo* designed immunogens, we tested the potential of Trivax2 to boost site-specific responses following a priming immunization with preRSVF (Fig 5a). Strikingly, we found that Trivax2 boosting yielded significantly higher levels of site 0, II and IV antibodies (4.4, 2.3 and 5.3-fold, respectively) compared to boosting immunizations with preRSVF (Fig 5e). This shows that *de novo* proteins can offer a level of control over serum specificities that is out of reach for preRSVF and potentially other immunogens based on viral fusion proteins. While overall serum neutralization titers remained inferior compared to preRSVF boosting (Fig 5c), we found that Trivax2 boosting resulted in a 4.2-fold lower ratio of binding to neutralizing antibodies, a common measure to assess antibody quality (*44, 45*) (Fig 5f). As supported by the significant enhancement of serum responses against sites 0, II and IV upon Trivax2 boosting, these results show the unique potential of *de novo* designed immunogens to increase the quality of the antibody response compared to repeated boosting immunizations with a viral fusion protein.

In parallel, we performed an immunogenicity study in NHPs to test the trivalent cocktail in a closer-to-human antibody repertoire (Fig 6a). This experiment was designed to test the activity of Trivax1 in both RSV-naïve and RSV-experienced animals in an attempt to provide further insights into the ability of computationally designed immunogens to elicit focused antibody responses. The previously designed site II immunogen showed promise in NHPs, but the induced neutralizing titers were low and inconsistent across animals even after repeated immunizations (*10*). In contrast to mice, NHPs immunized with Trivax1 developed robust levels of RSVF cross-reactive serum titers in all animals (Fig 6b), and again antibodies induced were directed against all three epitopes (site 0: 23 ± 5 %, site II: 51 ± 6 %, site IV: 25 ± 5 %) (Fig 6c). Strikingly, 6/7 NHPs showed RSV neutralizing serum levels after a single boosting immunization (median IC_50_ = 312) (Fig 6d). Neutralization titers were maximal at day 84 (median IC_50_ = 408), and measurements were confirmed by an independent laboratory (Fig S19).

**Fig 6.**
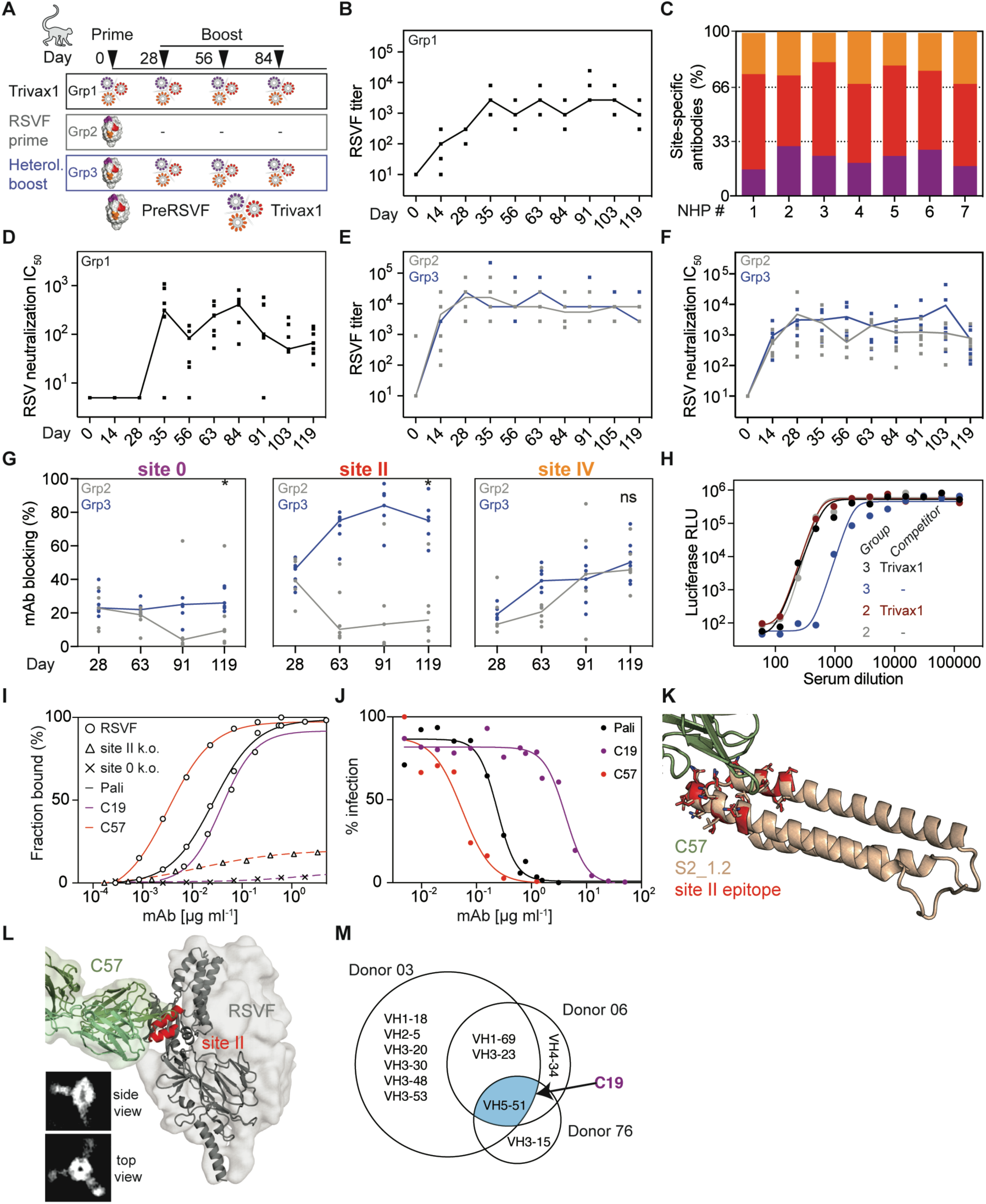
Immunogenicity of Trivax1 in NHPs and analysis of vaccine-induced monoclonal antibodies. (A) NHP immunization scheme. (B) Trivax1 immunized NHPs (group 1) developed robust titers cross-reacting with preRSVF. (C) Site-specific antibody responses for each animal as dissected by competition SPR using D25 (site 0), motavizumab (site II) and 101F (site IV) IgGs as competitors, indicating a balanced response towards all sites. (D) RSV neutralization titers of group 1. (E) PreRSVF titer in group 2 (grey) and 3 (blue). (F) RSV neutralization titers of groups 2 and 3. (G) Dynamics of site-specific antibody levels as measured by competition SPR. Site 0 and site II-specific titers were significantly higher in group 3 compared to 2 following Trivax1 boosting (* p < 0.05, Mann-Whitney U test). (H) RSV neutralization curves upon depletion of day 91 sera with site 0, II, IV-specific scaffolds. 60% of the neutralizing activity is depleted in group 3, whereas no significant decrease is observed in the control group 2. (I) ELISA binding curves of isolated monoclonal antibodies C19 and C57 to preRSVF and site-specific knockouts, in comparison to palivizumab. (J) *In vitro* RSV neutralization of isolated monoclonal antibodies compared to palivizumab. (K) X-ray structure of C57 Fab fragment in complex with the site II scaffold. (L) Model of C57 bound to preRSVF, as confirmed by negative-stain electron microscopy. (M) Lineage analysis (Venn-diagram) of previously identified site 0 nAbs from three different human donors (*41*) showing that VH5-51 derived nAbs were found in all three donors (blue). The elicited site 0 nAb C19 was found to be a close homolog of the VH5-51 lineage in humans.

Beyond naïve subjects, an overarching challenge for vaccine development for pathogens such as influenza, dengue and others, is to focus or reshape pre-existing immunity of broad specificity onto defined neutralizing epitopes that can confer long-lasting protection (*14, 23, 46*). To mimic a serum response of broad specificity towards RSV, we immunized 13 NHPs with preRSVF. All animals developed strong preRSVF-specific titers (Fig 6e) and cross-reactivity with all the epitope-focused immunogens, indicating that epitope-specific antibodies were primed against all three epitopes (Fig S20). Six of those RSVF-primed animals did not receive boosting immunizations so that we could follow the dynamics of epitope-specific antibodies over time (group 2). Seven of the RSVF-primed animals were boosted three times with Trivax1 (group 3) (Fig 6a). While RSVF-specific antibody and neutralization titers remained statistically comparable in both groups (Fig 6e, f), we found that Trivax1 boosting significantly increased antibodies targeting site II and site 0, but not site IV (Fig 6g). In the non-boosted control group, site II and site 0 responses dropped from 37% and 17% at day 28 to 13% and 4% at day 91, respectively (Fig 6g). Compared to the control group, Trivax1 boosting resulted in 6.5-fold higher site II specific responses on day 91 (84% vs 13%), and 6.3-fold higher site 0-specific titers (25% vs 4%) (Fig 6g). In contrast, site IV specific responses increased to similar levels in both groups, 43% and 40% in group 2 and 3, respectively.

To evaluate the functional relevance of reshaping the serum antibody specificities, we depleted site 0, II and IV specific antibodies from pooled sera. In the Trivax1 boosted group, we observed a 60% drop in neutralizing activity as compared to only a 7% drop in the non-boosted control group (Fig 6h). Together, these data largely support the results obtained in mice, showing that epitope-focused immunogens reshaped a serum response of broad specificity towards a focused response that predominantly relied on site 0, II and IV-specific antibodies for RSV neutralization.

Looking further into the molecular basis of the neutralizing activity triggered by the epitope-focused immunogens, we isolated two epitope-specific monoclonal Abs (mAbs) from Trivax1-immunized (group 1) animals using single B cell sorting. Through a panel of binding probes, we confirmed that one mAb targeted antigenic site II (C57) and the other site 0 (C19) (Fig 6i and Fig S21). Importantly, both C57 and C19 neutralized RSV *in vitro* (IC_50_ = 0.03 µg ml^-1^ and IC_50_ = 4.2 µg ml^-1^, respectively). While C19 was less potent, antibody C57 neutralized RSV with approximately one order of magnitude higher potency compared to the clinically used antibody palivizumab, which is similar to the potency of motavizumab and previously reported antibodies induced by a site II epitope-scaffold (*10*) (Fig 6j).

To elucidate the molecular basis for the potent site II-mediated neutralization, we solved a crystal structure of C57 in complex with S2_1.2 at a resolution of 2.2 Å (Fig 6k and Fig S13). C57 recognizes the site II epitope in its native conformation, with a full-atom RMSD of 1.21 Å between the epitope in S2_1.2 and site II in preRSVF. Negative stain EM of the complex C57-preRSVF further confirmed binding to site II, allowing binding of three Fabs per RSVF trimer (Fig 6l).

While RSV nAbs are not known for a strictly constrained VH usage as described for certain influenza and HIV broadly neutralizing antibodies (*47, 48*), an important milestone for computationally designed, epitope-focused immunogens is to elicit nAbs similar to those found in humans upon natural infection or vaccination. In one of the largest RSV antibody isolation campaigns, Gilman et al. isolated 30 site 0-specific RSV nAbs from three human donors (*41*), derived from 11 different VH genes (Fig 6m). Among those, the VH5-51 was the only lineage shared among all three donors, suggesting that, albeit not an obligate lineage, it is a common precursor for site 0 nAbs in humans. We found that the closest human homolog to C19 is indeed the VH5-51 lineage (89% sequence identity), showing that computationally designed immunogens can elicit nAbs with similar properties to those found in humans after viral infection (Fig S21).

In summary, the computational design strategies developed enabled the design of scaffolds presenting epitopes of unprecedented structural complexity with atomic level accuracy. Upon cocktail formulation, the *de novo* designed proteins consistently induced RSV neutralization in naïve animals, mediated through three defined epitopes. In addition, the designed antigens were functional in a heterologous prime-boost immunization regimen, inducing more focused antibody responses towards selected, *bona fide* neutralization epitopes and an overall increased quality of the antibody response.

## Discussion & Conclusions

A major barrier for the *de novo* design of functional proteins is that most functional sites are structurally irregular and/or consist of multiple discontinuous segments. Robust computational methods to endow *de novo* proteins with such complex functional sites have been lacking.

Towards this end, we showcased two *de novo* design strategies to engineer protein scaffolds to present epitopes with high structural complexity. We have shown that using template-based *de novo* design, irregular and discontinuous epitopes were successfully stabilized in heterologous scaffolds. However, this design strategy required extensive optimization by *in vitro* evolution and the designs remained suboptimal regarding their biochemical and biophysical properties. Moreover, this approach lacks the ability to control the topological features of the designed proteins, constituting an important limitation for functional protein design.

To overcome these limitations, we developed the TopoBuilder, a template-free design approach that assembles protein topologies tailored directly to functional sites of interest. Most previous efforts to design functional proteins have focused on transplanting regular, linear helical segments to pre-existing protein structures (*7-9, 27, 28*). Towards more complex binding sites in *de novo* proteins, recent studies reported the design of *de novo* proteins containing multiple segments, but remained limited to regular helical binding motifs embedded in regular helical structures(*49, 50*).

Compared to other approaches, the TopoBuilder has distinctive features and significant advantages to design *de novo* proteins with structurally complex motifs. First, it assembles topologies tailored to the structural requirements of the functional motif from the start of the design process, rather than through the adaptation (and often destabilization) of a protein structure to accommodate the functional site. Second, the topology assembly resulted in designed sequences that stably folded and bound with high affinity without requiring iterative rounds of optimization through directed evolution, as often necessary in computational protein design efforts (*9, 49, 51*).

As to the functional aspect of our design work, we tested whether computationally designed immunogens targeting multiple epitopes could induce physiologically relevant levels of functional antibodies *in vivo*. The elicitation of antibodies targeting conserved epitopes that can mediate broad and potent neutralization remains a central goal for vaccines against pathogens that have frustrated conventional vaccine development efforts (*16*). Structure-based immunogen design holds great promise to overcome this challenge, and different strategies have been developed to elicit focused nAb responses (*15, 17*). For RSV, the development of a prefusion-stabilized version of RSVF has yielded a superior antigen compared to its postfusion counterpart (*44, 52*), largely attributed to the fact that most preRSVF-specific antibodies are neutralizing (*41, 53*). Given that Trivax only presents a small fraction (14%) of the antigenic surface of preRSVF, the substantially lower serum titers, and consequent lower bulk serum neutralization elicited by Trivax in naïve animals may be expected, which will likely require significant optimization in terms of delivery and formulation to increase the magnitude of the response.

Nevertheless, we provide compelling evidence that *de novo* designed proteins presenting three viral epitopes in a cocktail formulation, induced relevant levels of serum neutralization in the majority of naïve mice and NHPs. Beyond bulk serum titers, Trivax offers an unprecedented level of control over antibody specificities to the single epitope level, which has remained out of reach for many immunogens based on viral fusion proteins yet is deemed necessary for the development of next-generation vaccines that require the induction of well-defined antibody specificities (*16, 54*).

Moreover, *de novo* proteins used as boosting immunogen profoundly reshaped the serum composition, leading to increased levels of desirable antibody specificities and an overall improved quality of the antibody response. These findings are important for pathogens such as influenza, where the challenge is to overcome immunodominance hierarchies (*55*), which have been established during repeated natural infections. The ability to selectively boost subdominant nAbs targeting defined, broadly protective epitopes that are surrounded by strain-specific epitopes could overcome long-standing challenges in vaccine development, given that cross-neutralizing antibodies can persist for years once elicited (*56*).

Altogether, this study provides a blueprint for the computational design of epitope-focused vaccines. Besides antigens for viral epitopes, the ability to stabilize structurally complex epitopes in a *de novo* protein with a defined protein topology may prove useful to elicit and isolate novel mAbs against tertiary epitopes with potentially unique allosteric or therapeutic properties. Beyond immunogens, our work presents a new approach for the design of *de novo* functional proteins, enabling the assembly of customized protein topologies tailored to structural and functional requirements of the motif. Ultimately, we show that our *de novo* designed proteins trigger functional phenotypes that are largely out of reach for natural proteins (or close variations of thereof), which is the most important motivation for computational protein design efforts. The ability to design *de novo* proteins presenting functional sites with high structural complexity will be broadly applicable to expand the structural and sequence repertoires, and above all, the functional landscape of natural proteins.

## Contributions

FS, CY and BEC conceived the work and designed the experiments. FS and CY performed computational design and experimental characterization. JB developed the TopoBuilder. JTC, CY, GC, TK, XW and TJ solved x-ray structures. LAA performed NMR characterization and solved the NMR structure. YW, CIC, and YL isolated monoclonal antibodies from NHPs. IK and JPJ performed and analysed samples by electron microscopy. SSV, MG, SR, PC, SG, MV, ED, EO, DD, TG, VM, JFE and MAR performed experiments and analysed data. JTB contributed to the design and planning of animal studies. FS, CY and BEC wrote the manuscript, with input from all authors.

## Data availability

All code used for this study is available through a public github repository: https://github.com/lpdi-epfl/trivalent_cocktail. It contains the TopoBuilder source code, RosettaScripts used for the design, analysis scripts and detailed information on how designs were selected. Structures have been deposited in the Protein Data Bank under accession codes 6S3D (S0_2.126 in complex with D25), 6XWI (S0_2.126 solution NMR structure), 6XXV (S2_1.2 in complex with the Fab fragment of the elicited antibody C57) and 6VTW (S4_2.45 in complex with 101F). Chemical shift data for S0_2.126 were deposited in BMRB under accession code 34481.

## Acknowledgements

We thank William R. Schief, Pablo Gainza and Bruno Lemaitre for helpful discussions and comments on the manuscript. We thank James E. Crowe, Jr. for providing RSVF site IV antibodies used in this study. We thank Kelvin Lau, Aline E. Christine and Florence Pojer in PTPSP facility at EPFL for crystallography support. We thank Andrew McCarthy at ESRF for beam line support, as well as Heather Hotchin, Amy Beierschmitt, and Roberta Palmour from the Behavioural Science Foundation for the NHP immunization and PBMC isolation. We thank the EPFL phenogenomics center (Céline Waldvogel, Raphaël Doenlen) for help with the animal experiments and the protein expression core facility (David Hacker, Laurence Durrer, Soraya Quinche) for help with mammalian protein expression. We thank Davide Demurtas from CIME and Sergey Nazarov from PTBIOEM (EPFL, Lausanne, Switzerland) for electron microscopy support. We thank the flow cytometry core facility at the EPFL for technical support and the gene expression core facility for help with next-generation sequencing. The computational simulations were facilitated by the CSCS Swiss National Supercomputing Centre as well by SCITAS at EPFL.

## Materials and Methods

### Computational design of template-based epitope-focused immunogens

#### Site 0

The structural segments entailing the antigenic site 0 were extracted from the prefusion stabilized RSVF Ds-Cav1 crystal structure, bound to the antibody D25 (PDB 4JHW) (*30*). The epitope consists of two segments: a kinked helical segment (residues 196-212) and a 7-residue loop (residues 63-69).

The MASTER software (*57*) was used to perform structural searches over the Protein Data Bank (PDB, from August 2018), containing 141,920 protein structures, to select template scaffolds with local structural similarities to the site 0 motif. A first search with a C*_α_*RMSD threshold below 2.5 Å failed to identify any close structural matches both in terms of local mimicry as well as global topology features. Thus, a second search was performed, where extra structural elements that support the epitope in its native environment were included as part of the query motif to bias the search towards matches that favoured motif-compatible topologies, rather than those with close local similarities. The extra structural elements included were the two buried helices that directly contact the site 0 in the preRSVF structure (4JHW residues 70-88 and 212-229). The search yielded initially 7,600 matches under a threshold of 5 Å of backbone RMSD, which were subsequently filtered for proteins with a length between 50 and 160 residues, high secondary structure content and accessibility of the epitope for antibody binding. Remaining matches were manually inspected to select template-scaffolds suitable to present the native conformation of antigenic site 0. Subsequently, we selected a computationally designed, highly stable, helical repeat protein (*36*) consisting of 8 regular helices (PDB 5CWJ) with an RMSD of 4.4 Å to the query (2.82 Å for site 0 epitope segments). To avoid steric clashes with the D25 antibody, we truncated the 5CWJ template structure at the N-terminus by 31 residues, resulting in a structure composed of 7 helices.

Using Rosetta FunFolDes (*35*) the truncated 5CWJ topology was folded and designed to stabilize the grafted site 0 epitope recognized by D25. We generated 25,000 designs and selected the top 300 by Rosetta energy score (RE); redesigned backbones that presented obvious flaws, such as low core packing scores, distorted secondary structural elements and buried unsatisfied atoms were discarded. From the top 300 designs, 3 were retained for follow-up iterative cycles of structural relaxation and design using Rosetta FastDesign (*58*), generating a total of 100 designed sequences.

The best 9 designs by Rosetta energy score were recombinantly expressed in E. coli. Two designed sequences derived from the same backbone were successfully expressed and purified. The best variant was named S0_1.1, and subjected to experimental optimization using yeast surface display (Fig S4-S5). In one of the libraries, we found a truncated sequence (S0_1.17) enriched for expression and binding, which served as a template for a second round of computational design (Fig S4-S5). We performed 25,000 folding and design simulations using Rosetta FunFolDes (*35*). The best 300 decoys by total Rosetta energy score were extracted, and each structure was relaxed using the Rosetta Relax application (*59*). We computed the mean total RE, and selected designs that showed a lower energy score than the mean of the design population (RE = −155.2), RMSD drift of the epitope after relaxing of less than 0.7 Å, and a cavity volume <60 Å^3^. We selected one of the best 5 scoring decoys, truncated the C-terminal 29 and N-terminal 23 residues which did not contribute to epitope stabilization, and introduced a disulfide bond between residue 1 and 43. Four sequences were experimentally tested (S0_1.37-40). The best variant according to binding affinity to the target antibodies, S0_1.39, bound with 5 nM affinity to antibody D25, and, importantly, also showed binding to the 5C4 antibody (K_D_ = 5 nM).

#### Site IV

When the design simulations were carried out, there was no structure available of the full RSVF protein in complex with a site IV-specific nAb, nevertheless a peptide epitope of this site recognized by the 101F nAb had been previously reported (PDB 3O41) (*31*).

The crystallized peptide-epitope corresponds to the residues 429-434 of the RSVF protein. Structurally, the 101F-bound peptide-epitope adopts a bulged strand and several studies suggest that 101F recognition extends beyond the linear *β*-strand segment, contacting other residues located in antigenic site IV (*60*). Despite the apparent structural simplicity of the epitope, structural searches for designable scaffolds failed to yield promising starting templates. However, we noticed that the antigenic site IV of RSVF is self-contained within an individual domain that could potentially be excised and designed as a soluble folded protein. To maximize these contacts, we first truncated the seemingly self-contained region from RSVF pre-fusion structure (PDB 4JHW, residue: 402-459) forming a *β*-sandwich and containing site IV. We used Rosetta FastDesign to optimize the core positions of this minimal topology, obtaining our initial design (dubbed S4_wt). However, S4_wt did not show a funnel-shaped energy landscape in Rosetta *ab initio* simulations, and we were unable to recombinantly express S4_wt in *E.coli*.

In an attempt to improve the conformation and stabilization of S4_wt, we used Rosetta FunFolDes to fold and design this topology, while keeping the conformation of the site IV epitope fixed. Out of 25,000 simulations, the top 1% decoys according to RE score and overall RMSD were manually inspected, and 12 designed sequences were selected for recombinant expression in *E.coli*. One of the designs (S4_1.1) initially bound to 101F with a K_D_ ∼ 85 μM, and was subjected to experimental optimization through saturation mutagenesis on the chosen residues proximal to the contact region of the scaffold-101F antibody interface (Fig S2-3). By the combining the enriched mutations, we found a double mutant (S4_1.5) with a binding affinity improved by three orders of magnitude (K_D_ = 35 nM)

Rosetta scripts and analysis scripts use to perform the computational design are available in the GitHub repository accompanying this paper, together with detailed instructions to run the code.

### TopoBuilder - Motif-centric *de novo* design

Given the limited availability of suitable starting templates to host structurally complex motifs such as site 0 and site IV, we developed a template-free design protocol, which we named TopoBuilder. In contrast to adapting existing topologies to accommodate the epitope, the design goal is to build protein scaffolds around the epitope from scratch, using idealized secondary structures (beta strands and alpha helices). The TopoBuilder consists of three stages:

(1) Topological sampling in 2D space

To quickly define the fold space compatible with the target structural motif we use the αβα-Form topology definition scheme, a string-based descriptor that allows the extensive enumeration of multilayer protein topologies with alternating secondary structure elements and all possible connections between the secondary structural elements (*37, 38*). Putative folds are then selected according to basic topological rules (e.g. lack of crossover loops and chain directionality of the functional motif).

(2) 3D projection and parametric sampling

The selected 2D topologies are projected into the 3D space by assembling idealized secondary structures (SSE) around the fixed functional motif. These 3D structures, referred to as ‘sketches’, are further refined by coarsely sampling structural features of the fold (e. g. distances and orientations between SSEs) using parametric sampling.

The length, orientation and 3D-positioning are defined by the user for each secondary structure with respect to the epitope, which is extracted from its native environment. The topologies built were designed to meet the following criteria: (1) Small, globular proteins with a high-contact order between secondary structures and the epitope, to allow for stable folding and accurate stabilization of the epitope in its native conformation; (2) Context mimicry, i.e. respecting shape constraints of the epitope in its native context (Fig S6). For assembling the topology, the default distances between alpha helices was set to 11 Å and for adjacent beta strands was 5 Å. For each discontinuous structural sketch, a connectivity between the secondary structural elements was defined and loop lengths were selected to connect the secondary structure elements with the minimal number of residues that can cover a given distance, while maintaining proper backbone geometries.

For site 0, the short helix of S0_1.39 preceding the epitope loop segment was kept, and a third helix was placed on the backside of the epitope to: (1) provide a core to the protein and (2) allow for the proper connectivity between the secondary structures. A total of three different orientations (25°, 0° and −45° degrees to the plane formed by site 0) where tested for the designed supporting alpha helix (Fig 3 and Fig S10).

In the case of site IV, the known binding region to 101F (residues 428F-434F) was extracted from prefusion RSVF (PDB 4JHW). Three antiparallel beta strands, pairing with the epitope, plus an alpha helix on the buried side, were assembled around the 101F epitope. Three different configurations (45°, (−45°,0°,10°) and −45° degrees with respect to the β-sheet) were sampled parametrically for the alpha helix (Fig 3 and Fig S7).

(3) Flexible backbone sampling, sequence design and selection To refine the structural features of the sketches at the all-atom level and design sequences that stabilize these structures, we use Rosetta FunFolDes as described before (*10, 35*).

Approximately 25,000 trajectories were generated for each sketch. The newly generated backbones were further subjected to layer-based FastDesign (*58*), where each amino acid position was assigned a layer (combining ‘core’, ‘boundary’, ‘surface’ and ‘sheet’ or ‘helix’) on the basis of its exposure to solvent and secondary structure type, which dictated the allowed amino acid types at that position.

After iterative cycles of sequence design, unconstrained FastRelax (*61*) (i.e. sidechain repacking and backbone minimization) was applied over the designs to evaluate their conformational stability of the epitope region. After each relax cycle, structural changes upon the epitope region were evaluated (epitope RMSD drift). Designs with epitope RMSD drifts higher than 1.2 Å were discarded. Designs were also ranked and selected according to hydrophobic core packing (packstat score), with a cutoff of 0.5 for site 0 and 0.6 for the site IV design series, and a cavity volume of < 50 Å^3^. Between 1,000 and 10,000 of the designed sequences were generated from this computational protocol.

Rosetta scripts and analysis scripts to perform the designs are available in the GitHub repository accompanying this paper, together with detailed instructions to run the code.

### Mouse immunizations

All animal experiments were approved by the Veterinary Authority of the Canton of Vaud (Switzerland) according to Swiss regulations of animal welfare (animal protocol number 3074). Female Balb/c mice (6-week old) were purchased from Janvier labs. The immunogens were thawed on ice, mixed with equal volumes of adjuvant (Sigma Adjuvant System, Sigma) according to the manufacturer’s instructions. Mice were injected subcutaneously with 100 μl vaccine formulation, containing in total 10 μg of immunogen (equimolar ratios of each immunogen for Trivax immunizations).

Immunizations were performed on day 0, 21 and 42. 100-200 μl of blood was drawn on day 0, 14 and 35. Mice were euthanized at day 56 and blood was taken by cardiac puncture.

### NHP immunizations

Twenty-one African green monkeys (AGM, 3-4 years) were divided into three experimental groups with at least two animals of each sex. AGMs were pre-screened as seronegative against prefusion RSVF (preRSVF) by ELISA. Vaccines were prepared 1 hour before injection, containing 50 μg preRSVF or 300 μg Trivax1 in 0.5 ml PBS, mixed with 0.5 ml alum adjuvant (Alhydrogel, Invivogen) for each animal. AGMs were immunized intramuscularly at day 0, 28, 56, and 84. Blood was drawn at days 14, 28, 35, 56, 63, 84, 91, 105 and 119.

### RSV neutralization assay

The RSV neutralization assay was performed as described previously (*14*). Briefly, Hep2 cells were seeded in Corning 96-well tissue culture plates (Sigma) at the density of 40,000 cells/well in 100 μl of Minimum Essential Medium (MEM, Gibco) supplemented with 10% FBS (Gibco), L-glutamine 2 mM (Gibco) and penicillin-streptomycin (Gibco), and grown overnight at 37 °C with 5% CO_2_. Serial dilutions of heat-inactivated sera were prepared in MEM without phenol red (M0, Life Technologies, supplemented with 2 mM L-glutamine and penicillin/streptomycin) and were incubated with 800 pfu/well (final MOI 0.01) RSV-Luc (Long strain carrying a luciferase gene) for 1 hour at 37 °C. Serum-virus mixture was added to Hep2 cell layer, and incubated for 48 hours. Cells were lysed in lysis buffer supplemented with 1 µg/ml luciferin (Sigma) and 2 mM ATP (Sigma), and luminescence signal was read on a Tecan Infinite 500 plate reader. The neutralization curve was plotted and fitted using the GraphPad version 7.0 with a variable slope fitting model, weighted by 1/Y^2^. Similar to a previous study by McLellan and colleagues (*52*), we used the monoclonal antibody palivizumab to benchmark the serum neutralization activity. Clinical studies in humans have shown that, when palivizumab was dosed at a concentration of 15 mg/kg, trough serum concentrations reach ∼40 μg/ml, which provides protection in infants (*43*). Similarly, palivizumab serum concentrations of 25-30 μg/ml were shown to afford protection in cotton rats, yielding 99% reduction in RSV lung titer (*62*). In our neutralization assay, 40 μg/ml of palivizumab yields an IC_50_ of ∼100 (palivizumab IC_50_ = 0.48 ± 0.04 μg/ml). Thus, IC_50_ values of ∼100 are considered physiologically relevant neutralization titer.

### Serum fractionation

Monomeric Trivax1 immunogens (S2_1.2, S0_1.39 and S4_1.5) were used to deplete the site 0, II and IV specific antibodies in immunized sera. HisPurTM Ni-NTA resin slurry (Thermo Scientific) was washed with PBS containing 10 mM imidazole. Approximately 1 mg of each immunogen was immobilized on Ni-NTA resin, followed by two wash steps to remove unbound scaffold. 60 μl of sera pooled from all animals within the same group were diluted to a final volume of 600 μl in the wash buffer, and incubated overnight at 4 °C with 500 μl Ni-NTA resin slurry. As control, the same amount of sera was incubated with Ni-NTA resin that did not contain scaffolds. Resin was pelleted down at 13,000 rpm for 5 minutes, and the supernatant (depleted sera) was collected for subsequent neutralization assays.

### Cell staining and single-cell flow cytometry sorting

Peripheral blood was collected at day 91, followed by PBMC preparation. PBMCs were stained by a cocktail of antibodies for identifying memory B cells as described previously (*63*). Briefly, frozen PBMCs were thawed and treated with 10,000 U/ml DNase I (Roche) in RPMI 1640 supplemented with 10% fetal bovine serum (FBS) media, followed by Aqua Dead Cell Staining (Life Technologies). A cocktail of antibodies containing CD3 (APC-Cy7; SP34-2, BD Pharmingen), CD8 (Pacific blue; RPA-T8, BD Pharmingen), CD14 (Qdot 605; M5E2, VRC), CD20 (PE-Alexa Fluor 700; 2H7, VRC), CD27 (PE-Cy7; M-T271, BD Pharmingen), IgG (FITC; G18-145, BD Pharmingen), and IgM (PE-Cy5; G20-12, BD Pharmingen) was used to stain the PBMCs. To sort RSVF-specific B cells, preRSVF conjugated with allophycocyanin conjugate (APC) was incorporated in the above-stated antibody cocktail. Following staining, the cells were sorted at a single-cell density into 96-well plates with lysis buffer using a four-laser FACS Aria III cell sorter. RSVF-specific memory B cells were defined as CD3^−^CD8^−^Aqua Blue^−^CD14^−^CD20^+^IgG^+^CD27^+^IgM^−^RSVF*^hi^*.

### Single-cell RT-PCR, antibody cloning and expression

The sorted cells were lysed by the lysis buffer followed by single-cell reverse transcription and PCR reactions to amplify Ig sequences as described previously (*63, 64*). Briefly, in each well containing the lysed cell, 150 ng random hexamers, 0.4 mM dNTPs, 100 U SuperScript III (Life Technologies), and 3.5 μl of water were added, followed by incubation at 42 °C for 10 min, 25 °C for 10 min, 50 °C for 60 min, and 94 °C for 5 min to generate cDNA. Nested PCR was then performed with 2 μl of cDNA in 25 μl reactions with the HotStar Taq Plus Kit (QIAGEN), using 5’ leader sequence–specific and 3’ IgG-specific primers (*64*). In the second round of nested PCR, 1.75 μl of the 1^st^ PCR product was used as template. All nested PCRs were incubated at 94 °C for 5 min followed by 50 cycles of 94 °C for 30 s, 50 °C (heavy chain) or 52 °C (kappa or lambda chain) for 45 s, and 70 °C for 1 min with a final elongation at 70 °C for 10 min before cooling to 4 °C. Nested PCR products were evaluated on 2% 96-well E Gel (Life Technologies), and positive wells with a specific band of ∼450 bp were purified and sequenced. After sequencing verification of productive V(D)J sequences, the Ig variable domains of selected B cell clones were cloned into Ab IgG heavy and light chain expression vector, respectively, as described previously (*63, 65*). The cloning PCR reaction was performed in a total volume of 50 μl with 1 μl of the 2^nd^ nested PCR product as template, using Expand High-Fidelity PCR Kit according to the manufacturer’s instructions (Roche). The PCR reaction consisted of 5 μl of 10x reaction buffer, 1 μl of 10 mM dNTPs, 1 μl each of 25 μM of 3’ and 5’ cloning primers, 1 μl of DNA polymerase and water to 50 μl. The cloning primers were described previously (*64*). The PCR reaction had an initial denaturation at 95 °C for 3 min, followed by 20 cycles of 95 °C for 30 s, 50 °C for 30 s and 68 °C for 2 min. There was a final elongation at 68 °C for 8 min before cooling to 4 °C followed by evaluation on 1% agarose gel. Positive bands (∼400 bp for heavy chain VDJ and ∼350 bp for light chain VJ) were then purified, digested with restriction enzymes, and ligated into eukaryotic expression vectors containing human Igγ1H, Igγ2, or Igκ1 L chain Ab expression cassettes (*66, 67*). After transformation of ligated products into competent cells, bacterial colonies were sequenced for the insertion of correct sequences. The antibody expression vectors were co-transfected into 293F cells and the cell culture supernatants containing secreted antibody IgG were purified by columns packed in-house containing Protein A Sepharose beads (GE Healthcare).

### Analysis of RSVF-binding Abs

The purified antibodies were tested for specificity by enzyme-linked immunosorbent assays (ELISAs). RSVF, the designed epitope scaffolds or site-specific knockout probes of RSVF (*41*) were coated onto plates at 2 μg/ml in PBS overnight at 4 °C. After blocking with PBS + 0.05% Tween 20 + 5% milk, mAbs were added into wells in 3-fold dilution series and incubated for 1 h at RT. Goat anti-human Fc-HRP conjugate (Invitrogen) was used at a 1:10,000 dilution in blocking buffer for 1 h at RT. The bound mAb was detected with 100 μl/well of TMB substrate (Life Technologies) for 1 min before the addition of 100 μl of 3% H_2_SO_4_ to stop the reaction. The optical density (OD) was measured at 450 nm. Between each step of the ELISA, plates were washed four times with PBS supplemented with 0.05% Tween 20.

### Site saturation mutagenesis library (SSM)

The SSM library was assembled by overhang PCR, in which 11 selected positions surrounding the epitope in the S4_1.1 design model were allowed to mutate to all 20 amino acids, with one mutation allowed at a time. Each of the 11 libraries was assembled by primers (Table S1) containing the degenerate codon ‘NNK’ at the selected position. All 11 libraries were pooled, and transformed into EBY-100 yeast strain with a transformation efficiency of 1×10^6^ transformants.

### Combinatorial library

Combinatorial sequence libraries were constructed by assembling multiple overlapping primers (Table S2) containing degenerate codons at selected positions for combinatorial sampling of hydrophobic amino acids in the protein core. The theoretical diversity was between 1×10^6^ and 5×10^6^. Primers were mixed (10 µM each), and assembled in a PCR reaction (55 °C annealing for 30 sec, 72 °C extension time for 1 min, 25 cycles). To amplify full-length assembled products, a second PCR reaction was performed, with forward and reverse primers specific for the full-length product. The PCR product was desalted, and transformed into EBY-100 yeast strain with a transformation efficiency of at least 1×10^7^ transformants (*68*).

### Protein expression and purification

#### Designed scaffolds

All genes of designed proteins were purchased as DNA fragments from Twist Bioscience, and cloned via Gibson assembly into either pET11b or pET21b bacterial expression vectors. Plasmids were transformed into *E.coli* BL21 (DE3) (Merck) and grown overnight in LB media. For protein expression, precultures were diluted 1:100 and grown at 37 °C until the OD_600_ reached 0.6, followed by the addition of 1 mM IPTG to induce protein expression. Cultures were harvested after 12-16 hours at 22°C. Pellets were resuspended in lysis buffer (50 mM Tris, pH 7.5, 500 mM NaCl, 5% Glycerol, 1 mg/ml lysozyme, 1 mM PMSF, 1 µg/ml DNase) and sonicated on ice for a total of 12 minutes, in intervals of 15 seconds sonication followed by 45 seconds pause. The lysates were clarified by centrifugation (20,000 rpm, 20 minutes) and purified via Ni-NTA affinity chromatography followed by size exclusion on a HiLoad 16/600 Superdex 75 column (GE Healthcare) in PBS buffer.

#### Antibodies - IgG and Fab constructs

Plasmids encoding cDNAs for the heavy chain of IgG were purchased from Genscript and cloned into the pFUSE-CHIg-hG1 vector (Invivogen). Heavy and light chain DNA sequences of antibody fragments (Fab) were purchased from Twist Bioscience and cloned separately into the pHLsec mammalian expression vector (Addgene, #99845) via Gibson assembly. HEK293-F cells were transfected in an equal ratio with heavy and light chains, and cultured in FreeStyle medium (Gibco) for 7 days. Supernatants were collected by centrifugation and purified using a 1 ml HiTrap Protein A HP column (GE Healthcare) for IgG expression and 5 ml kappa-select column (GE Healthcare) for Fab purification. Bound antibodies/Fabs were eluted with 0.1 M glycine buffer (pH 2.7), immediately neutralized by 1 M Tris ethylamine buffer (pH 9), and buffer exchanged to PBS.

#### Prefusion stabilized RSVF

The construct encoding the thermostabilized the preRSVF protein corresponds to the sc9-10 DS-Cav1 A149C Y458C S46G E92D S215P K465Q variant (referred to as DS2) reported by Joyce and colleagues (*69*). The sequence was codon-optimized for mammalian cell expression and cloned into the pHCMV-1 vector flanked with two C-terminal Strep-Tag II and one 8x His tag. Expression and purification were performed as described previously (*14*).

#### Nanoring-based immunogens

The full-length N gene derived from the human RSV strain Long, ATCC VR-26 (GenBank accession number AY911262.1) was PCR amplified and cloned into pET28a+ at NcoI-XhoI sites to obtain the pET-N plasmid. Immunogens presenting sites II and IV epitopes were cloned into the pET-N plasmid at NcoI site as pET-S2_1.2-N and pET-S4_1.5-N, respectively, and the site 0 immunogen was cloned at the pET-N XhoI site to obtain pET-N-S0_1.39. Expression and purification of the nanoring fusion proteins was performed as described previously (*14*).

#### Ferritin-based immunogens

The gene encoding Helicobacter pylori ferritin (GenBank ID: QAB33511.1) was cloned into the pHLsec vector for mammalian expression, with an N-terminal 6x His Tag. The sequence of the designed immunogens (S0_2.126 and S4_2.45) were cloned upstream of the ferritin gene, spaced by a GGGGS linker. Ferritin particulate immunogens were produced by co-transfecting a 1:1 stochiometric ratio of ferritin and immunogen-ferritin in HEK-293F cells, as previously described for other immunogen-nanoparticle fusion constructs (*70*). The supernatant was collected 7-days post transfection and purified via Ni-NTA affinity chromatography and size exclusion on a Superose 6 increase 10/300 GL column (GE).

### Negative-stain transmission electron microscopy

#### Sample preparation

RSVN and Ferritin-based nanoparticles were diluted to a concentration of 0.015 mg/ml. The samples were absorbed on carbon-coated copper grid (EMS, Hatfield, PA, United States) for 3 mins, washed with deionized water and stained with freshly prepared 0.75% uranyl formate.

#### Data acquisition

The samples were viewed under an F20 electron microscope (Thermo Fisher) operated at 200 kV. Digital images were collected using a direct detector camera Falcon III (Thermo Fisher) with the set-up of 4098 X 4098 pixels. The homogeneity and coverage of staining samples on the grid was first visualized at low magnification mode before automatic data collection. Automatic data collection was performed using EPU software (Thermo Fisher) at a nominal magnification of 50,000X, corresponding to pixel size of 2 Å, and defocus range from −1 µm to −2 µm.

#### Image processing

CTFFIND4 program (*71*) was used to estimate the contrast transfer function for each collected image. Around 1000 particles were manually selected using the installed package XMIPP within SCIPION framework (*72*). Manually picked particles were served as input for XMIPP auto-picking utility, resulting in at least 10,000 particles.

Selected particles were extracted with the box size of 100 pixels and subjected for three rounds of reference-free 2D classification without CTF correction using RELION-3.0 Beta suite (*73*).

#### RSVF-Fabs complex formation and negative stain EM

20 μg of RSVF trimer was incubated overnight at 4 °C with 80 µg of Fabs (Motavizumab, D25 or 101F). For complex formation with all three monoclonal Fabs, 80 µg of each Fab was used. Complexes were purified on a Superose 6 Increase 10/300 column using an Äkta Pure system (GE Healthcare) in TBS buffer. The main fraction containing the complex was directly used for negative stain EM.

Purified complexes of RSVF and Fabs were deposited at approximately 0.02 mg/ml onto carbon-coated copper grids and stained with 2% uranyl formate. Images were collected with a field-emission FEI Tecnai F20 electron microscope operating at 200 kV. Images were acquired with an Orius charge-coupled device (CCD) camera (Gatan Inc.) at a calibrated magnification of ×34,483, resulting in a pixel size of 2.71 Å. For the complexes of RSVF with a single Fab, approximately 2,000 particles were manually selected with Cryosparc2 (*74*). Two rounds of 2D classification of particle images were performed with 20 classes allowed. For the complexes of RSVF with D25, Motavizumab and 101F Fabs, approximately 330,000 particles were picked using Relion 3.0 (*73*) and subsequently imported to Cryosparc2 for two rounds of 2D classification with 50 classes allowed.

### Determining binding affinities by Surface plasmon resonance (SPR)

SPR measurements were performed on a Biacore 8K (GE Healthcare) with HBS-EP+ as running buffer (10 mM HEPES pH 7.4, 150 mM NaCl, 3 mM EDTA, 0.005% v/v Surfactant P20, GE Healthcare). Ligands were immobilized on a CM5 chip (GE Healthcare # 29104988) via amine coupling. Approximately 2000 response units (RU) of IgG were immobilized, and designed monomeric proteins were injected as analyte in two-fold serial dilutions. The flow rate was 30 µl/min for a contact time of 120 seconds followed by 400 seconds dissociation time. After each injection, surface was regenerated using 3 M magnesium chloride (101F as immobilized ligand) or 0.1 M glycine at pH 4.0 (Motavizumab and D25 IgG as an immobilized ligand). Data were fitted using 1:1 Langmuir binding model within the Biacore 8K analysis software (GE Healthcare #29310604).

### Dissection of serum antibody specificities by SPR

To quantify the epitope-specific antibody responses in bulk serum from immunized animals, we performed an SPR competition assay with the monoclonal antibodies (D25, Motavizumab and 101F) as described previously (*14*). Briefly, approximately 400 RU of preRSVF were immobilized on a CM5 chip via amine coupling, and serum diluted 1:10 in running buffer was injected to measure the total response (RU_non-blocked surface_). After chip regeneration using 50 mM NaOH, the site 0/II/IV epitopes were blocked by injecting saturating amounts of either D25, Motavizumab, or 101F IgG, and serum was injected again to quantify residual response (RU_blocked surface_).

The delta serum response *ΔSR* was calculated as follows:

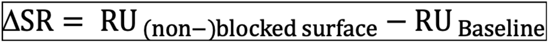

Percent blocking was calculated for each site as:

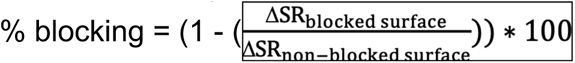

### Size exclusion chromatography multi-angle light scattering (SEC-MALS)

Size exclusion chromatography with an online multi-angle light scattering device (miniDAWN TREOS, Wyatt) was used to determine the oligomeric state and molecular weight for the protein in solution. Purified proteins were concentrated to 1 mg/ml in PBS (pH 7.4), and 100 µl of the sample was injected into a Superdex 75 300/10 GL column (GE Healthcare) with a flow rate of 0.5 ml/min, and UV_280_ and light scattering signals were recorded. Molecular weight was determined using the ASTRA software (version 6.1, Wyatt).

### Circular Dichroism

Far-UV circular dichroism spectra were measured using a Jasco-815 spectrometer in a 1 mm path-length cuvette. The protein samples were prepared in 10 mM sodium phosphate buffer at a protein concentration of 30 µM. Wavelengths between 190 nm and 250 nm were recorded with a scanning speed of 20 nm min^-1^ and a response time of 0.125 secs. All spectra were averaged 2 times and corrected for buffer absorption. Temperature ramping melts were performed from 25 to 90 °C with an increment of 2 °C/min in presence or absence of 5 mM TCEP reducing agent. Thermal denaturation curves were plotted by the change of ellipticity at the global curve minimum to calculate the melting temperature (T_m_).

### Yeast surface display

Libraries of linear DNA fragments encoding variants of the designed proteins were transformed together with linearized pCTcon2 vector (Addgene #41843) based on the protocol previously described by Chao and colleagues (*68*). Transformation procedures generally yielded ∼10^7^ transformants. The transformed cells were passaged twice in SDCAA medium before induction. To induce cell surface expression, cells were centrifuged at 7,000 rpm. for 1 min, washed with induction media (SGCAA) and resuspended in 100 ml SGCAA with a cell density of 1 x 10^7^ cells/ml SGCAA. Cells were grown overnight at 30 °C in SGCAA medium. Induced cells were washed in cold wash buffer (PBS + 0.05% BSA) and labelled with various concentration of target IgG or Fab (101F, D25, and 5C4) at 4 °C. After one hour of incubation, cells were washed twice with wash buffer and then incubated with FITC-conjugated anti-cMyc antibody and PE-conjugated anti-human Fc (BioLegend, #342303) or PE-conjugated anti-Fab (Thermo Scientific, #MA1-10377) for an additional 30 min. Cells were washed and sorted using a SONY SH800 flow cytometer in ‘ultra-purity’ mode. The sorted cells were recovered in SDCAA medium, and grown for 1-2 days at 30 °C.

In order to select stably folded proteins, we washed the induced cells with TBS buffer (20 mM Tris, 100 mM NaCl, pH 8.0) three times and resuspended in 0.5 ml of TBS buffer containing 1 µM of chymotrypsin. After incubating five minutes at 30 °C, the reaction was quenched by adding 1 ml of wash buffer, followed by five wash steps. Cells were then labelled with primary and secondary antibodies as described above.

### ELISA

96-well plates (Nunc MediSorp plates; Thermo Scientific) were coated overnight at 4 °C with 50 ng/well of purified antigen (recombinant RSVF or designed immunogens) in coating buffer (100 mM sodium bicarbonate, pH 9) in 100 μl total volume. Following overnight incubation, wells were blocked with blocking buffer (PBS + 0.05% Tween 20 (PBST) containing 5% skim milk (Sigma)) for 2 hours at room temperature. Plates were washed five times with PBST. 3-fold serial dilutions were prepared and added to the plates in duplicates, and incubated at room temperature for 1 hour. After washing, anti-mouse (abcam, #99617) or anti-monkey (abcam, #112767) HRP-conjugated secondary antibody were diluted 1:1,500 or 1:10,000, respectively, in blocking buffer and incubated for 1 hour. Additional five wash steps were performed before adding 100 µl/well of Pierce TMB substrate (Thermo Scientific). The reaction was terminated by adding an equal volume of 2 M sulfuric acid. The absorbance at 450 nm was measured on a Tecan Safire 2 plate reader, and the antigen specific titers were determined as the reciprocal of the serum dilution yielding a signal two-fold above the background.

### Competition ELISA to dissect serum specificities

Competition ELISA was performed as described previously (*14*). Briefly, sera were serially diluted, mixed with 100 μg/ml competitor antigen and incubated overnight at 4 °C, before following the ELISA protocol as described above. The epitope-scaffolds (S2_1.2, S0_2.126 and S4_2.45), each multimerized on a self-assembling nanoparticle served as competitor antigens to dissect site II, 0 and IV specific responses, respectively. Naked nanoparticles served as control for non-specific competition. Reactivity against RSVF was assayed both in presence and absence of site-specific competitors. All samples were assayed in duplicates. Curves were plotted in GraphPad Prism 7.0. The area under the curve (AUC) was computed for the site-specific and control competitors to calculate % competition for each site.

### NMR

Protein samples for NMR were prepared in 10 mM sodium phosphate buffer, 50 mM sodium chloride at pH 7 with the protein concentration around 500 µM. All NMR experiments were carried out in an 18.8T (800 MHz proton Larmor frequency) Bruker spectrometer equipped with a CPTC ^1^H,^13^C,^15^N 5 mm cryoprobe and an Avance III console. Experiments for backbone resonance assignment consisted in standard triple resonance spectra HNCA, HN(CO)CA, HNCO, HN(CO)CA, CBCA(CO)NH and HNCACB acquired on a 0.5 mM sample doubly labelled with ^13^C and ^15^N (*75*). Sidechain assignments were obtained from HCCH-TOCSY experiments acquired on the same sample plus HNHA, NOESY-^15^N-HSQC and TOCSY-^15^N-HSQC acquired on a ^15^N-labeled sample, and 2D ^1^H-^1^H TOCSY and NOESY spectra acquired on unlabelled samples prepared in 10% and 100% D_2_O solutions. The ^15^N-resolved NOESY, 2D NOESY in 10% D_2_O and 2D NOESY in 100% D_2_O spectra were further used for structure calculations.

Spectra for backbone assignments were acquired with 40 increments in the ^15^N dimension and 128 increments in the ^13^C dimension, and processed with 128 and 256 points by using linear prediction. The HCCH-TOCSY was recorded with 64-128 increments in the ^13^C dimensions and processed with twice the number of points. ^15^N-resolved NOESY and TOCSY spectra were acquired with 64 and 128 increments in the indirect ^15^N and ^1^H dimensions, and processed with twice the number of points. 2D ^1^H-^1^H NOESY and TOCSY spectra were acquired with 256 increments in the indirect dimension, processed with 512 points. Mixing times for NOESY spectra were 120 ms and TOCSY spin locks were 60 ms. All spectra were acquired and processed with Bruker’s TopSpin 3.0 (acquisition with standard pulse programs) and analyzed manually with the program CARA (http://cara.nmr.ch/doku.php/home) to obtain backbone and sidechain resonance assignments. Peak picking and assignment of NOESY spectra (a ^15^N-resolved NOESY and a 2D NOESY) were performed automatically with the program UNIO-ATNOS/CANDID (*76, 77*) coupled to Cyana 2.1 (*78*), using standard settings in both programs. The run was complemented with dihedral angles derived from chemical shifts with Talos-n (*79*). The refinement statistics of the structure are presented in Table S7.

### X-ray crystallization and structural determination

#### Co-crystallization of complex D25 Fab with S0_2.126

After overnight incubation at 4 °C, the S0_2.126/D25 Fab complex was purified by size exclusion chromatography using a Superdex200 26 600 (GE Healthcare) equilibrated in 10 mM Tris pH 8, 100 mM NaCl and subsequently concentrated to ∼10 mg/ml (Amicon Ultra-15, MWCO 3,000). Crystals were grown at 291K using the sitting-drop vapor-diffusion method in drops containing 1 μl purified protein mixed with 1 μl reservoir solution containing 10% PEG 8000, 100 mM HEPES pH 7.5, and 200 mM calcium acetate. For cryo protection, crystals were briefly swished through mother liquor containing 20% ethylene glycol.

#### Data collection and structural determination of the S0_2.126/D25 Fab complex

Diffraction data were recorded at ESRF beamline ID30B. Data integration was performed by XDS (*80*) and a high-resolution cut at I/σ=1 was applied. The dataset contained a strong off-origin peak in the Patterson function (88% height rel. to origin) corresponding to a pseudo translational symmetry of ½, 0, ½. The structure was determined by molecular replacement with Phenix Phaser (*81*) using the D25 structure (*30*) (PDB 4JHW) as a search model. Manual model building was performed using Coot (*82*), and automated refinement using Phenix Refine (*83*). After several rounds of automated refinement and manual building, paired refinement (*84*) determined the resolution cut-off for final refinement. The final refinement statistics, native data and phasing statistics are summarized in Table S8.

#### Co-crystallization of complex 101F Fab with S4_2.45

The complex of S4_2.45 with the 101F Fab was prepared by mixing two proteins in 2:1 molar ratio for 1 hour at 4 °C, followed by SEC using a Superdex-75 column.

Complexes of S4_2.45 with the 101F Fab were verified by SDS–PAGE. Complexes were subsequently concentrated to 6–8 mg/ml. Crystals were grown using hanging drops vapor-diffusion method at 20 °C. The S4_2.45/101F protein complex was mixed with equal volume of a well solution containing 0.2 M Magnesium acetate, 0.1 M Sodium cacodylate pH 6.5, 20% (w/v) PEG 8000. Native crystals were transferred to a cryoprotectant solution of 0.2 M Magnesium acetate, 0.1 M Sodium cacodylate pH 6.5, 20% (w/v) PEG 8000 and 15% glycerol, followed by flash-cooling in liquid nitrogen.

#### Data collection and structural determination of the S4_2.45/101F Fab complex

Diffraction data were collected at SSRL facility, BL9-2 beamline at the SLAC National Accelerator Laboratory. The crystals belonged to space group P3_2_21. The diffraction data were initially processed to 2.6 Å with X-ray Detector Software (XDS) (*80*)(Table S9).

Molecular replacement searches were conducted with the program Phenix Phaser (*81*) using 101F Fab model (PDB 3O41) and S4_2.45/101F Fab computational model generated from superimposing the epitope region of S4_2.45 with the peptide-bound structure (PDB 3O41), and yielded clear molecular replacement solutions. Initial refinement provided a R_free_ of 42.43% and R_work_ of 32.3% and a complex structure was refined using Phenix Refine (*83*), followed by manual rebuilding with the program COOT (*82*). The final refinement statistics, native data and phasing statistics are summarized in Table S9.

#### Co-crystallization of complex C57 Fab with S2_1.2

The complex of S2_1.2 with C57 Fab was prepared by mixing two components at a molar ratio of 4:1 for overnight incubation. The S2_1.2/Mota Fab complex was purified by size exclusion chromatography using a Superdex75 16 600 (GE Healthcare) equilibrated in 10 mM Tris pH 7.8, 50 mM NaCl and was subsequently concentrated to 10 mg/ml. Crystals were grown using sitting-drop vapor-diffusion method at 291K in a condition containing 20% (w/v) PEG 3350, 200 mM ammonium tartrate dibasic. For cryo protection, crystals were briefly swished through mother liquor containing 25% glycerol.

#### Data collection and structural determination of the S2_1.2/C57 Fab complex

Diffraction data were recorded with X06DA (PXIII) beamline at Paul Scherer Institute, Switzerland. The diffraction data were integrated and processed to 2.2 Å by XDS with a high-resolution cut at I/σ=1 applied. The crystals belonged to space group C121. The structure was determined by the molecular replacement using the Phaser module in the Phenix program (*81, 83*). The searching of the initial phase was performed by using the motavizumab structure (PDB 3IXT) and the computational model of S2_1.2 as a search model. Manual model building was performed using Coot (*82*), and automated refinement using Phenix Refine (*83*). The final refinement statistics are summarized in Table S10.

### Next-generation sequencing of design pools

After sorting, yeast cells were grown overnight, pelleted and plasmid DNA was extracted using Zymoprep Yeast Plasmid Miniprep II (Zymo Research) following the manufacturer’s instructions. The coding sequence of the designed variants was amplified using vector-specific primer pairs, Illumina sequencing adapters were attached using overhang PCR, and PCR products were desalted (Qiaquick PCR purification kit, Qiagen). Next generation sequencing was performed using an Illumina MiSeq 2×150 bp paired end sequencing (300 cycles), yielding between 0.45-0.58 million reads/sample.

For bioinformatic analysis, sequences were translated in the correct reading frame, and enrichment values were computed for each sequence. We defined the enrichment value E as follows:

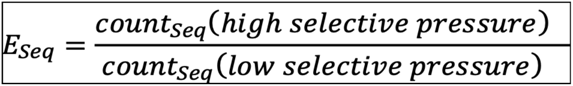

The high selective pressure corresponds to low labelling concentration of the respective target antibodies (100 pM D25, 10 nM 5C4 or 20 pM 101F), or a higher concentration of chymotrypsin protease (0.5 µM). The low selective pressure corresponds to a high labelling concentration with antibodies (10 nM D25, 1 µM 5C4 or 2 nM 101F), or without protease treatment. Only sequences that had at least one count in both sorting conditions were included in the analysis.

---

**Fig. S1.**
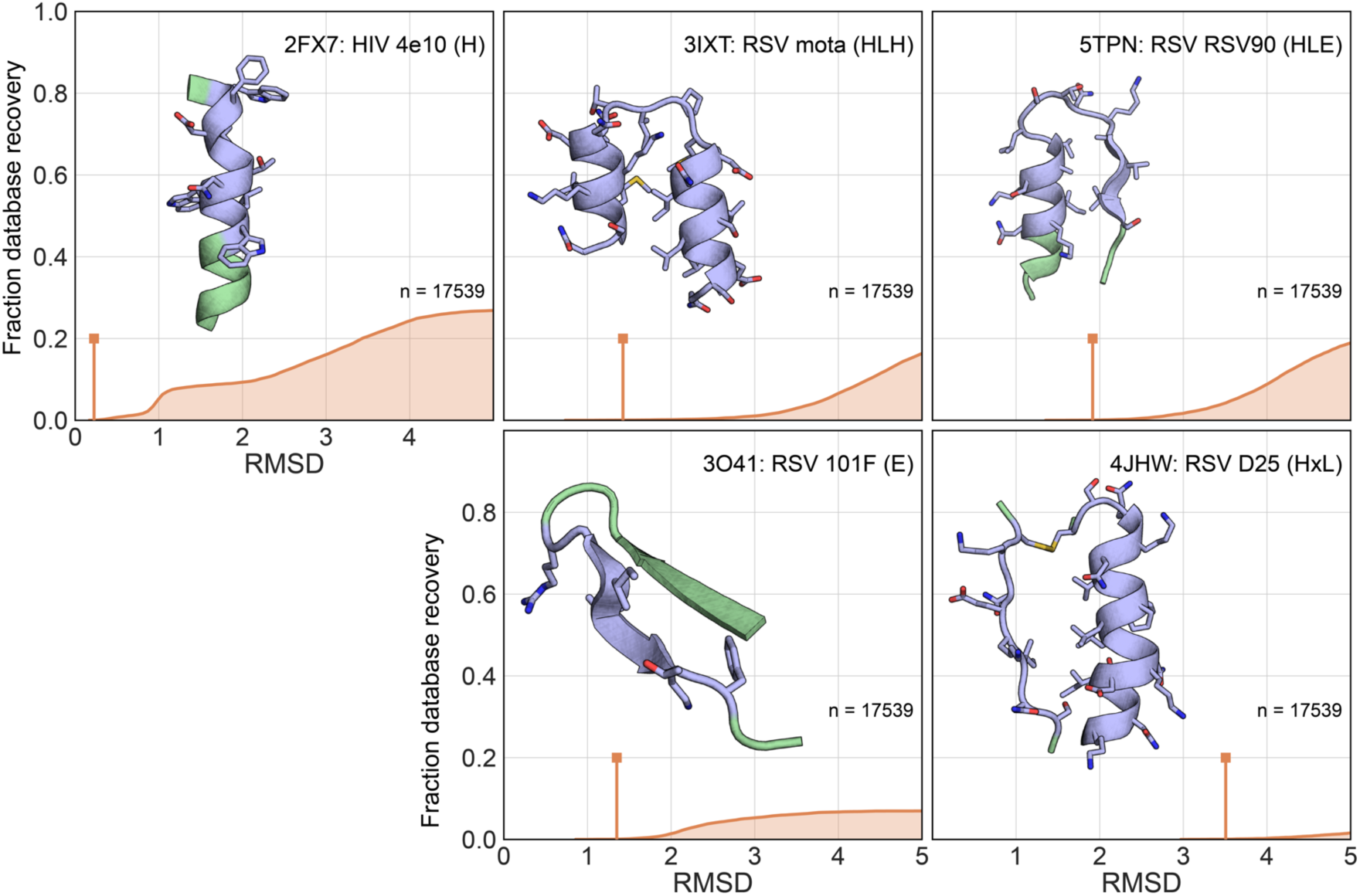
Number of natural template structures that can accommodate epitopes of different structural complexity. A MASTER search (*57*) was performed over the nrPDB30 database (nonredundant subset of the PDB with a 30% sequence identity cutoff) containing a total of 17,539 structures, querying the number of matches for different neutralization epitopes (colored in blue in the structures) of increasing structural complexity. The fraction of the database recovered is plotted on the y-axis. Matches were filtered for protein size <180 residues. The vertical line (orange) indicates the RMSD cutoff in Å for the first 10 scaffold identified. Secondary structure composition of the motifs is represented by: E - strand; L - Loop; H - helix; x - chain break.

**Fig. S2.**
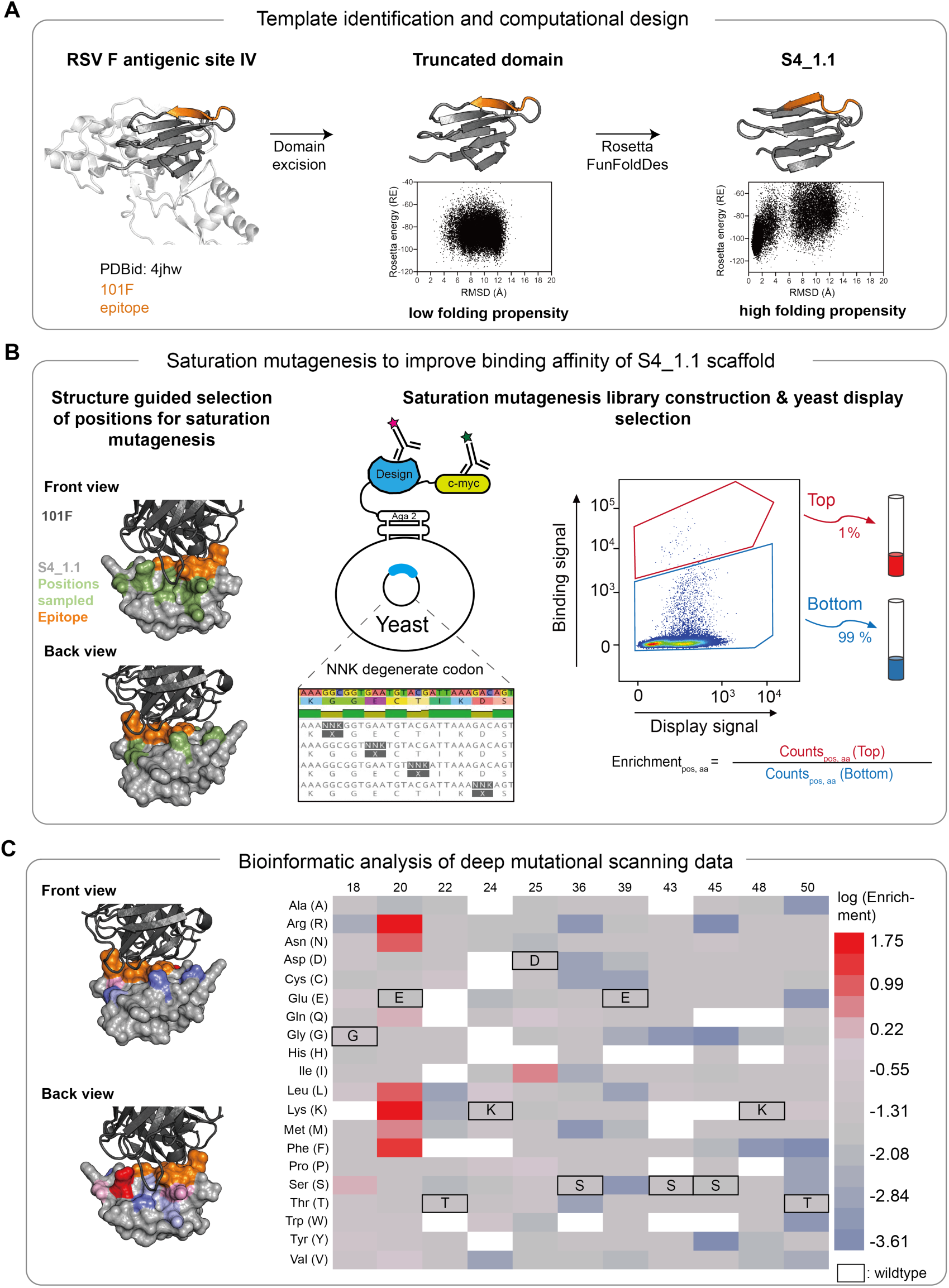
Computational design and experimental optimization of S4_1 design series. A) Template identification and computational design of S4_1.1. RSVF antigenic site IV is located in a small contained domain of preRSVF. This excised domain failed to show a folding funnel in Rosetta *abinitio* predictions, and failed to express recombinantly in E. coli. Using the excised domain as template, we folded and sequence-designed this topology using Rosetta FunFolDes, yielding design S4_1.1 which showed a strong funnel-shape energy landscape in *abinitio* folding simulations. B) Experimental optimization of S4_1.1 through saturation mutagenesis. A saturation mutagenesis library was constructed using overhang PCR for 11 positions proximal (green) to the site IV epitope (orange), allowing one position at a time to mutate to any of the 20 amino acids, encoded by the degenerate codon ‘NNK’. The library (size 11 positions x 32 codons = 352) was transformed in yeast, and designs were displayed on the cell surface. The selection was done by labeling the cells with 125 nM of 101F antibody. The top 1% of clones binding with high affinity to 101F antibody were then sorted, as well as the bottom 99% as shown. Following next-generation sequencing of the two populations, the enrichment values were computed for each sequence variant, corresponding to the relative abundance of each variant in the top versus bottom gate. C) Bioinformatic analysis of deep mutational scanning data. The log(Enrichment) is shown as heatmap (right) for each sequence variant, and mapped to the structure (left). White indicates missing data. Position 20 showed the highest enrichment for arginine and lysine, together with other less pronounced enrichments seen for other positions.

**Fig. S3.**
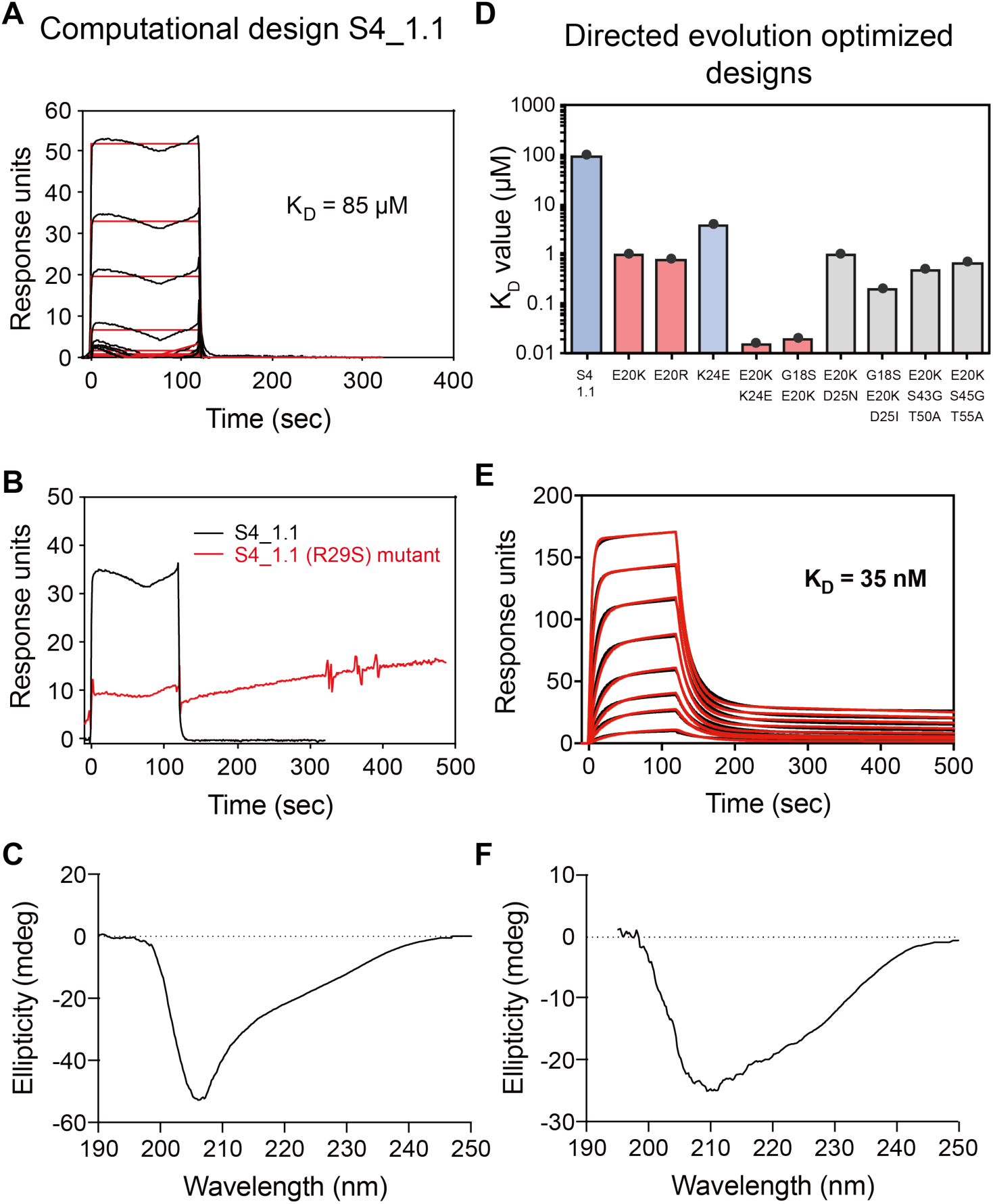
Experimental characterization of S4_1 design series. A) SPR measurement for the initial computational design S4_1.1 against 101F antibody revealed a dissociation constant (K_D_) higher than 85 µM. B) Despite low affinity, the R29S mutant showed that binding was specific to the epitope of interest. C) Circular dichroism spectrum of S4_1.1 at 20 °C. D) K_D_ for single and combined mutations of S4_1.1 that were identified in the deep mutational scanning screen. E20K/K24E double mutant (named S4_1.5) showed a K_D_ of 35 nM. E) SPR sensorgram of S4_1.5 against 101F. F) Circular dichroism spectrum of S4_1.5 at 20 °C.

**Fig. S4.**
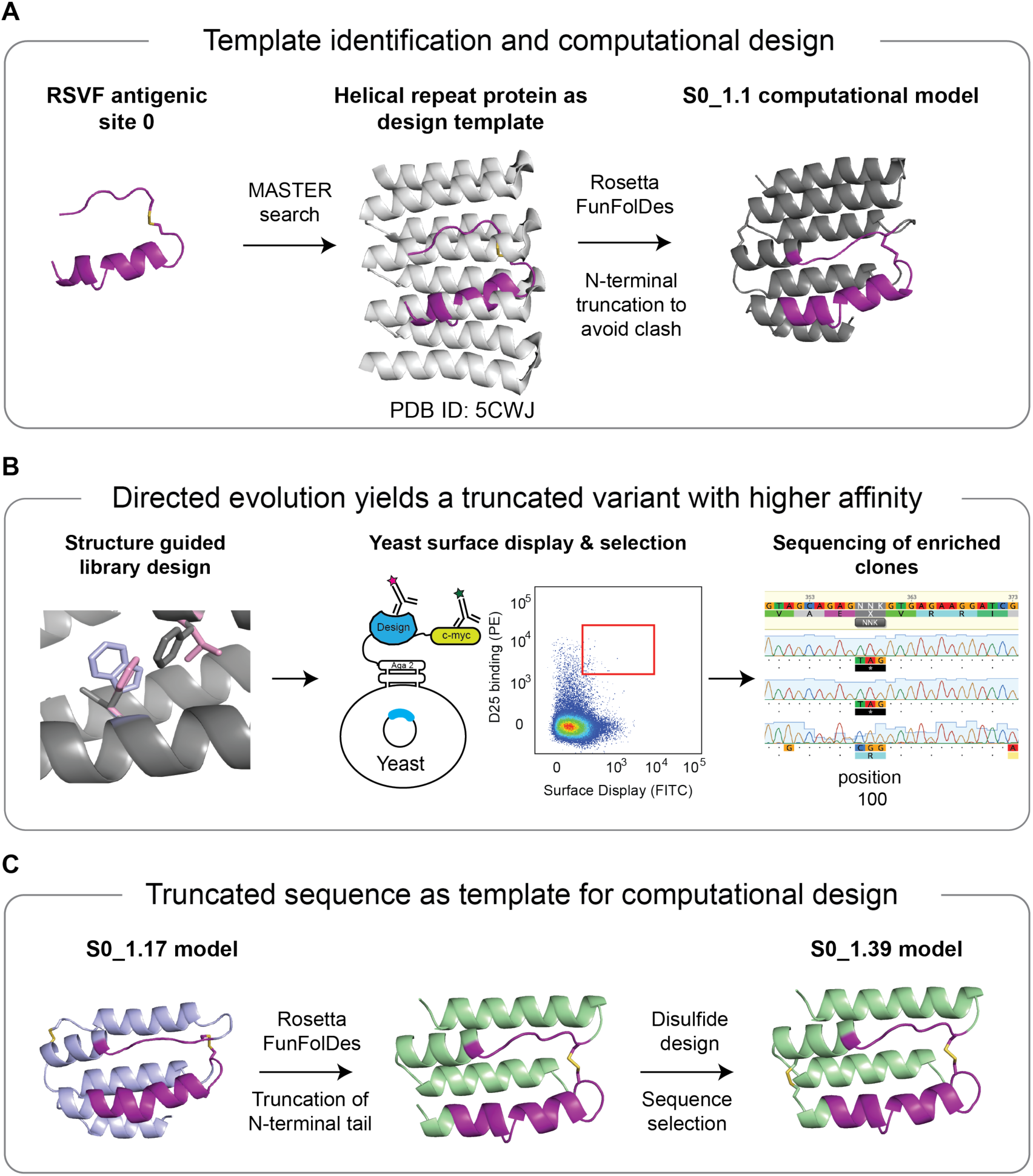
Computational design and experimental optimization of S0_1 design series. A) Template identification and design. Using MASTER, we identified a designed helical repeat protein (PDB 5CWJ) as design template to present and stabilize antigenic site 0 (see methods for details). The N-terminal 29 residues were truncated to avoid steric clashes with the D25 antibody, and Rosetta FunFolDes was used to design S0_1.1. B) Based on S0_1.1, a combinatorial sequence library was constructed and screened using yeast surface display. After three consecutive sorts of high-affinity binding clones, individual colonies were sequenced. Position 100 was frequently found to be mutated to a stop codon, leading to a truncated variant with increased expression yield, and a ∼5-fold improved binding affinity to D25 (Fig S5). C) A model of the truncated variant served as template for a second round of *in silico* folding and design. We further truncated the template by the N-terminal 14 residues, and introduced a disulfide bond between residues 1 and 43, leading to S0_1.39. See methods for full details on the design selection process.

**Fig. S5.**
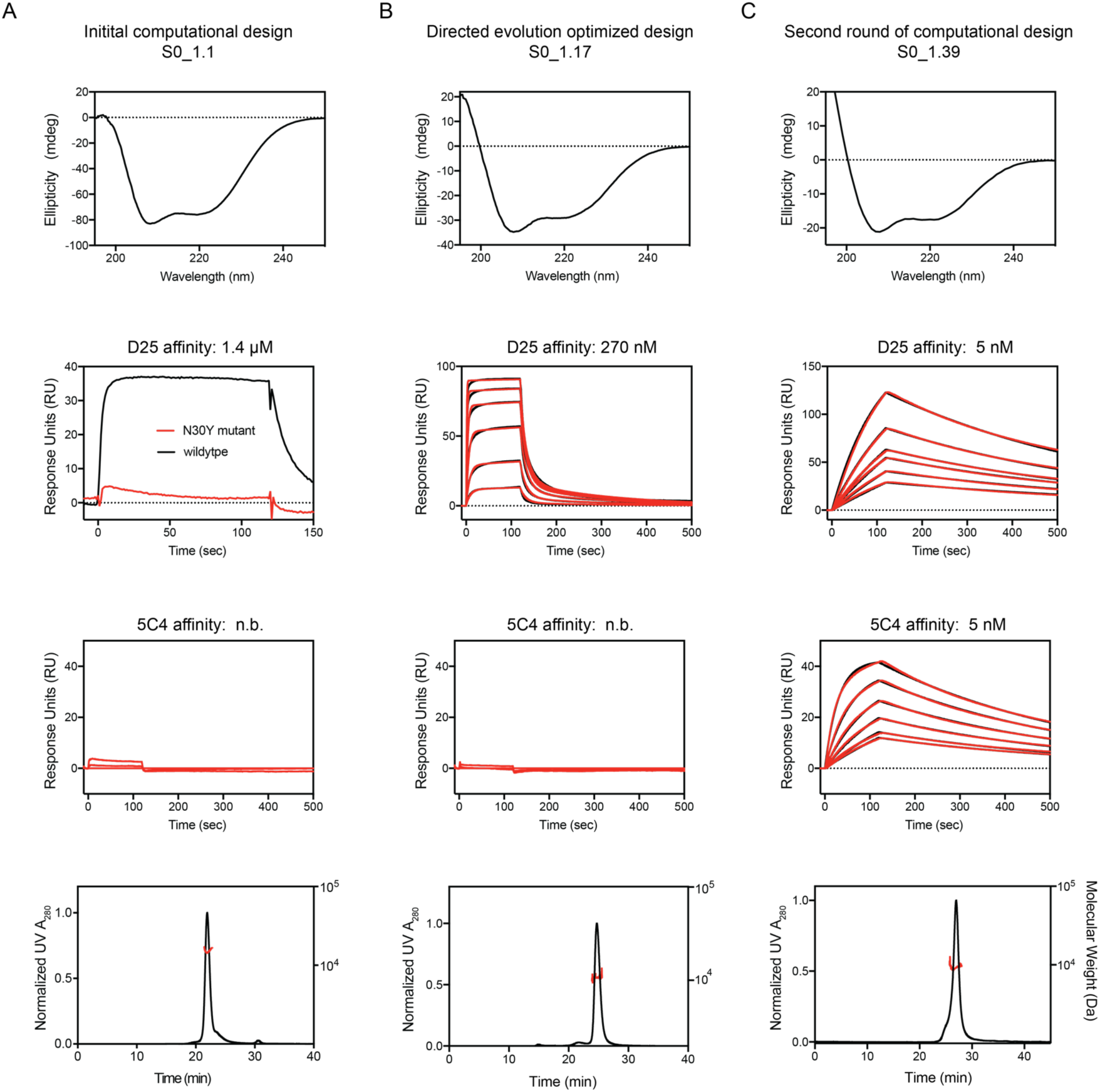
Biophysical characterization of the S0_1 design series. Top: Circular dichroism spectra at 20 °C. Middle: SPR measurements against D25 and 5C4. Bottom: Multi-angle light scattering coupled to size exclusion chromatography. A) S0_1.1 bound with a K_D_ of 1.4 µM to D25 and had no detectable binding to 5C4. To verify that the binding interaction was specific to the epitope we generated a knockout mutant (N30Y) and observed that the binding interaction was absent. B) S0_1.17 showed a K_D_ of 270 nM to D25 and no binding to 5C4. C) SPR sensorgrams of S0_1.39 binding to D25 and 5C4 antibodies. D25 or 5C4 IgG was immobilized as ligand on the sensor chip surface, and S0_1.39 was flown as analyte. All designs showed CD spectra typical of helical proteins and behaved as monomers is solution (top and bottom rows).

**Fig. S6.**
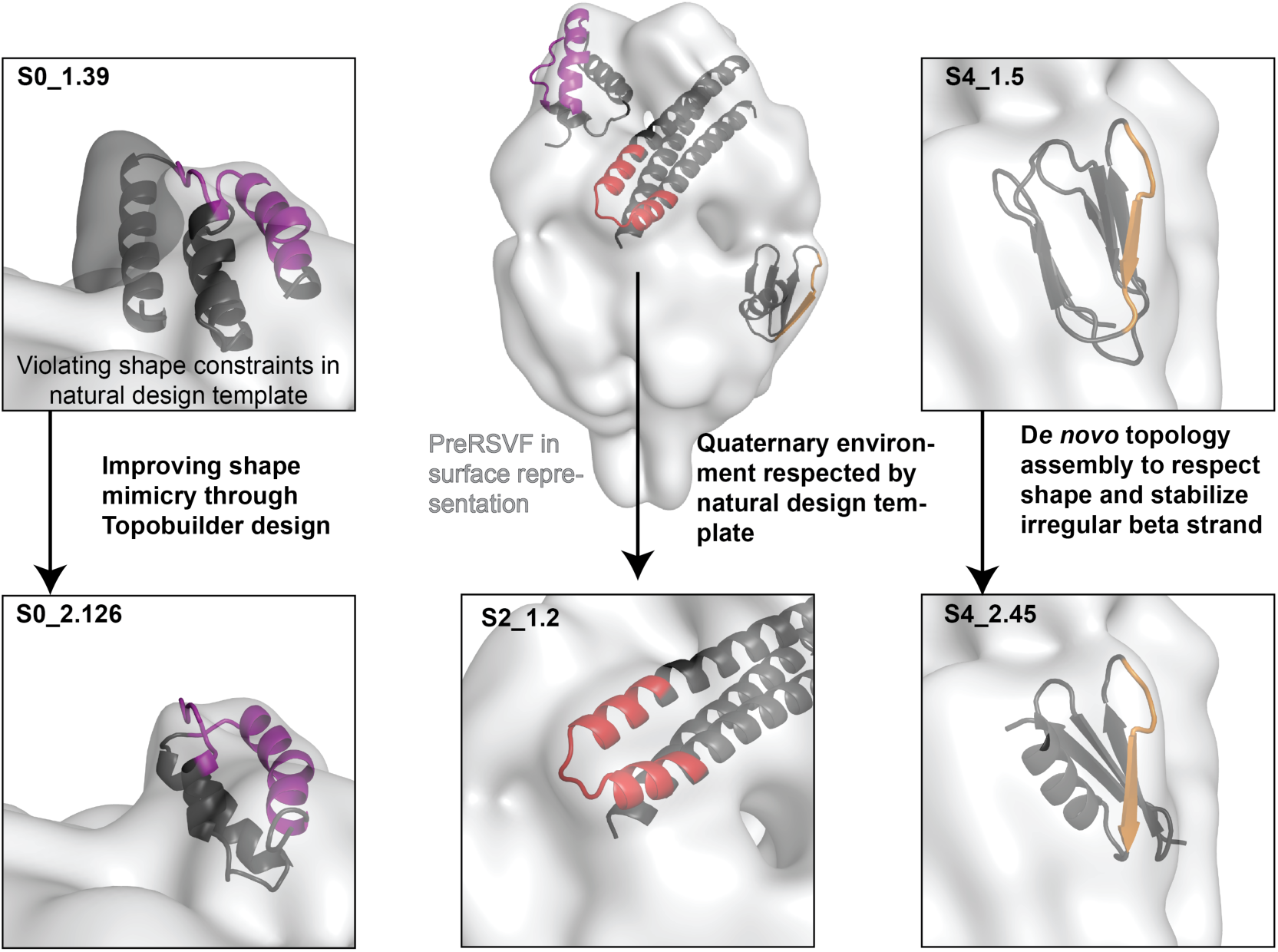
Shape mimicry of computationally designed immunogens compared to preRSVF. PreRSVF is shown in surface representation (light grey), with designed immunogens superimposed. Close-up views are shown for template-based designs (S0_1.39 and S4_1.5, top row). While site 0 is freely accessible for antibody binding in preRSVF, the C-terminal helix of S0_1.39 constrains its accessibility (dark grey surface). Through defined backbone assembly using TopoBuilder, S0_2.126 was designed, mimicking the native quaternary environment of site 0 (bottom left). RSVF antigenic site II, which is a structurally simple helix-turn-helix motif frequently found in natural proteins, was previously designed based on a design template that respects the quaternary constraints of site II in its native environment (S2_1.2, bottom middle). For site IV, a topology was assembled (S4_2.45) that respects the shape constraints while improving the stabilization of the irregular, bulged beta strand compared to the S4_1.5 design (right).

**Fig. S7.**
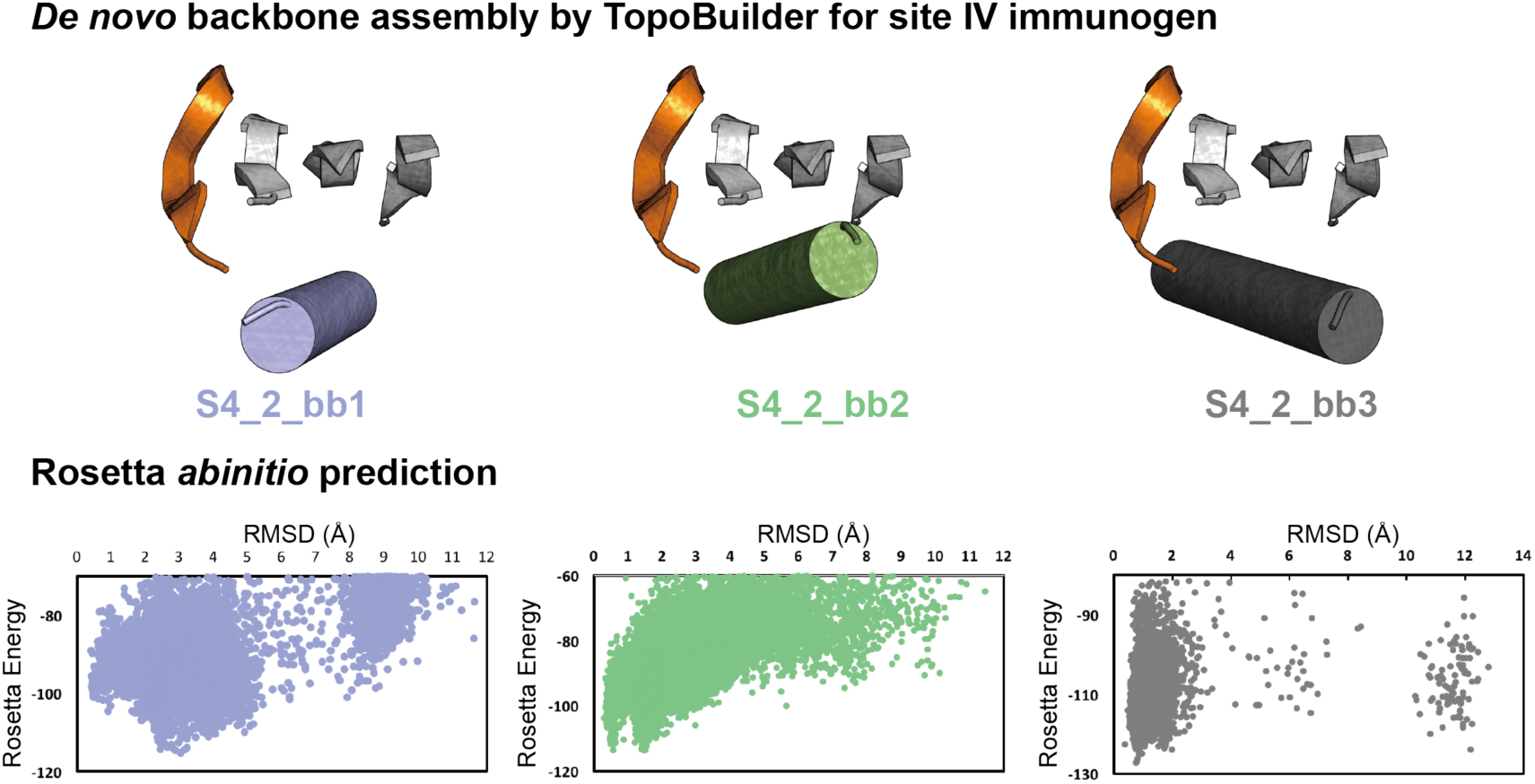
D*e novo* backbone assembly for site IV immunogen. The site IV epitope was stabilized with three antiparallel beta strands built *de novo*, and a helix packing in various orientations against this beta sheet (bb1-bb3). Rosetta *abinitio* simulations were performed to evaluate the ability of the sequences generated in the different backbones to fold into a low energy state close to the design model, indicating that S4_2_bb2 and bb3 have a stronger tendency to converge into the designed fold.

**Fig S8.**
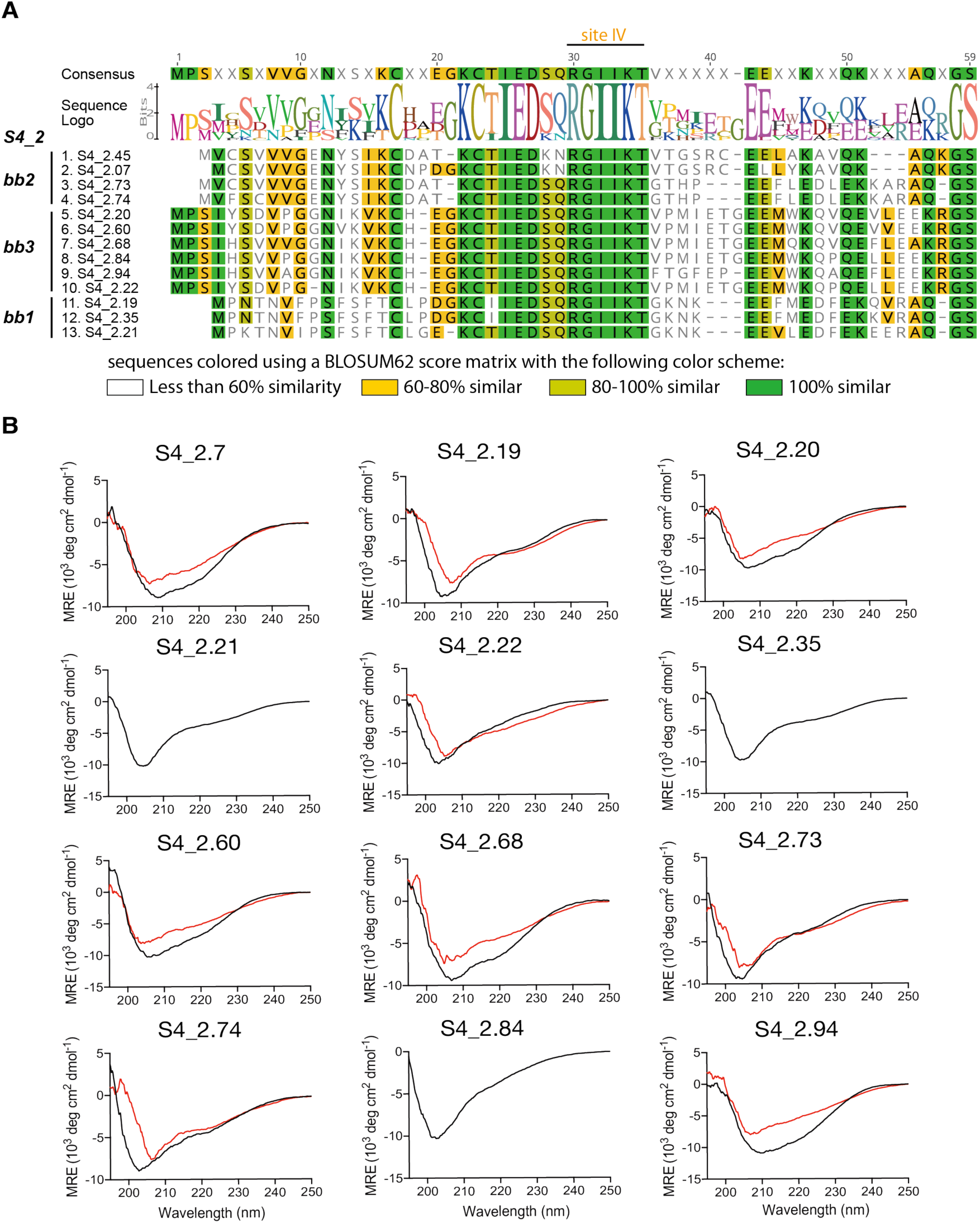
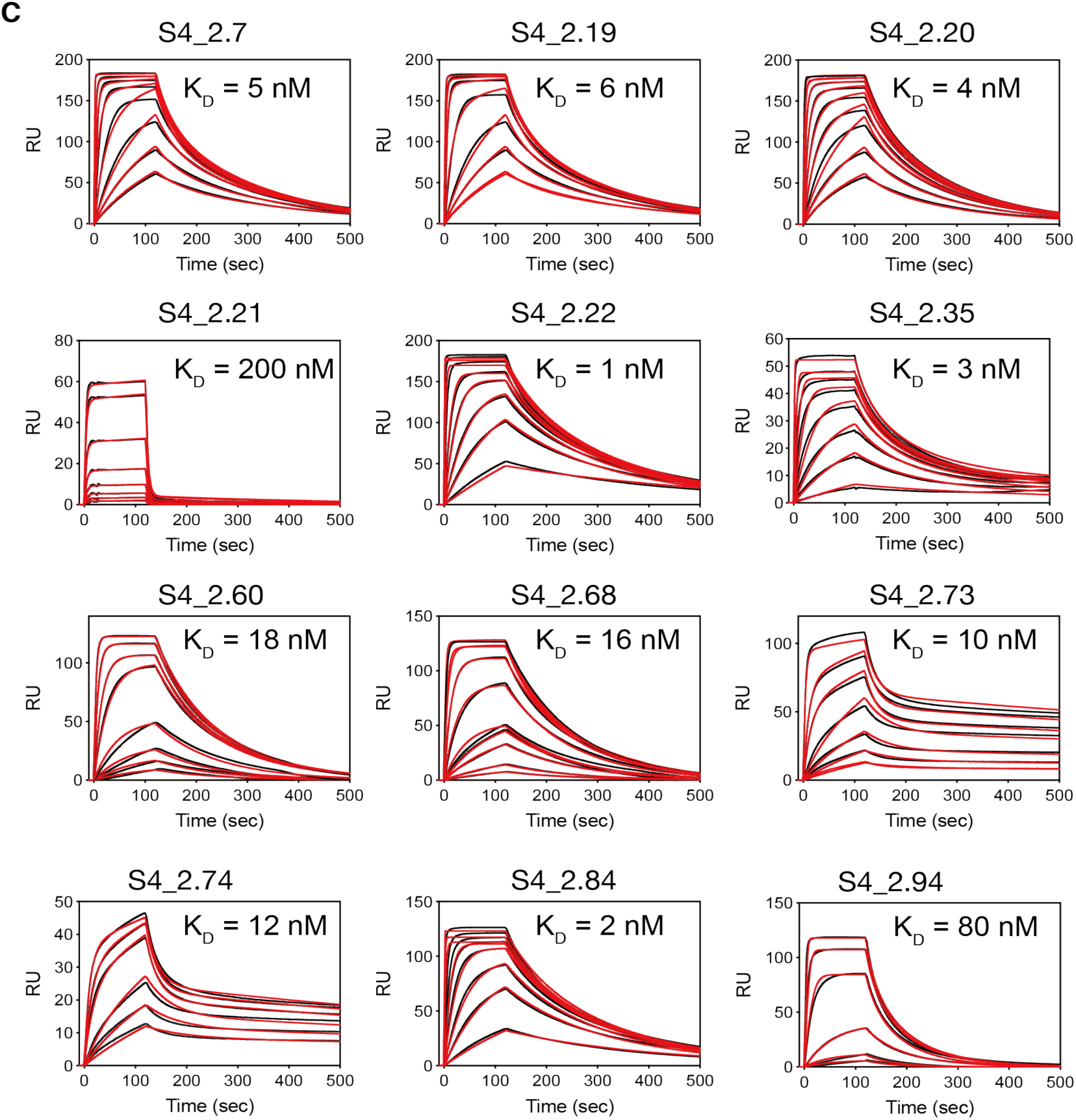
Biophysical characterization of S4_2 design series. A) Multiple sequence alignment of S4_2 sequences that were enriched after yeast screening for protease resistance and binding to 101F. Sequences are colored according to a BLOSUM62 score matrix. B) Circular dichroism spectra for 12 designs of the S4_2 design series that were enriched for protease resistance and binding to 101F in the yeast display selection assay. Black: spectrum at 20 °C. Red: spectrum at 90 °C. C) SPR sensorgrams for binding to 101F for the same designs. 101F IgG was immobilized on the CM5 sensor chip surface, and the designs were flown as analyte.

**Fig S9.**
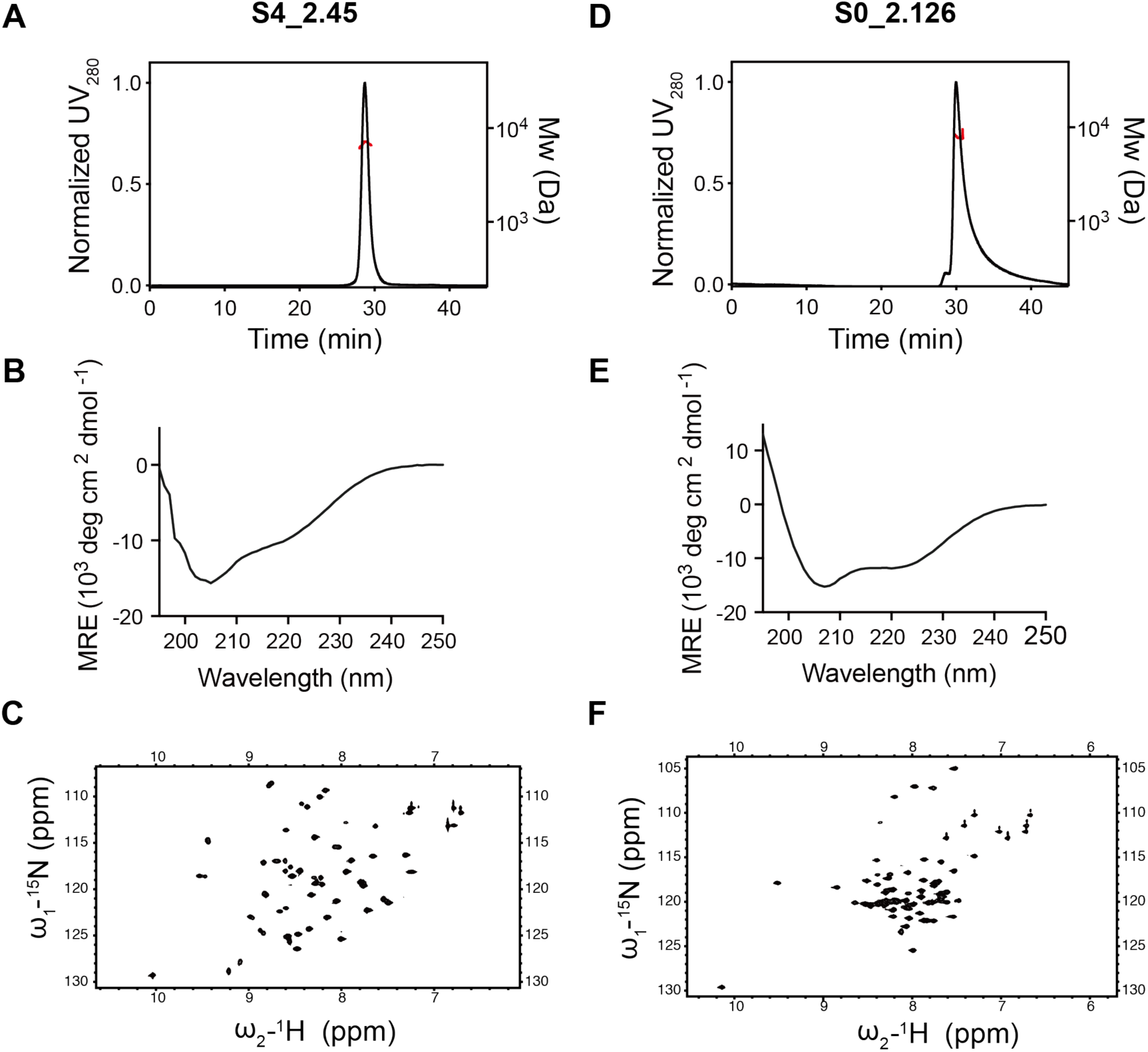
Biophysical characterization of the designs S4_2.45 and S0_2.126. A,D) Designs are monomeric in solution as shown by SEC-MALS profile. B,E) Circular dichroism spectra at 25 °C. C,F) 2D NMR of ^15^N HSQC spectra are well dispersed, confirming that the designs are well folded in solution. See zoom-in for S0_2.126 spectrum with its assignment on following page. Mw: Molecular Weight.

**Fig. S10.**
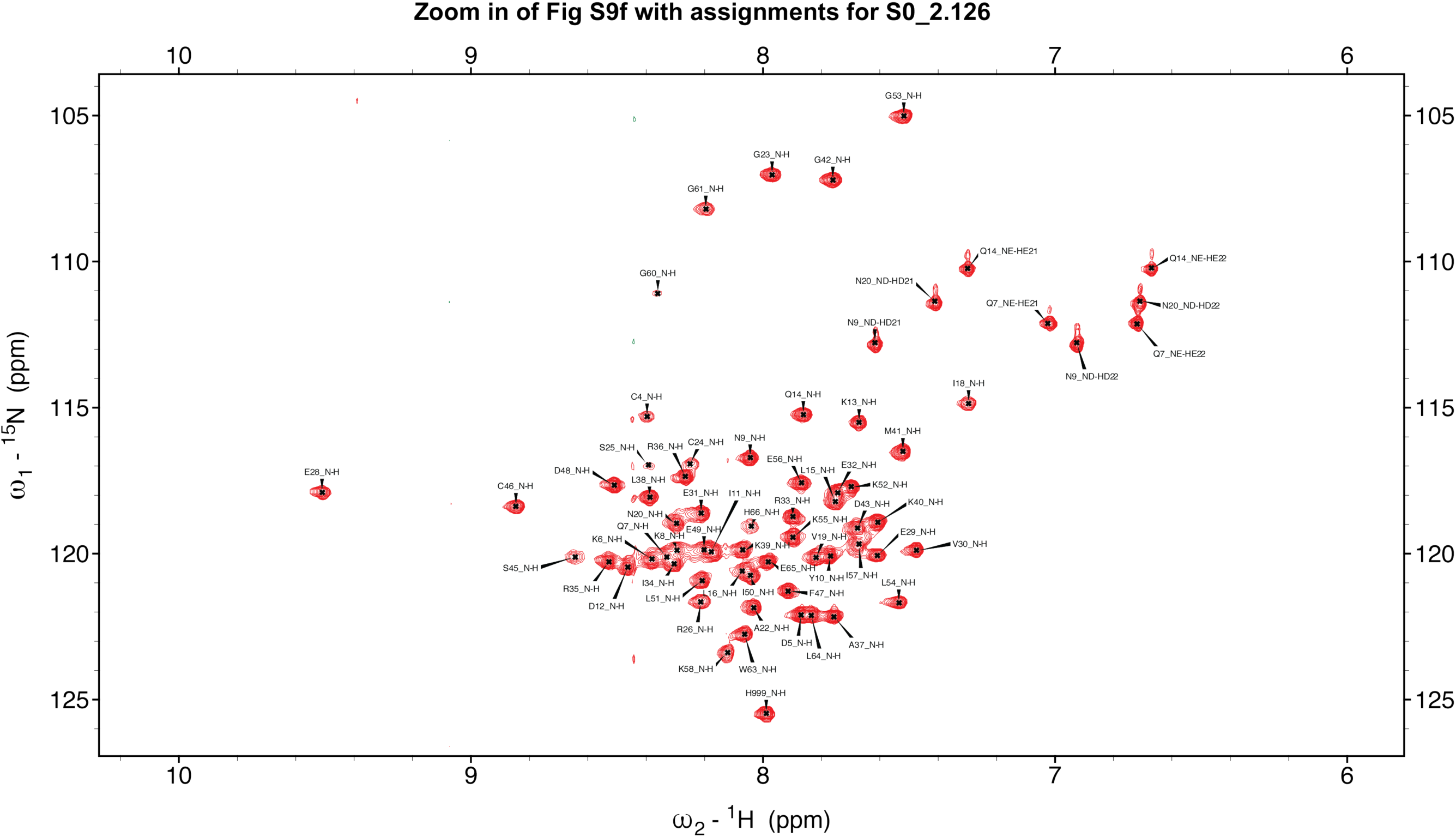
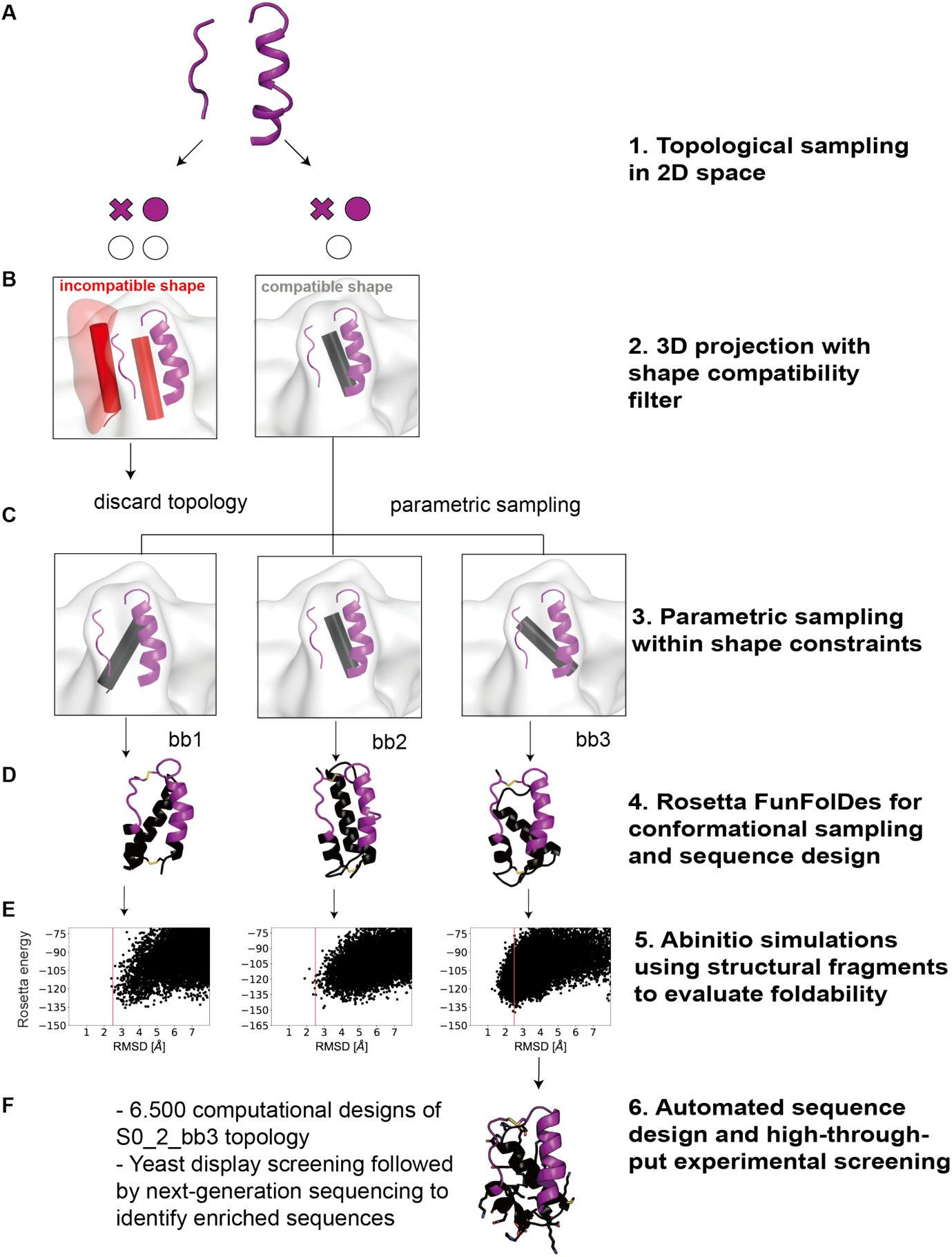
D*e novo* topology assembly to stabilize site 0 using TopoBuilder. A) Topologies that allow to connect the discontinuous site 0 motif are built as 2D forms, and then translated into 3D sketches. B) Generated sketches are evaluated for their compatibility with the shape constraints of preRSVF. A compatible shape (right) respects the quaternary constraints of the motif in its native environment, whereas an incompatible shape (left) is one that sterically constrains the motif’s accessibility in a non-native manner (red surface). C) Once a compatible topology is identified, the secondary structure orientation is parametrically sampled (3 different helical orientations, S0_2_bb1-bb3). D) Three customized helical orientations were assembled (S0_2_bb1-bb3) to support site 0 epitope, and evaluated for their ability to fold into the designed topology by Rosetta *abinitio* simulations. E) S0_2_bb3-based designs showed a funnel-shaped energy landscape. F) S0_2_bb3 backbone was selected for automated sequence design and experimental screening.

**Fig. S11.**
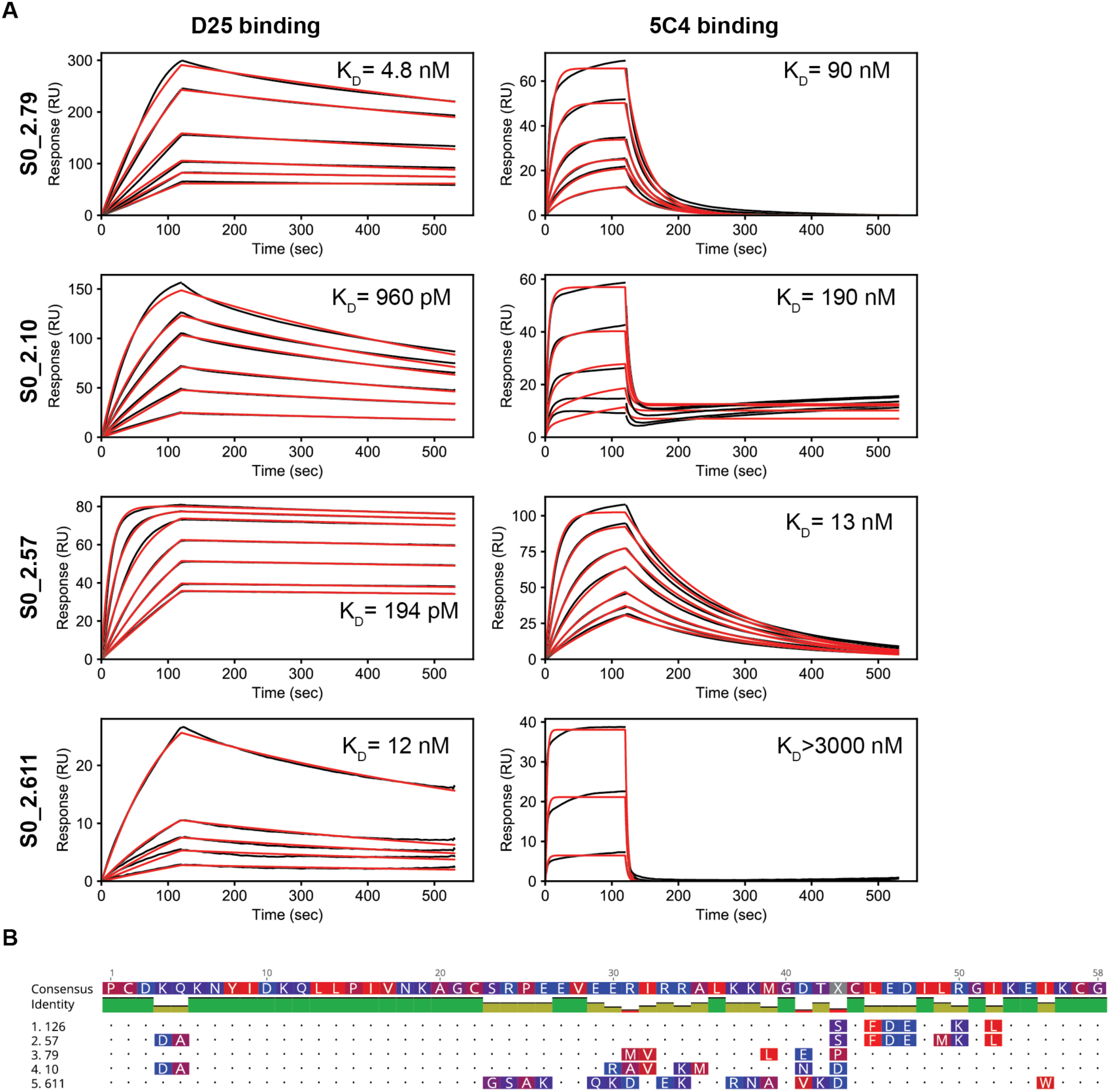
Biophysical characterization of S0_2 design series. A) SPR affinity measurement of *de novo* site 0 scaffolds against D25 or 5C4. IgGs were immobilized on the sensor chip surface, and scaffolds were injected as analyte. Dissociation constants shown are kinetic fits using a 1:1 Langmuir model. B) Sequence alignment of experimentally characterized sequences, in comparison to S0_2.126. The closest sequence homolog to S0_2.126 is S0_2.57, differing in 3 amino acids.

**Fig. S12.**
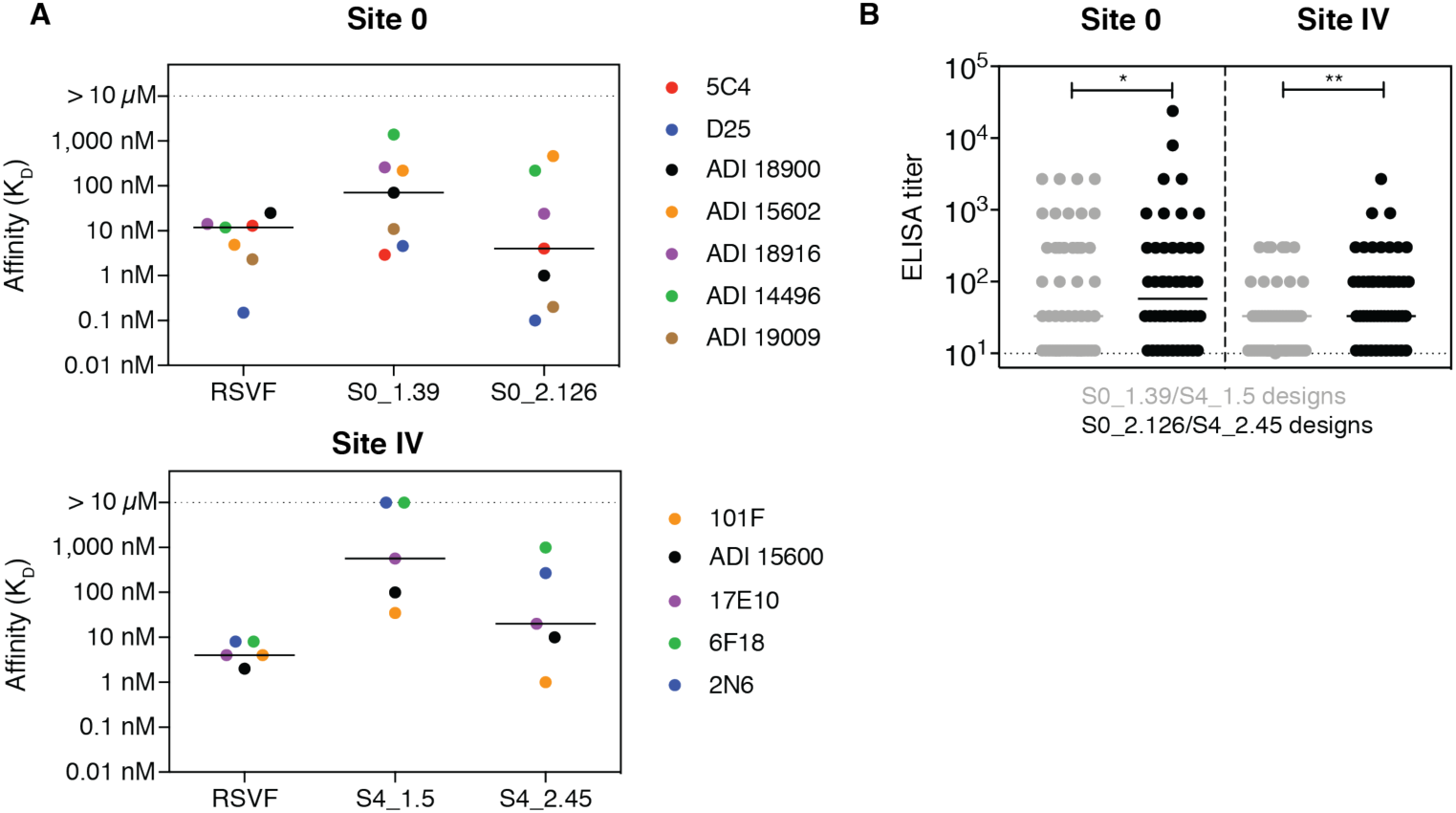
Binding affinity of designed immunogens towards panels of site-specific, human neutralizing antibodies and human sera. A) Top: Binding affinity (K_D_, determined by SPR flowing Fabs as analyte) of S0_1.39 and S0_2.126 towards a panel of site-specific neutralizing antibodies, in comparison to preRSVF. Antibodies shown for site 0: 5C4, D25 (*30*), ADI-14496, ADI-18916, ADI-15602, ADI-18900 and ADI-19009 (*41*). Bottom: For site IV, the binding affinity was tested against: 101F (*31*), ADI-15600 (*41*), 17E10, 6F18 and 2N6 (*85*). A comparison between S4_1.5 and S4_2.45 to preRSVF is shown. The higher binding affinity of the second-generation designs (S0_2.126 and S4_2.45) compared to the first-generation indicates a greatly improved, near-native epitope mimicry of the respective antigenic sites in the designed immunogens. B) ELISA reactivity of designed immunogens with sera obtained from 50 healthy human adults that were seropositive for preRSVF. Both S0_2.126 and S4_2.45 showed significantly increased reactivity compared to the first-generation designs, confirming an improved epitope-mimicry on the serum level (* p< 0.05 and ** p< 0.01, Wilcoxon test). Data are representative from one out of two independent experiments.

**Fig. S13.**
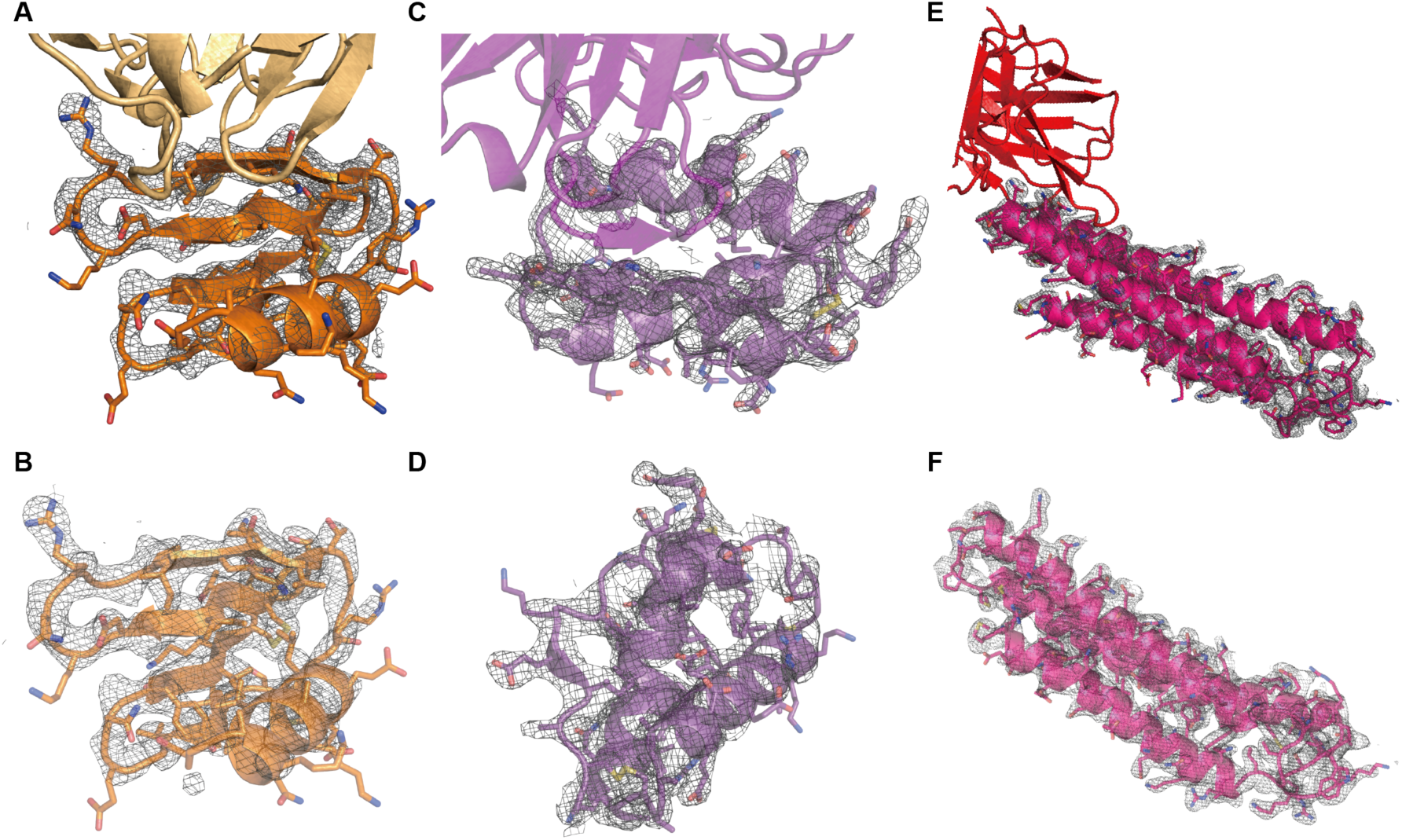
2mFo-mF_c_ electron density maps of the presented x-ray structures. Maps are contoured at 1 σ and carved around the scaffold at 1.6 Å. Side chains are shown in stick representation. A) S4_2.45 bound to 101F Fab. B) S4_2.45 scaffold only. C) S0_2.126 bound to D25 Fab. D) S0_2.126 scaffold. E) S2_1.2 scaffold bound to the Fab of the NHP elicited antibody C57. F) S2_1.2 scaffold only.

**Fig. S14.**
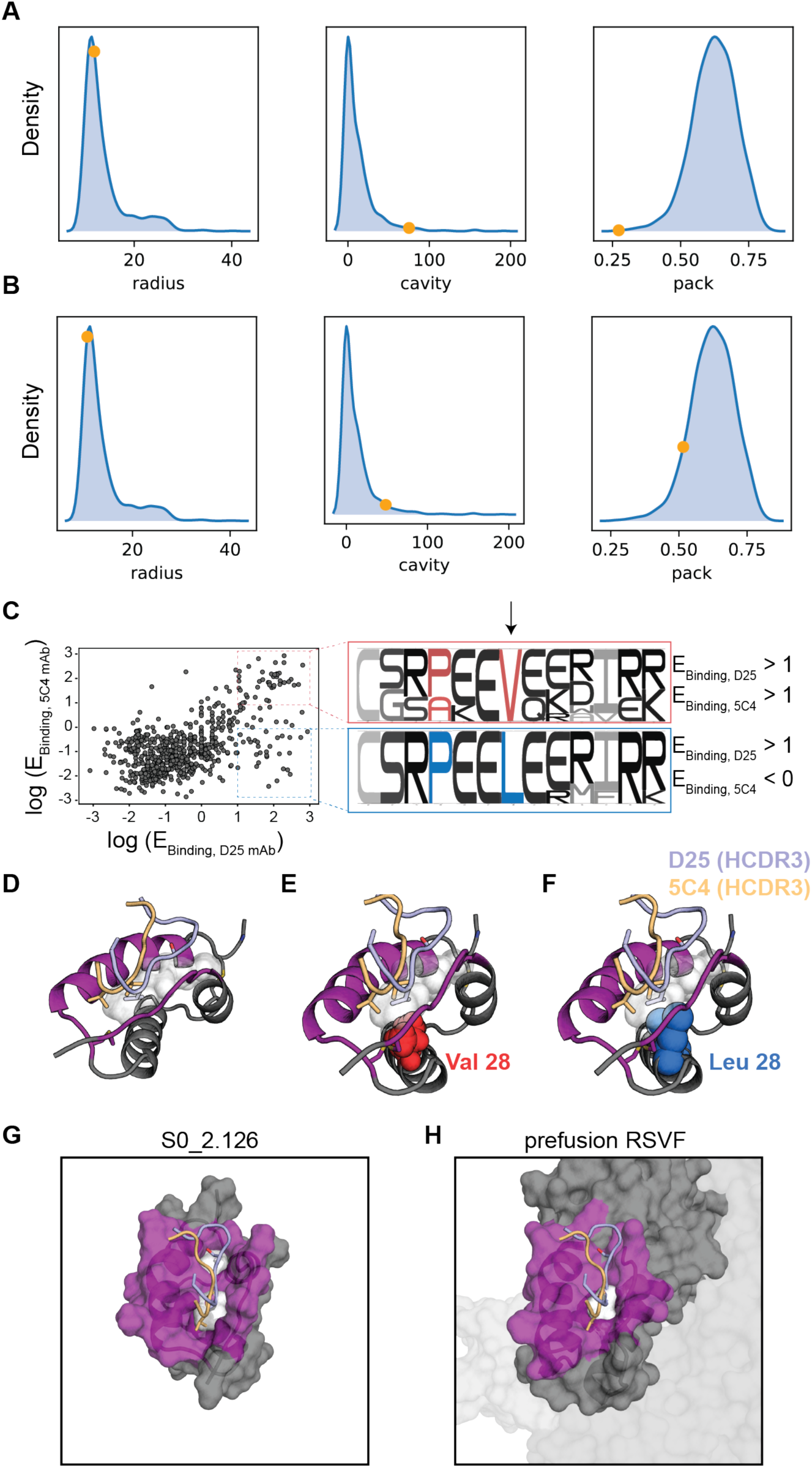
S0_2.126 maintains a large core cavity that is rarely found in natural proteins of similar size. A-B) Comparison of S0_2.126 Rosetta scores and structural metrics against natural proteins of similar size. Protein structures within the same size as S0_2.126 (57 +/- 5 residues) were downloaded from the CATH database and filtered by 70% sequence homology, yielding a representative database of natural proteins with similar size as S0_2.126 (n = 1,013 structures). Proteins were then minimized and scored by Rosetta to compute their radius of gyration (radius), intra-protein cavities (cavity) and core packing (packstat). The plots show the distribution for the metrics across 1,013 natural proteins (blue density plot), S0_2.126 is shown with the orange dot. The NMR structure of and computational model of S0_2.126 are shown in (A) and (B), respectively. The distributions show that, despite similar radius of gyration, S0_2.126 has a substantial cavity volume as well as very low core packing compared to natural proteins of similar size. CATH database and scores were pre-calculated, loaded and visualized using the rstoolbox python library (*86*). C) Next-generation sequencing of S0_2.126 library (as shown in Fig 3). Sequences with strong enrichment for both 5C4 and D25 binding have a valine in position 28 (red), whereas sequences enriched for D25 only have a leucine in the same position (blue). D) Model of S0_2.126 (grey, site 0 highlighted in purple). A binding cleft (white surface) between the helical and the loop segment of site 0 is engaged by the heavy chain CDR3 of two structurally characterized site 0 nAbs (D25/5C4). E-F) Residue 28 is located in a key position to form the D25/5C4 binding cleft, a valine allows binding of both D25 and 5C4 whereas a leucine in the same position abrogates binding to 5C4. G-H) Comparison of the binding cleft between S0_2.126 (G) and preRSVF (H), illustrating that it has been accurately preserved in the designed scaffold.

**Fig. S15.**
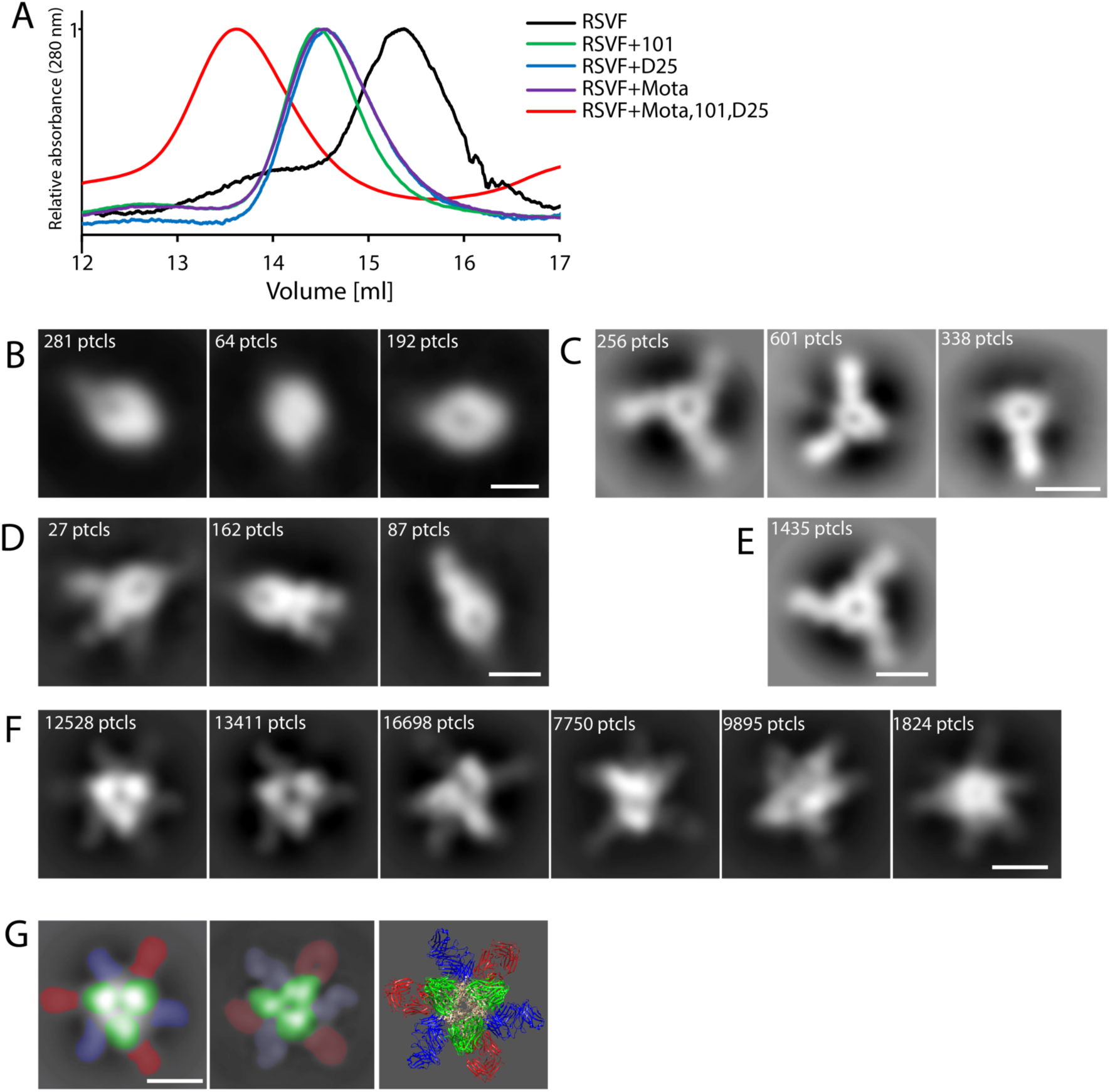
Electron microscopy analysis of site-specific antibodies in complex with RSVF trimer. A) Superposed size-exclusion profiles of unliganded RSVF (black line) and RSVF in complex with 101F (green line), D25 (blue line), Mota (purple line) and all three (101F, D25, Mota - red line) Fabs. B-F) Representative reference-free 2D class averages of the unliganded RSVF trimer (B) and RSVF in complex with 101F (C), D25 (D), Mota (E) or all three (F) Fabs. Fully-saturated RSVF trimers bound by Fabs are observed, as well as sub-stoichiometric classes. G) Left panel: reference-free 2D class average of RSVF trimer with three copies of 101F, D25 and Mota Fabs visibly bound. The predicted structure of RSVF in complex with 101F, D25 and Mota was used to simulate 2D class averages in Cryosparc2, and simulated 2D class average with all three types of Fabs is shown in the middle panel. Right panel: predicted structure of RSVF trimer with bound 101F, D25 and Mota Fabs based on the existing structures of RSVF with individual Fabs (PDB 4JHW, 3QWO and 3O45). Fabs are colored as follow: red - 101F; blue - Mota; green - D25. Scale bar - 100 Å.

**Fig. S16.**
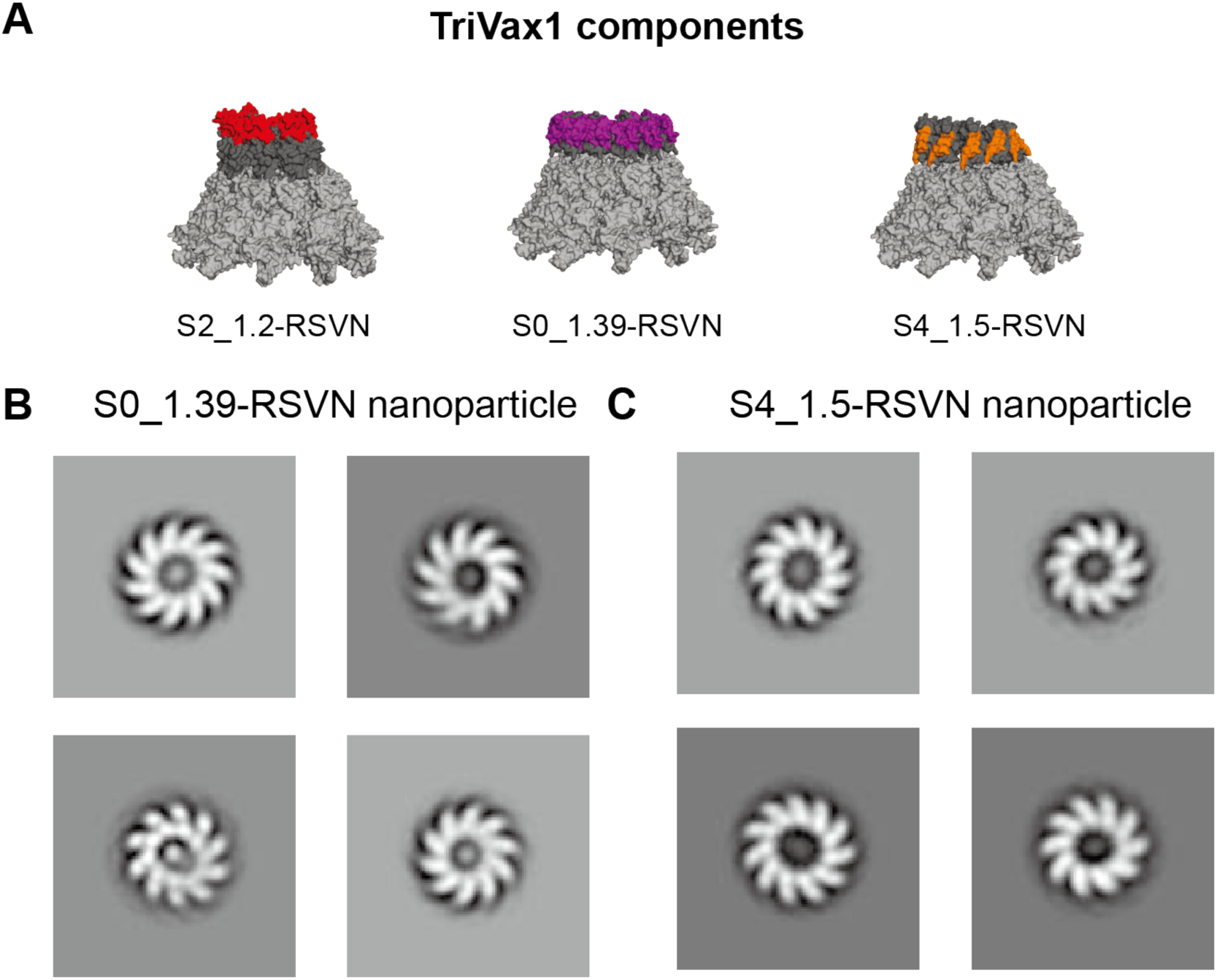
Composition and EM analysis of Trivax1 RSVN nanoparticles. A) Trivax1 contains equimolar amounts of site II, 0 and IV epitope focused immunogens fused to the self-assembling RSVN nanoparticle with a ring-like structure (n = 10-11 subunits). The site II-RSVN nanoparticle has been described previously (*14*). Surface representation of the computational models for the nanoparticles-immunogen fusion proteins with the epitopes highlighted. B,C) Negative stain electron microscopy for S0_1.39-RSVN and S4_1.5-RSVN nanoparticles confirms that the ring-like structure is maintained upon fusion of the designed immunogens.

**Fig. S17.**
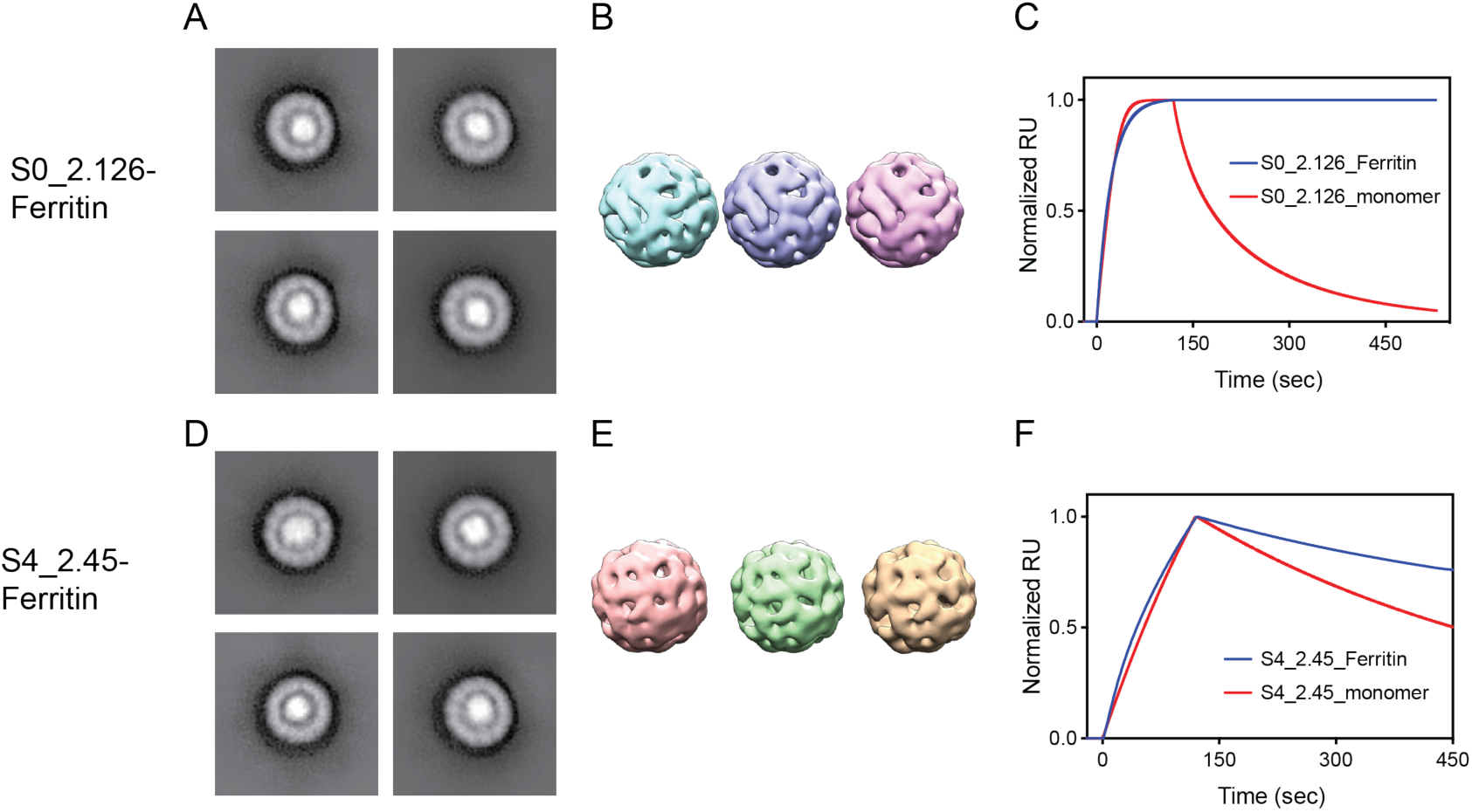
EM analysis of Trivax2 ferritin nanoparticles. A,B,D,E) Negative stain electron microscopy (A,D) and 3D reconstruction (B,E) for S0_2.126 and S4_2.45 fused to ferritin nanoparticles. C) Binding affinity of S0_2.126 nanoparticle (blue) to 5C4 antibody in comparison to S0_2.126 monomer (red), showing that S0_2.126 has been successfully multimerized and antibody binding sites are accessible. F) Binding of S4_2.45 to 101F antibody when multimerized on ferritin nanoparticle (blue) compared to monomeric S4_2.45 (red), indicating that the scaffold is multimerized and the epitope is accessible for antibody binding.

**Fig. S18.**
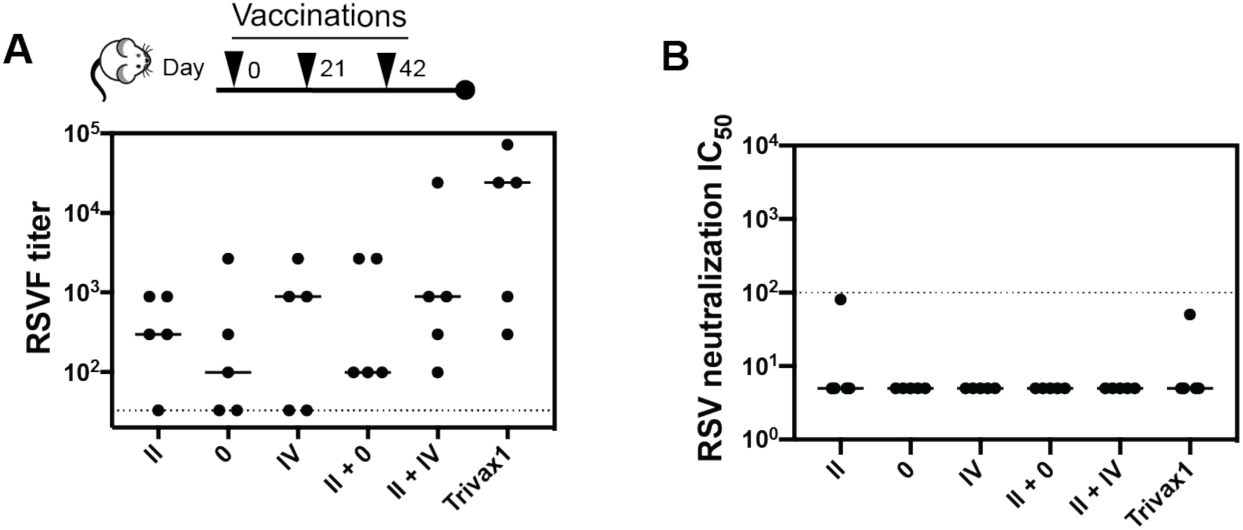
Mouse immunization studies with Trivax1. A) RSVF cross-reactivity of epitope-focused immunogens formulated individually, as cocktail of two, and three (Trivax1). B) RSV neutralizing serum titers of mice immunized with designed immunogens and combinations thereof.

**Fig. S19.**
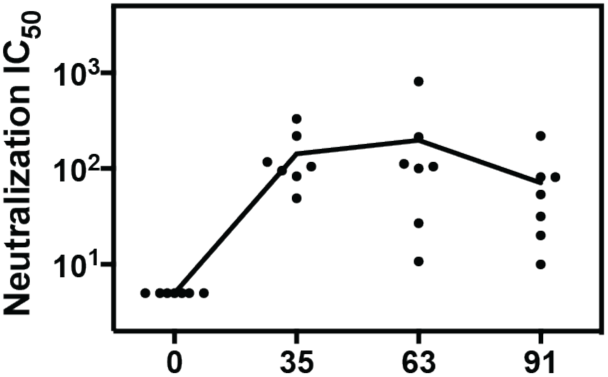
Confirmation of NHP neutralization titer by an independent laboratory. Sera from indicated timepoints were tested for RSV neutralization by an independent laboratory in a different RSV neutralization assay, using a Vero-118 cell line and a GFP readout. See (*87*) for method details.

**Fig. S20.**
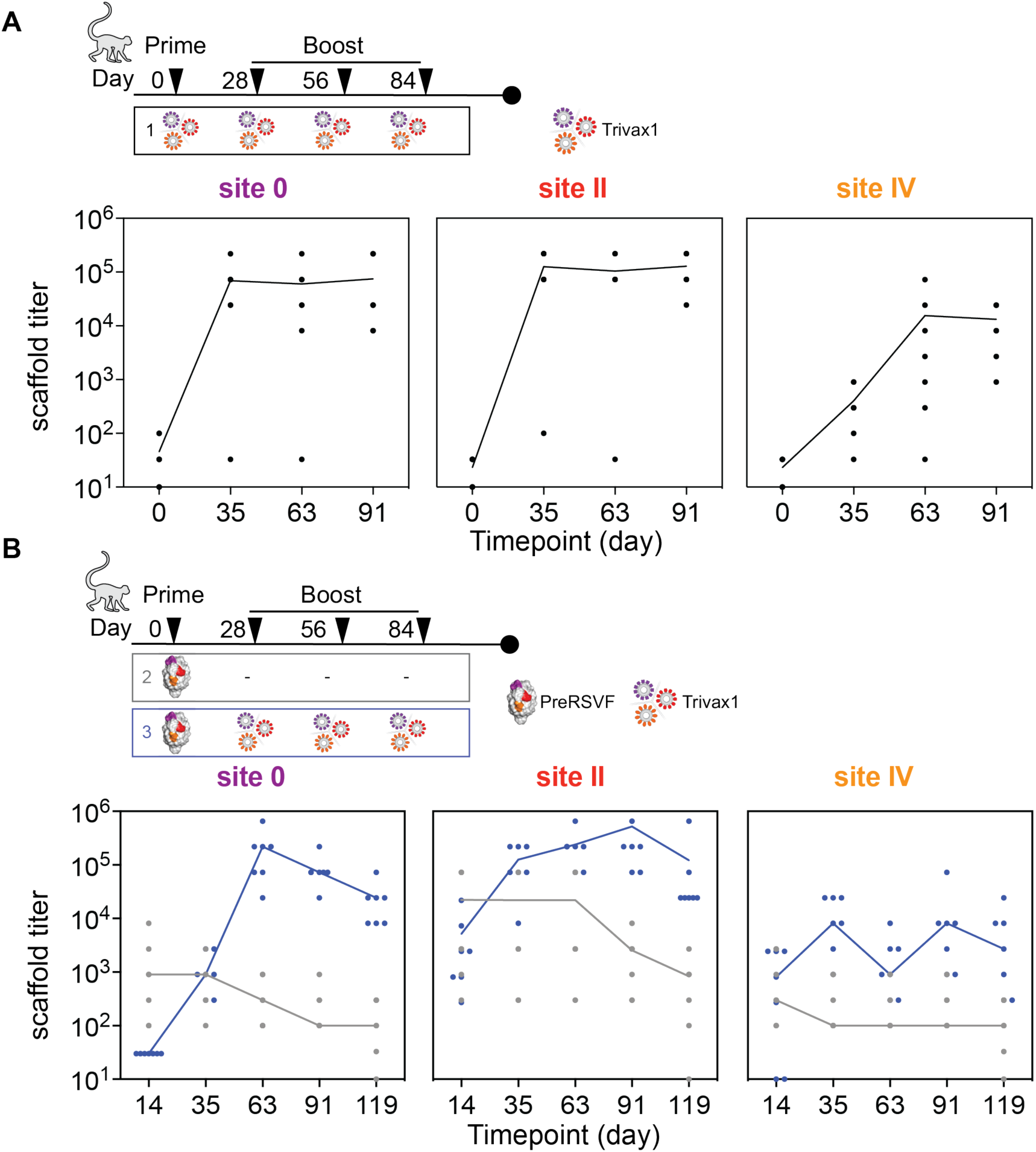
NHP serum reactivity with designed immunogens. A) ELISA titer of NHP group 1 (immunized with Trivax1) measured at different timepoints. All animals responded to Trivax1 immunogens at day 91, with site IV immunogen reactivity lower compared to site 0 and site II reactivity. B) ELISA titer of NHP group 2 (grey, preRSVF prime) and 3 (blue, preRSVF prime, Trivax1 boost). Following the priming immunization, all animals developed detectable cross-reactivity with the designed immunogens, indicating that the designed scaffolds recognized relevant antibodies primed by preRSVF.

**Fig. S21.**
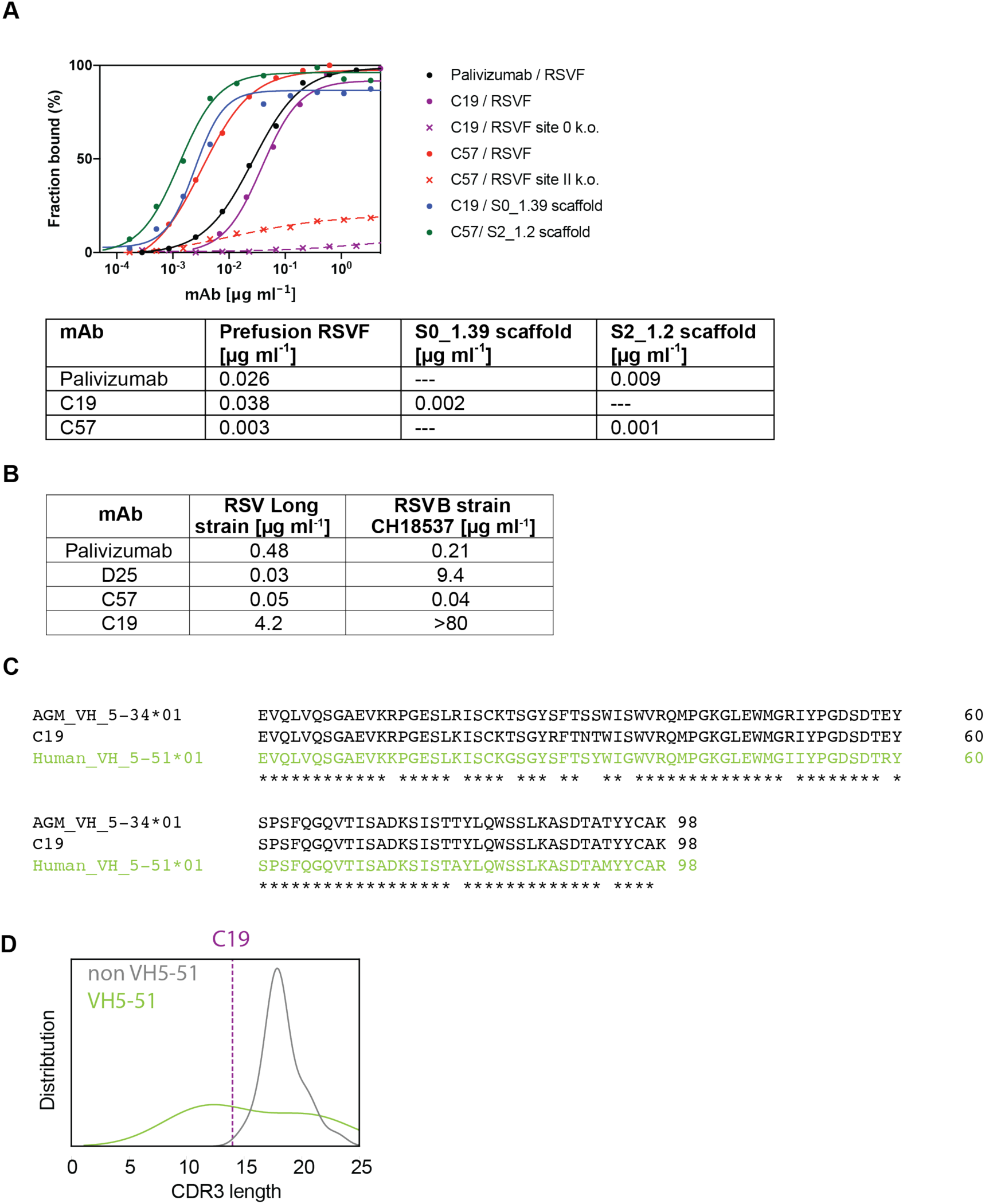
Monoclonal antibody characterization. A) ELISA binding curves of Palivizumab and elicited mAbs C57 and C19. PreRSVF, site-specific knockout mutations of preRSVF (site 0: K201E, N208R, N63R, K68E; site II: N262R, K272E, S275R) (*41*) or the designed scaffolds were coated on the plate, and IgGs were added in serial dilutions (performed in triplicates). ELISA signal was normalized from 0 to 1 and curves were fitted using GraphPad Prism. IC_50_ values are given in µg/ml. B) *In vitro* RSV neutralization potency of mAbs (IC_50_ in µg/ml) of RSV Long strain and B strain CH18537, performed in duplicates. C) Sequence alignment of African green monkey (AGM) VH5-34*01 gene, the site 0 specific C19 antibody, and the closest human homolog VH5-51*01. VH sequence of C19 was blasted against a database of AGM heavy chain germlines genes described previously (*88*), revealing the VH5-34*01 lineage as the inferred germline precursor of C19 (98.3% identity). The closest human homolog to AGM VH5-34*01 is the VH5-51*01 gene (94.1% homology) (*88*). D) CDR3 length distribution of known human site 0 nAbs (*41*). All antibodies that do not derive from VH5-51 (total of 25 antibodies deriving from 10 different VH lineages) show a CDR3 length, with a median of 18 amino acids. In contrast, VH5-51 antibodies (5 different antibodies derived from 3 different donors) show CDR3 lengths with a median of 14 amino acids, identical to the CDR3 length of C19 (dashed line).

**Table S1.**
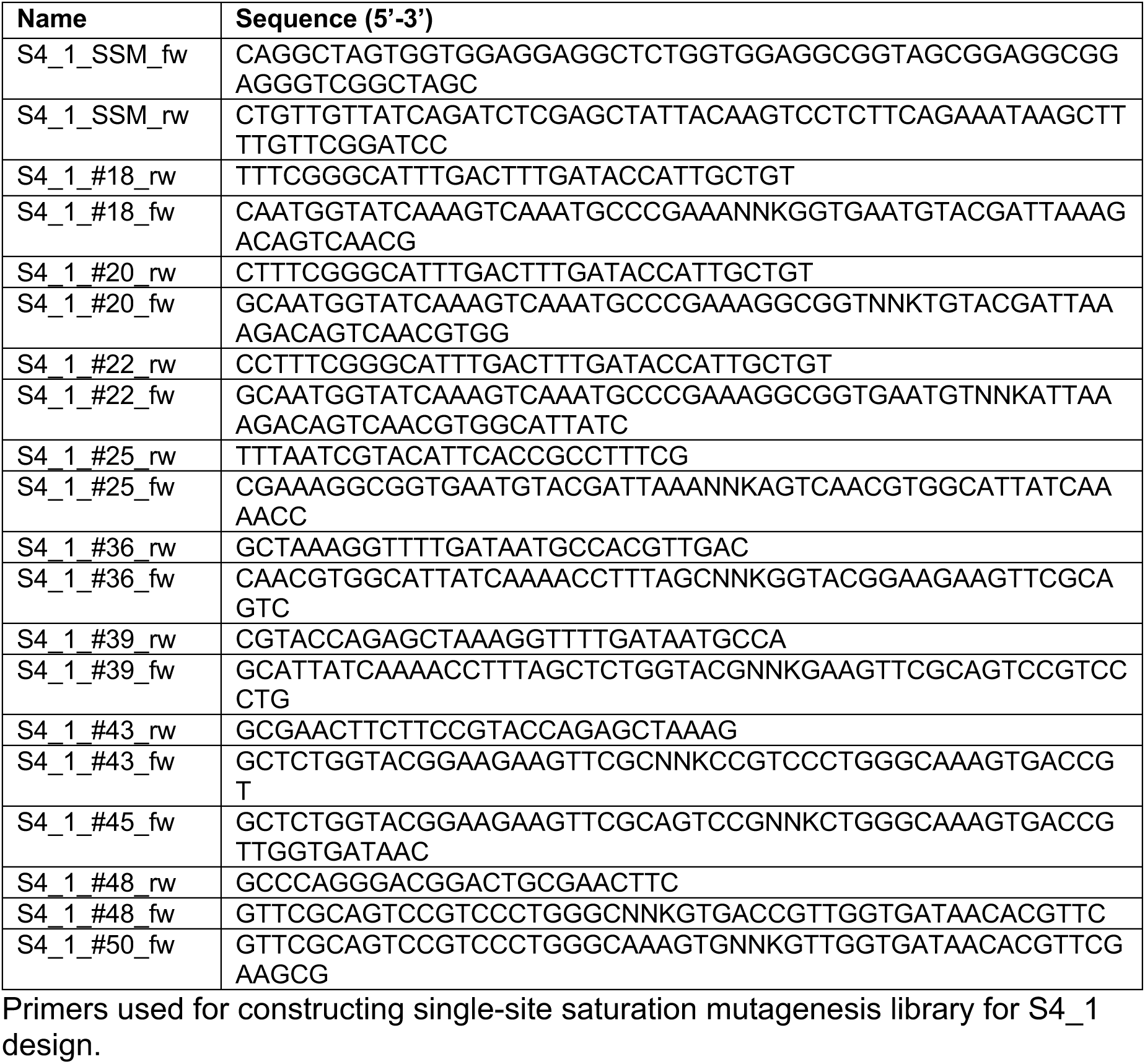

**Table S2.**
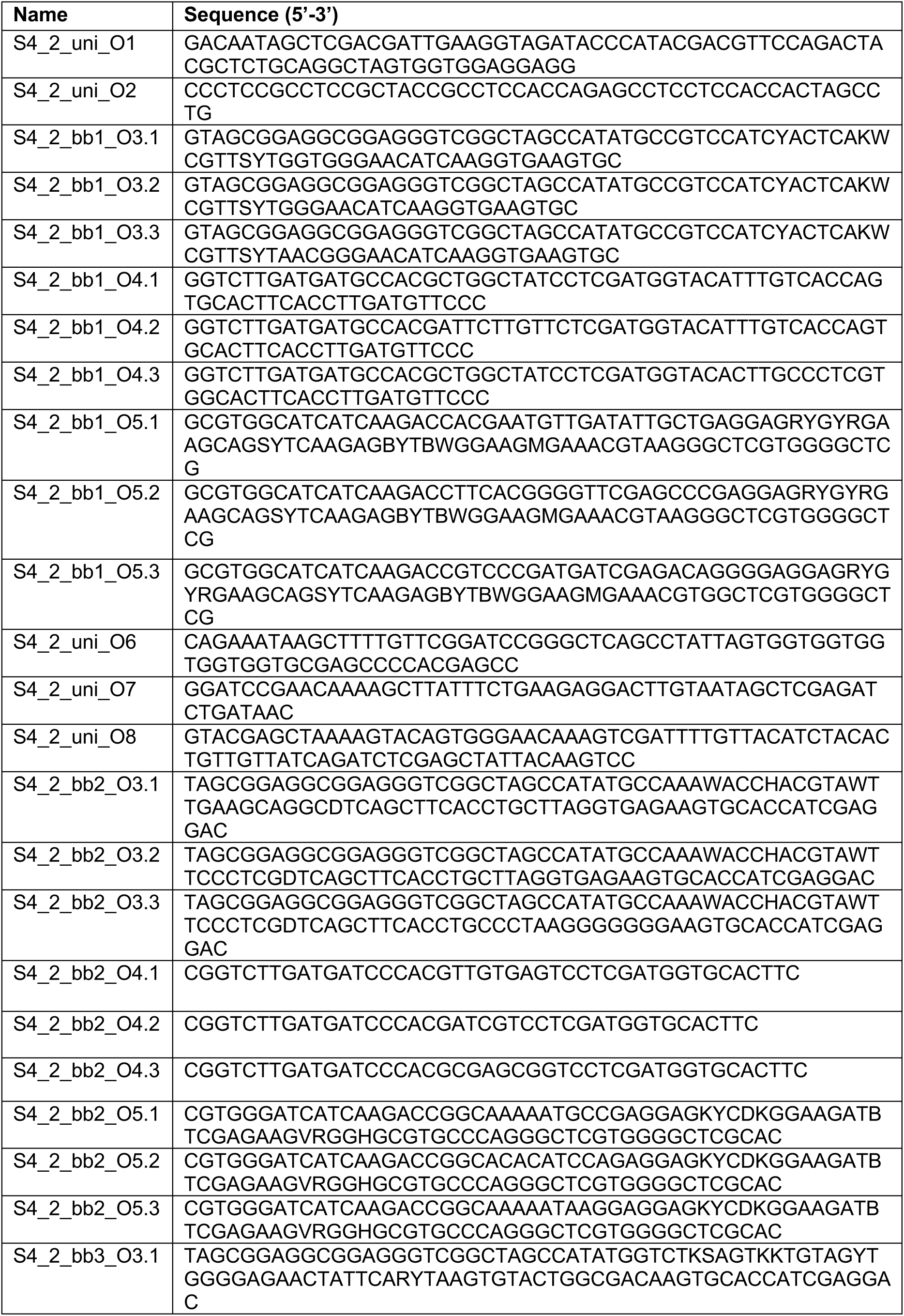
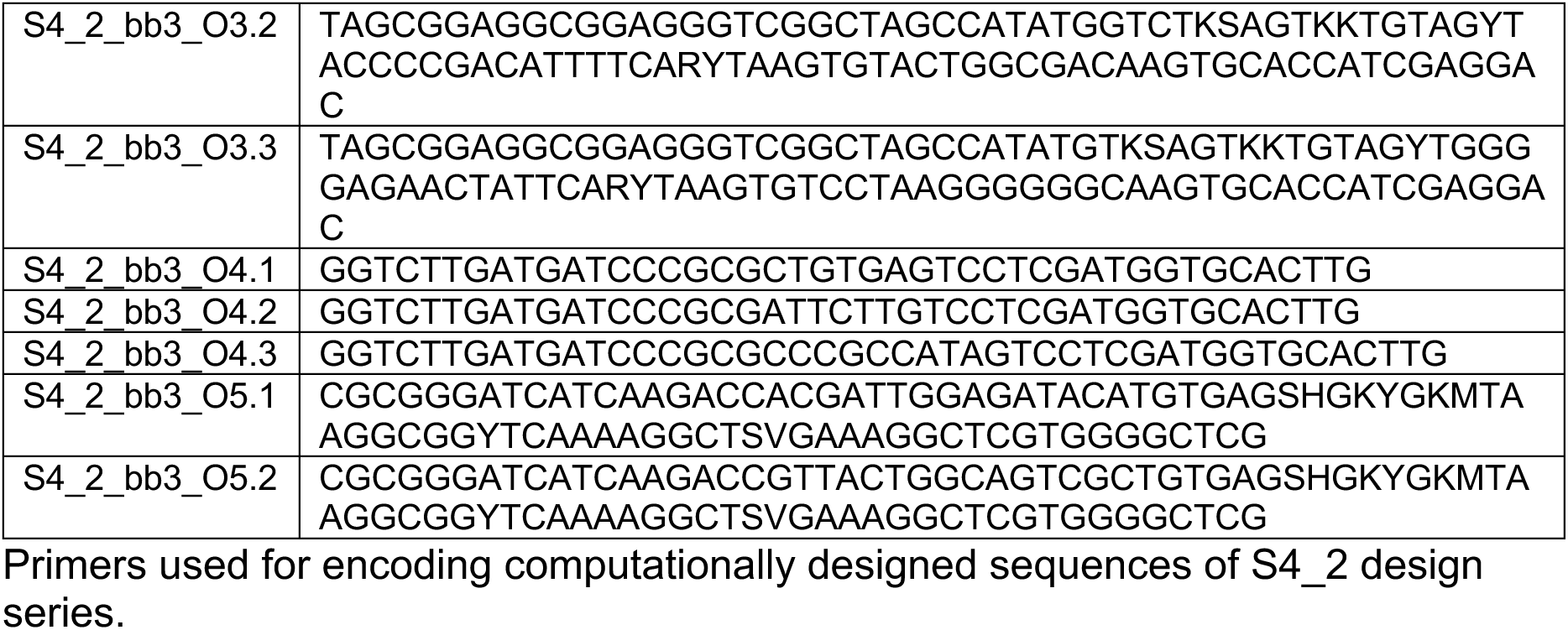

**Table S3.**
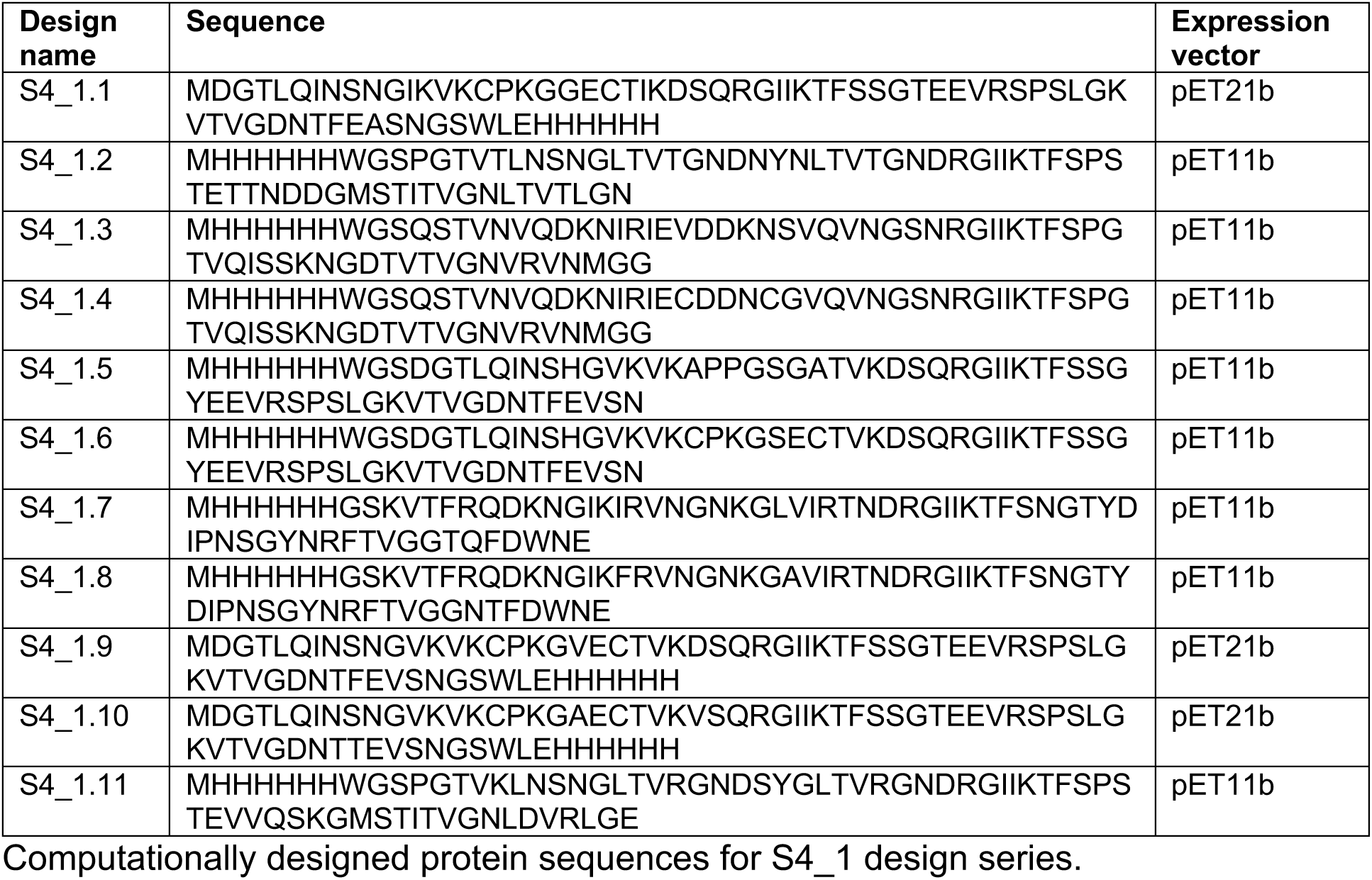

**Table S4.**
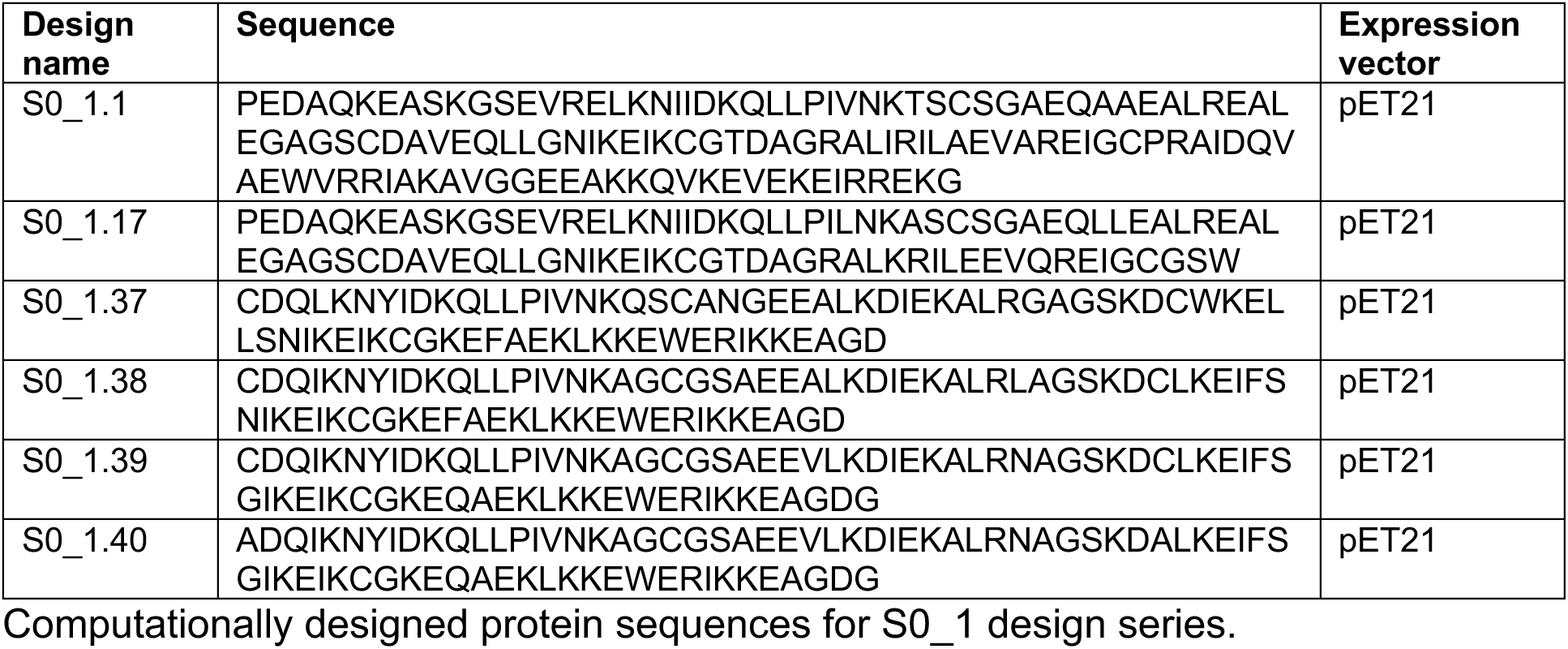

**Table S5.**
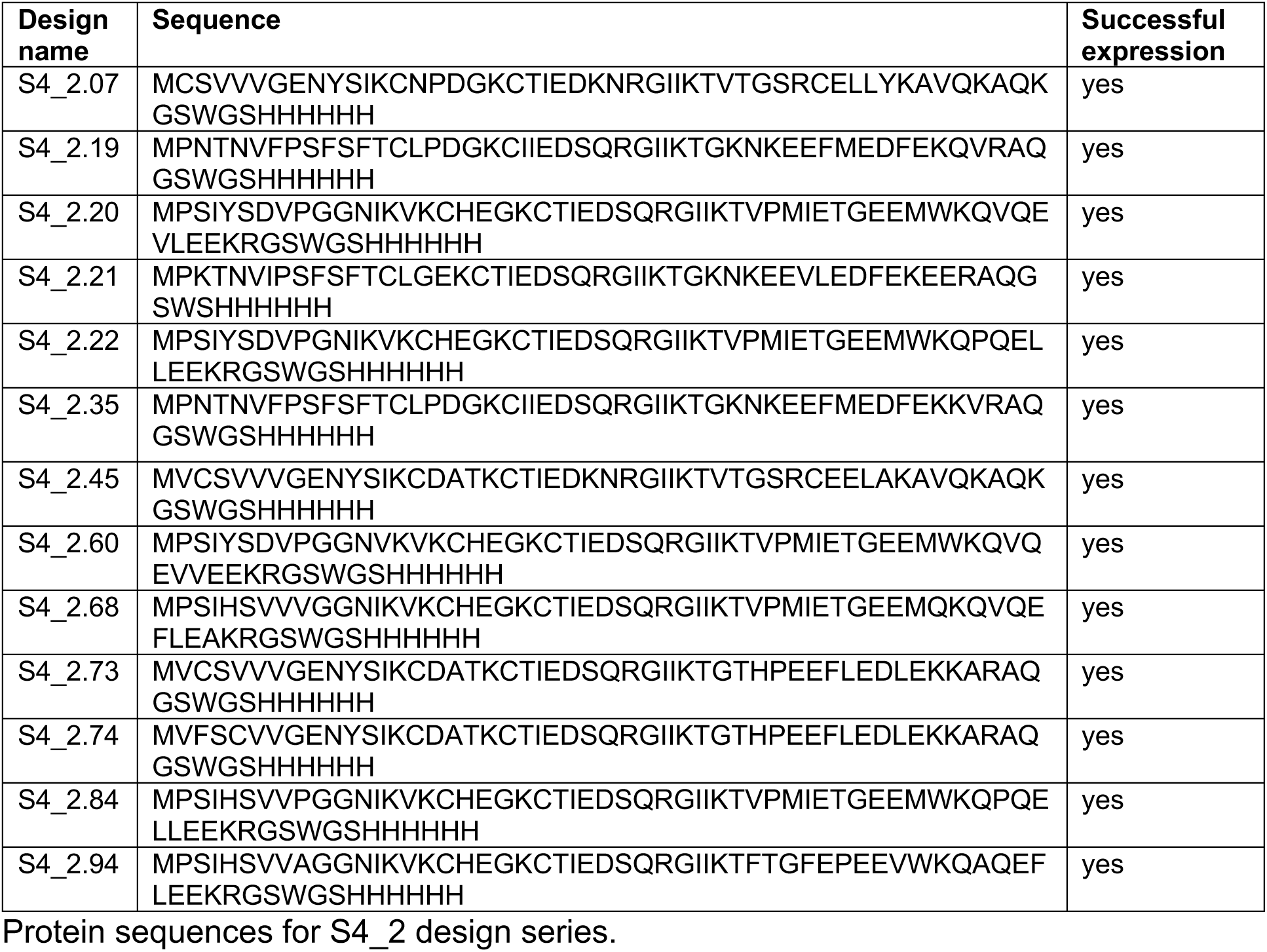

**Table S6.**
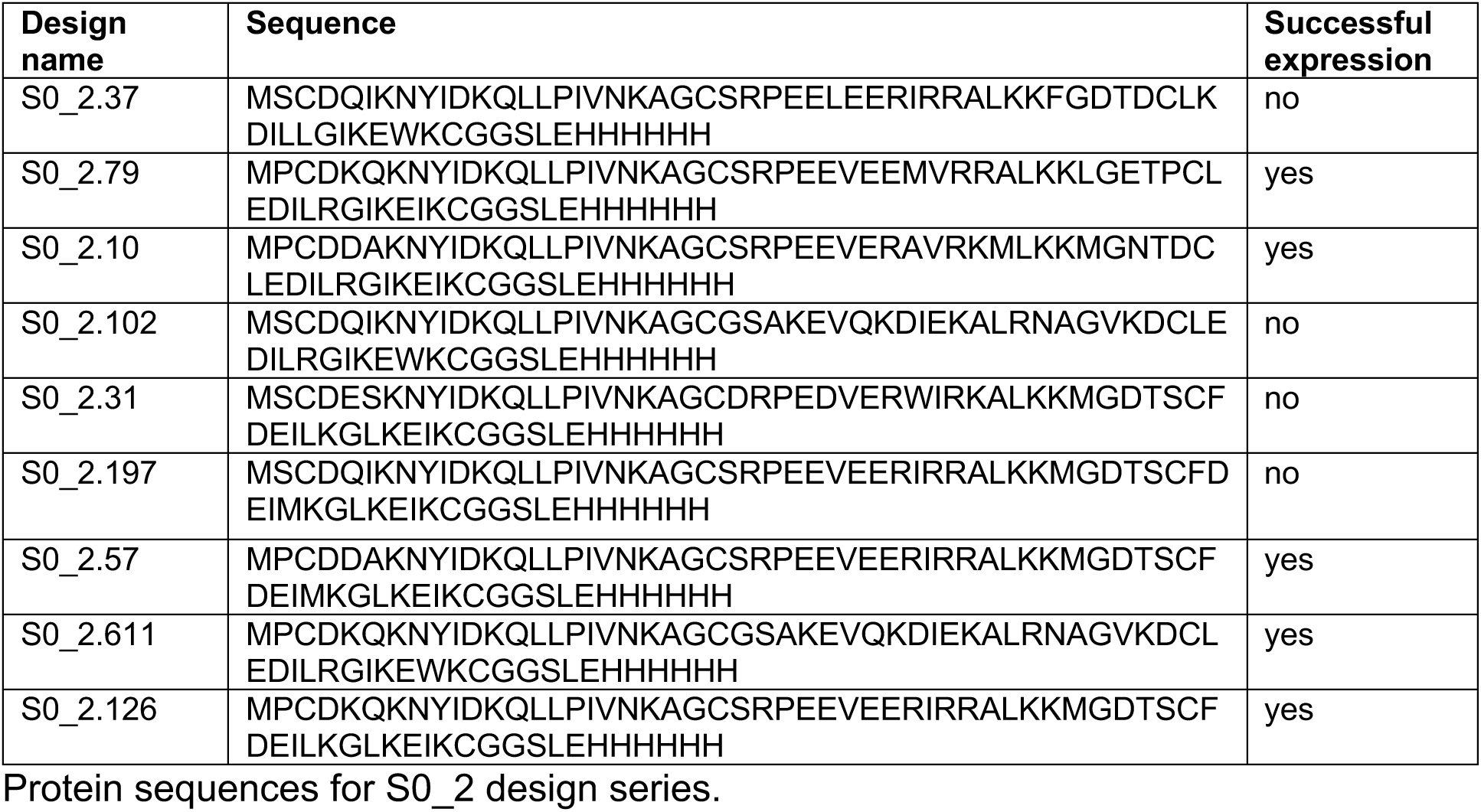

**Table S7.**
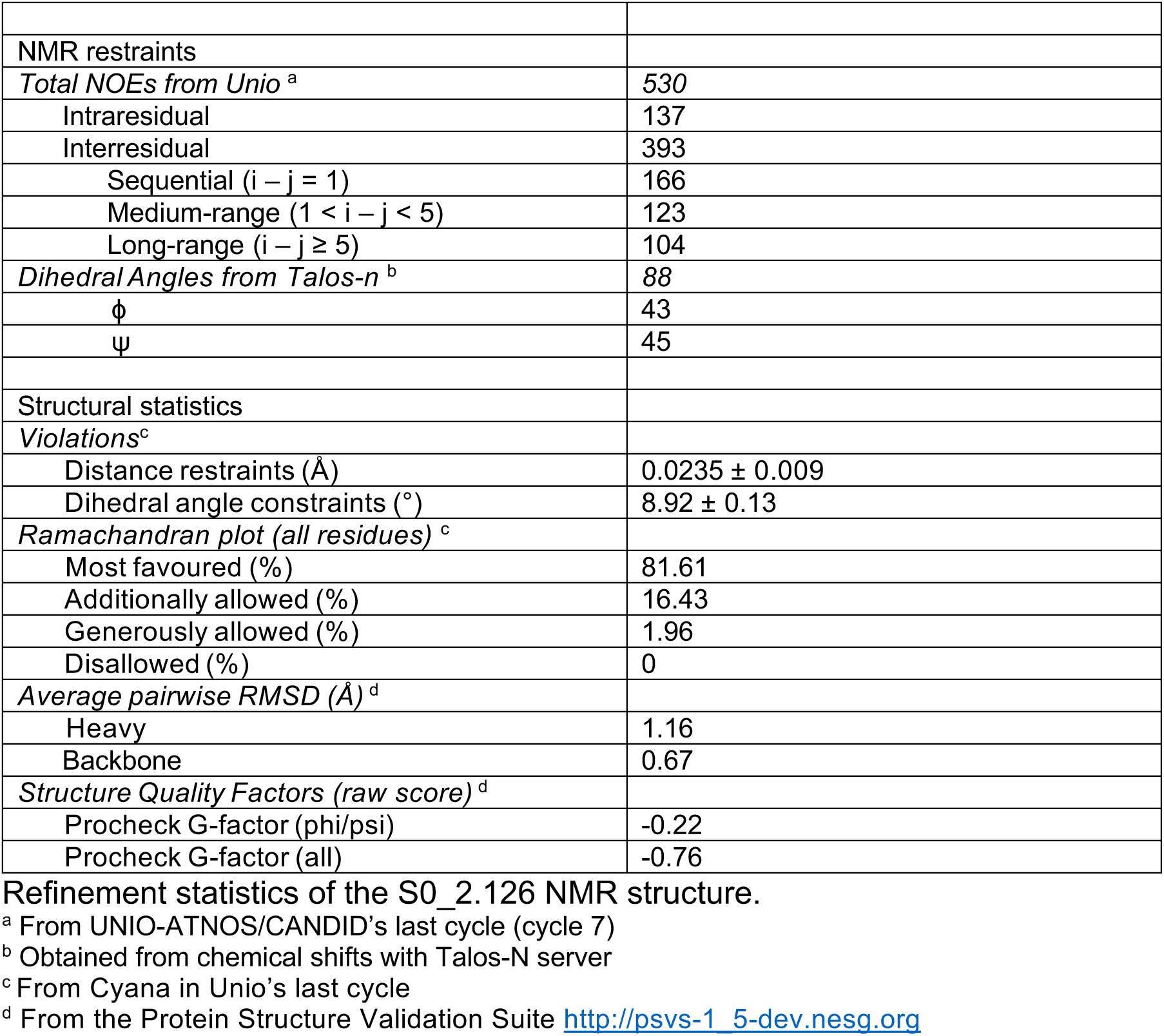

**Table S8.**
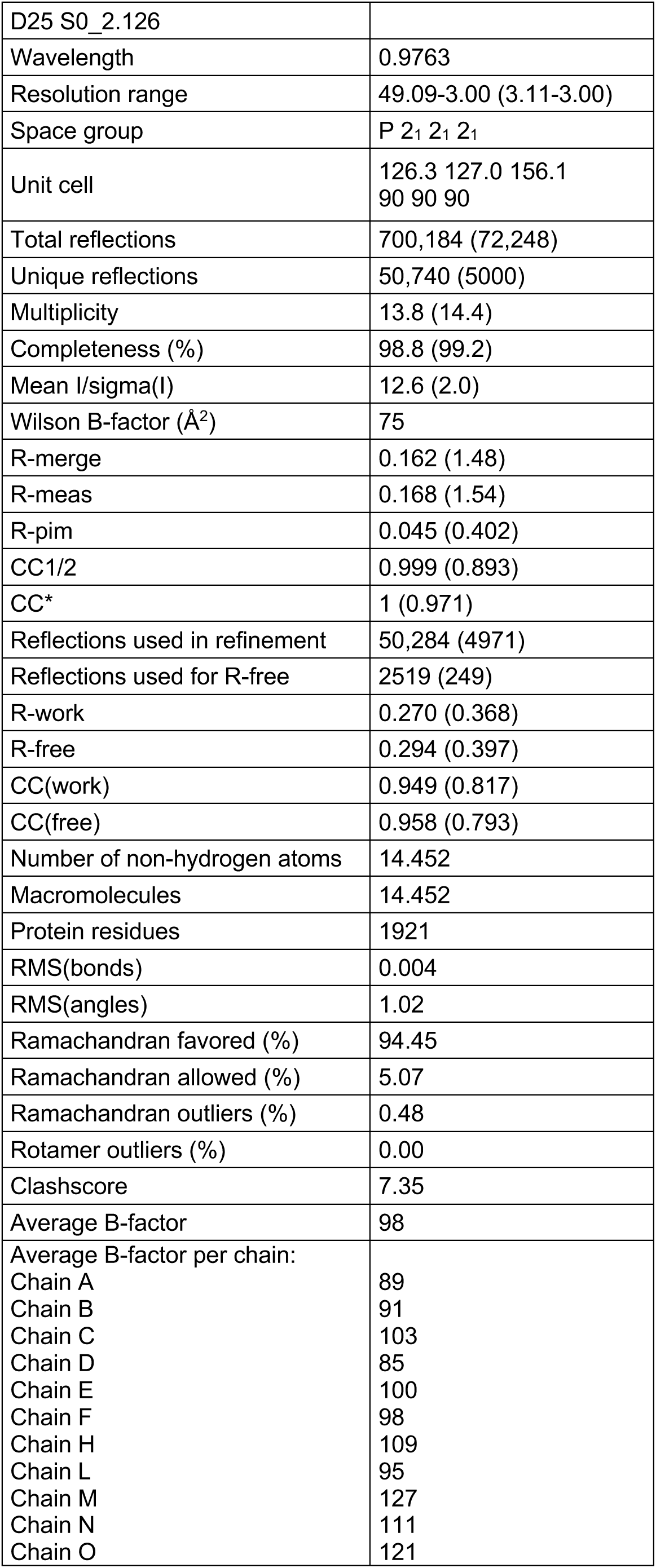
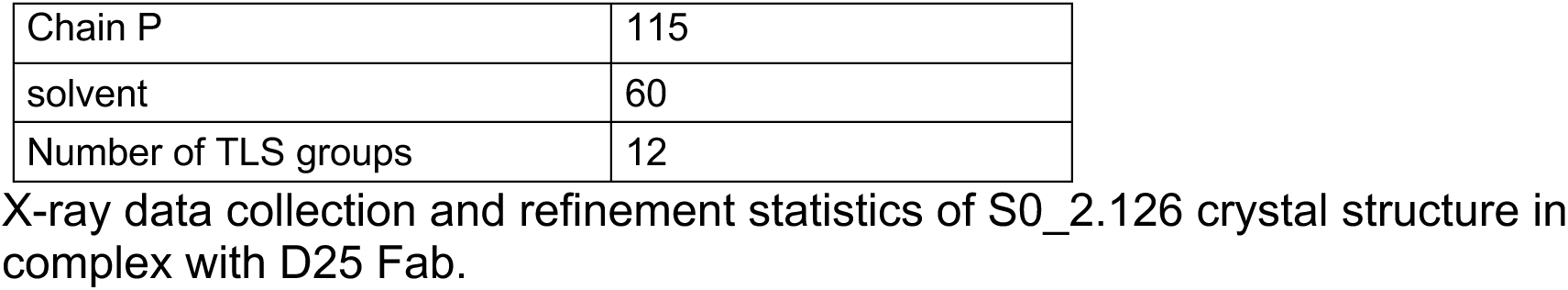

**Table S9.**
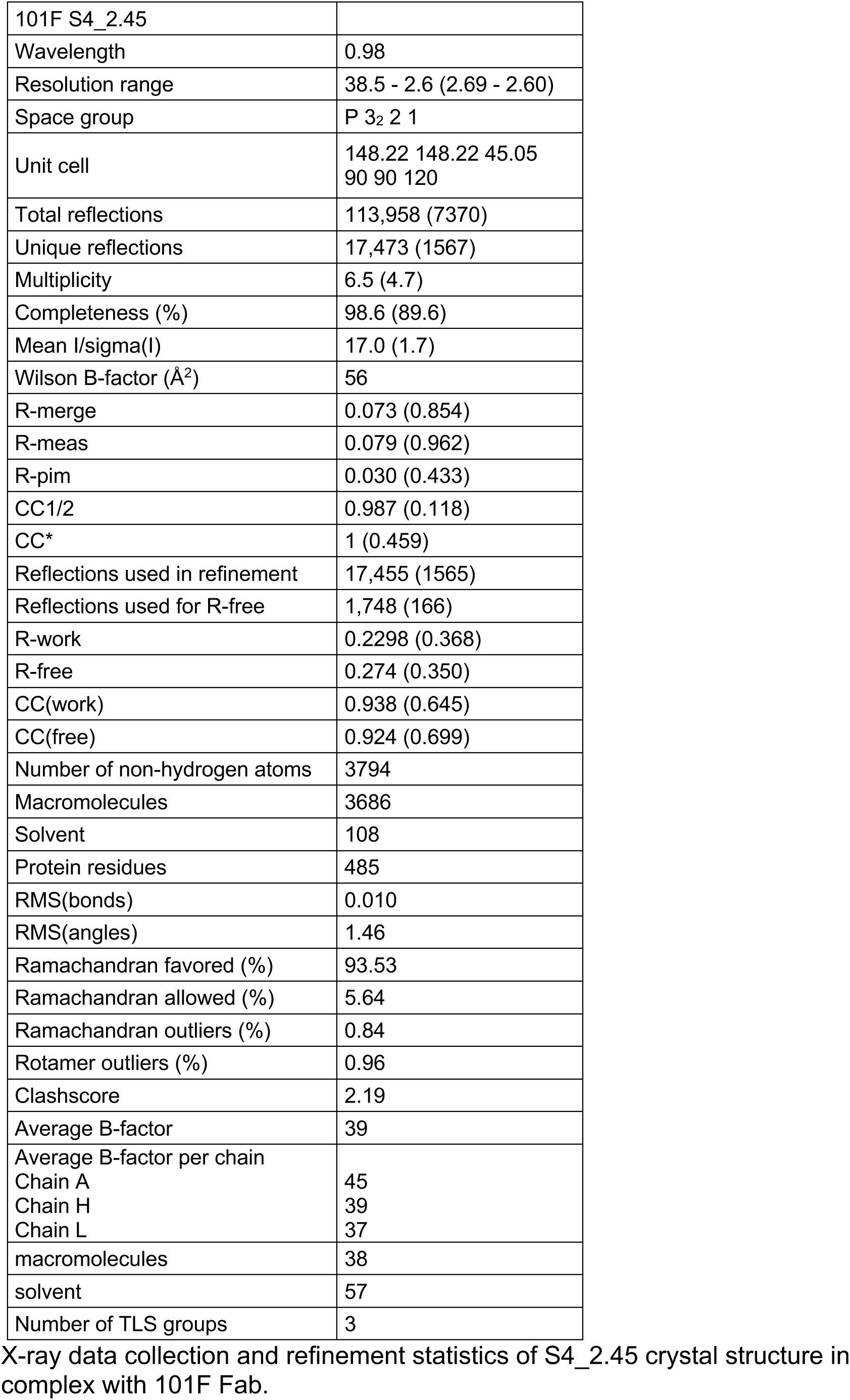

**Table S10.**
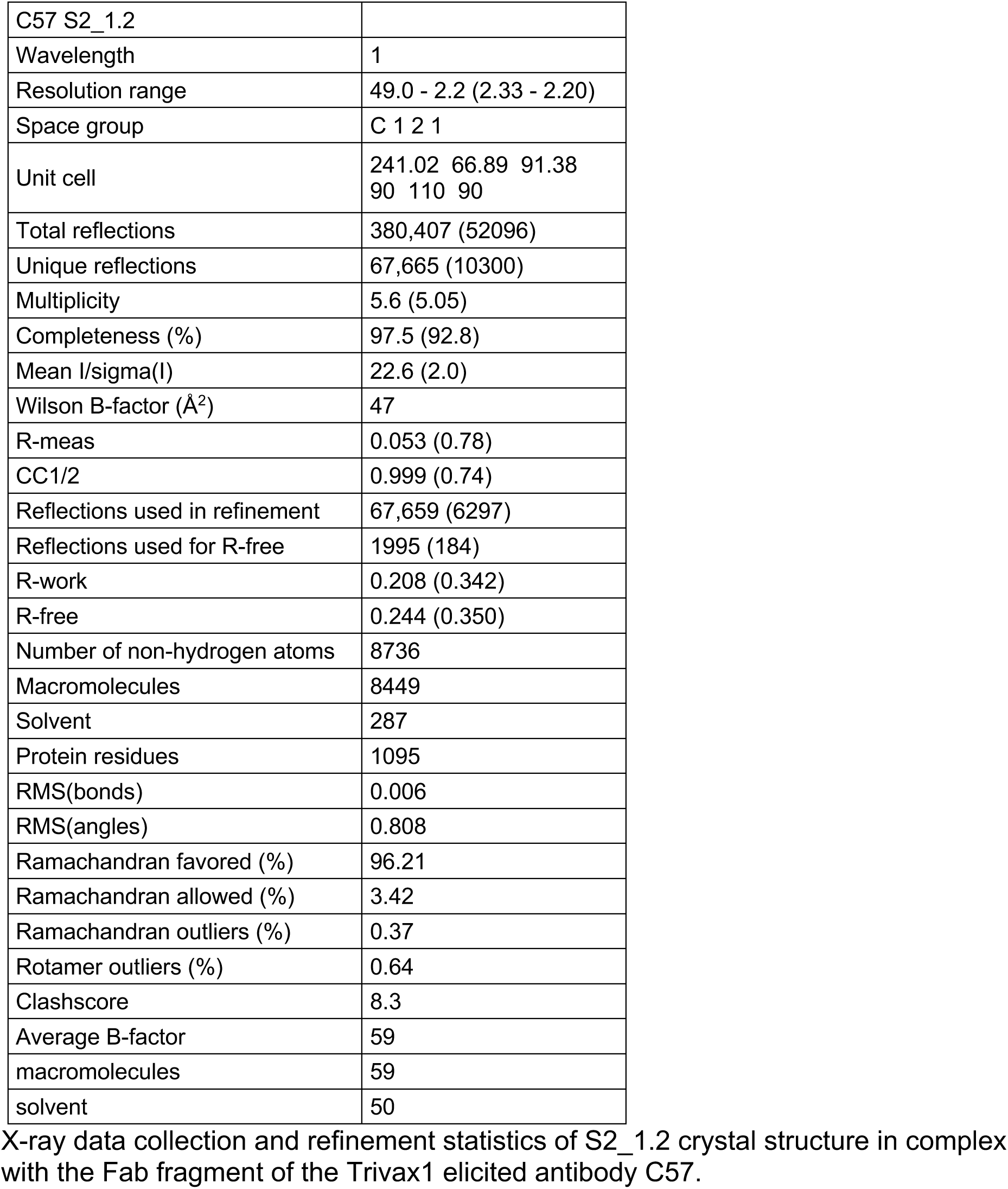

